# Emergence of specific binding and catalysis from a designed generalist binding protein

**DOI:** 10.1101/2025.01.30.635804

**Authors:** Yuda Chen, Sagar Bhattacharya, Lena Bergmann, Galen J. Correy, Sophia K. Tan, Kaipeng Hou, Justin Biel, Lei Lu, Ian Bakanas, Alexander N. Volkov, Ivan V. Korendovych, Nicholas F. Polizzi, James S. Fraser, William F. DeGrado

**Affiliations:** Department of Pharmaceutical Chemistry & Cardiovascular Research Institute, University of California, San Francisco, CA 94158, USA; Department of Bioengineering and Therapeutic Sciences, University of California, San Francisco, CA 94158, USA; VIB Centre for Structural Biology, Vlaams Instituut voor Biotechnologie (VIB), Brussels, Belgium; Jean Jeener NMR Centre, Vrije Universiteit Brussel (VUB), Brussels, Belgium; Department of Chemistry & Biochemistry, Baylor University, Waco, TX 76798, USA; Department of Cancer Biology, Dana-Farber Cancer Institute, Boston, MA 02215, USA; Department of Biological Chemistry and Molecular Pharmacology, Harvard Medical School, Boston, MA 02215, USA

## Abstract

The evolution of binding and catalysis played a central role in the emergence of life. While natural proteins have finely tuned affinities for their primary ligands, they also bind weakly and promiscuously to other molecules, which serve as starting points for stepwise, incremental evolution of entirely new specificities. Thus, modern proteins emerged from the joint exploration of sequence and structural space. The ability of natural proteins to bind small molecule fragments in well-defined geometries has been widely evaluated using methods including crystallographic fragment screening. However, this approach had not been applied to *de novo* proteins. Here, we apply this method to explore the binding specificity of a *de novo* small molecule-binding protein ABLE. As in Nature, we found ABLE was capable of forming weak complexes, which were excellent starting points for designing entirely new functions, including a binder of a turn-on fluorophore and a highly efficient Kemp eliminase enzyme (*k*_cat_/*K*_M_ = 2,200,000 M^-1^s^-1^) approaching the diffusion limit. This work illustrates how simultaneous consideration of both sequence and chemical structure diversity can guide the emergence of new function in designed proteins.

## Main Text

Proteins form the essential machinery of life. Understanding their evolution has enriched our understanding of the emergence of life and has practical implications for the design and engineering of practical catalysts and pharmaceuticals. The evolution of proteins is believed to have occurred through the self-association of short peptides to form intermolecularly folded assemblies with protein-like tertiary structures^1,2^. When linked together, these assemblies gave rise to proteins, in which individual peptide sequences could mutate independently to create asymmetric, functional proteins^3,4^ Once functional proteins emerged, they could evolve new functions and specificities through gene duplication followed by mutation of one of the gene copies^5^. Along these lines, Tawfik demonstrated a nuanced mechanism underlying the ongoing evolution of new functions in proteins. Highly evolved proteins are generally specific for their substrates, yet their active sites often display a degree of “promiscuity”, binding various molecules with specific geometries and functions, but with lower affinity than for the native ligand^6,7^. Such weak interactions can serve as starting points for the emergence of new functions by accrual of mutations on an evolutionary time scale^8^ (Fig. 1a). In this mechanism, a “specialist” protein for binding a biologically important ligand can also serve as a “generalist” for binding weakly to a diverse set of molecules. Given a positive genetic selection, the generalist activity can serve as a starting point for the evolution of a new specialist protein with increased affinity or catalytic activity for one member of the initial set of compounds that was initially bound with only low-affinity.

**Fig. 1.**
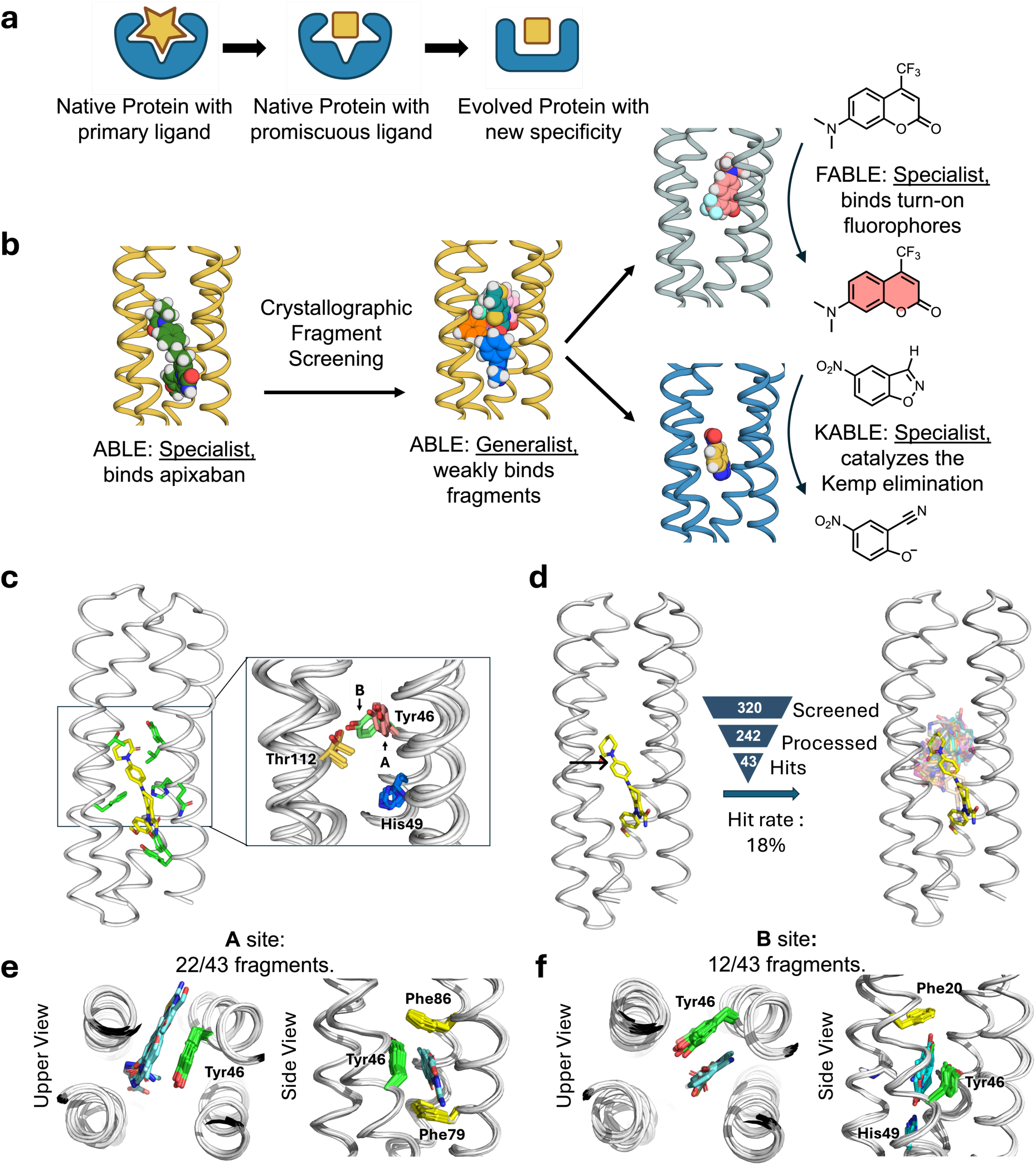
Nature-inspired evolution of *de novo* proteins using unveiled binding sites and conformations. **a**, A natural protein that has specialized for binding or catalysis of one molecule (star) can also exhibit promiscuous and weak binding to a second molecule (square). The protein can alter the shape of its binding site to match the new ligand by shifts in conformation as well as changes to the residues lining the binding pocket. **b**, Overview of the present work: we use crystallographic fragment screening to probe the ability of a *de novo* apixaban-binding protein, ABLE, to bind promiscuously with low affinity to a variety of small molecules. Based on this information, we then redesign the protein to bind turn-on fluorophores and to catalyze the Kemp Elimination reaction. **c**, Multiple conformations observed at active site residues of ABLE. The apixaban-bound ABLE structure (PDB: 6W70) is shown on the left with its binding site residues shown in green sticks. The inset shows the multiple conformations of Tyr46, His49, and Thr112 from apixaban-bound ABLE (PDB: 6W70) and the ligand-free forms (PDBs: 6W6X and 6X8N). These crystal structures have multiple monomers in their asymmetric units, each of which was superimposed in the inset. **d**, Crystallographic fragment screening reveals the chemical space of weak binders. The crystal structures of ABLE in complex with 43 fragments were each solved to between 1.3-1.6 Å resolution, using Pan-Dataset Density Analysis^35^. The structures of the complexes superposed on the ABLE complex (PDB: 6W70, chain A) are shown with the carbon atoms of apixaban in yellow sticks, and the remaining fragments’ carbon atoms in different colors. Hit rate was calculated by dividing the number of fragment hits by the number of processed datasets. **e-f**, Most fragments interact with ABLE at one of two binding sites, designated **A** and **B**, which differ by a conformational shift of Tyr46. Two views of the complexes are shown looking down the axis of the bundle (left) and rotated by approximately 90° (right). Active sites of multiple ABLE–fragment complexes, categorized as either **A** site or **B** site, are shown with interacting residues in green sticks and fragments in blue sticks. **e,** 22/43 fragments bind ABLE in an aromatic box formed by Tyr46, Phe79, Phe86, and the backbone of one helix (site **A**). **f,** 12/43 fragments interact with ABLE at site **B**, formed by Phe20, Tyr46, and the backbone of two helices. His49 sits at the bottom of the pocket, forming hydrogen bonds with polar atoms of the fragments.

The development of *de novo* protein design has been influenced by these hypothetical mechanisms of protein evolution. The first *de novo* proteins were designed by self-assembly of helical peptides^9,10^, followed by the inclusion of loops^11^ and asymmetric diversification to create catalytic metalloproteins^12,13^. Similarly, beta-hairpins bearing a Cys-Xxx-Xxx-Cys motif have been used to design a variety of catalytic and binding proteins^14–17^. The functional elaboration of *de novo* protein scaffolds has the advantage that an arbitrary number of amino acid insertions deletions and substitutions can be simultaneously made through design in ways that are not feasible through the more incremental stepwise process of natural protein evolution^18–22^. Indeed, using computational design, *de novo* proteins have been designed for various functions^16,18–30^. However, in each case, the design process has driven towards a single, specialized function. While the specificity of the proteins towards closely related ligands^25^ or substrates^24,28–32^ have been evaluated in isolated cases, the large-scale structural examination of specificity of *de novo* proteins has not been reported. Recent advances in X-ray crystallography throughput now enable the detection of low-affinity molecules bound to receptor proteins^33^. In drug discovery, this is applied by soaking hundreds to thousands of small-molecule fragments (<300 Da) into crystals and identifying binders through changes in the electron density^34^. Real-space background subtraction enhances sensitivity, allowing detection and modeling of bound molecules, with typical hit rates of 1-20%^35^. Here, we determine the specificity of a *de novo* protein towards a common chemotypes in a diverse set of small molecules using crystallographic fragment screening and then use this information to guide the design of entirely new binding and catalytic functions.

We focused on ABLE (**a**pixaban-**b**inding helica**l** bundl**e**), a *de novo* protein that was designed to bind with high specificity to the antithrombotic drug, apixaban^36^. Inspired by Tawfik’s hypothesis^7^, we examined how ABLE, initially designed as a specialist binder of the antithrombotic drug apixaban, could give rise to new specialists: one fluorophore-binder with high specificity for a different small molecule and a catalyst whose activity exceeds that of proficient natural enzymes^37,38^ (Fig. 1b). We first explored ABLE’s promiscuity by screening a library of several hundred small-molecule fragments at millimolar concentrations using X-ray crystallography. This approach identified weak molecular interactions not anticipated in the initial designs, which provided pathways to new specificities and catalytic activity, achieved through computational design. Notably, despite these functional transformations, the core tertiary structural framework of ABLE was retained (< 1.5 Å with Cα RMSD).

The trajectory of the design of these proteins, beginning with self-associating peptides^9–11^ and ultimately leading to highly refined binders and catalysts, has many parallels with natural evolution of proteins from assemblies of peptides to highly functional proteins^11,25,26,36,39–42^. Our work is particularly relevant given the functional diversity of helical bundles, one of the most ancient and adaptable protein architectures. Most transmembrane proteins, including G-protein coupled receptors (GPCRs), rely on helical bundles for signal transduction, transport, and enzymatic functions^43^. The enzymes that catalyze some of the most important reactions for the evolution of life, such as the formation and utilization of O_2_ and the generation of ion and proton gradients, are also helical bundles^44–46^. Similarly, soluble helical bundle proteins are involved in electron transport^47^, small-molecule binding^48^, and catalysis^49^. Given their broad functional diversity, helical bundles have thus played a crucial role in the early evolution of life. By linking modern protein design efforts to fundamental principles of evolution, the present study provides insights into how novel specificities and catalytic functions can emerge from a common structural framework.

### Conformational plasticity and binding specificity of ABLE from crystallographic fragment screening

While *de novo* proteins have been designed to discriminate variants of their substrates, their ability to bind with low affinity to structurally unrelated substrates has not been explored. We therefore evaluated the promiscuity of ABLE, which was designed to bind the antithrombotic drug apixaban in an extended grove within a helical protein (Fig. 1c).

We used X-ray crystallography to map regions of ABLE’s binding site that were capable of binding to a diverse set of 320 “fragments” (MW < 300 Da) of drug-like molecules with a wide range of aliphatic, aromatic and polar functional groups^50^. Each compound (10 mM) was coincubated with ABLE crystals to form complexes in the crystal lattice. We obtained high-quality diffraction data from 242 of the compounds in the library. Of these, 43 gave diffraction data with electron density that could be ascribed to a bound fragment (Fig. 1d).

The frequency of forming solvable complexes provides a measure of the promiscuity of the site, and the structures of the resulting complexes provide a view of the diversity of interactions that can be formed within the binding site. The hit rate for discovering fragments bound to ABLE (18%) is similar to the reported hit rates of 7% and 19% for Nsp3 macrodomain of SARS-CoV-2 and Chikungunya virus, respectively, using the same fragment library and screening method^34,51^ (Fig. 1d). All but four fragments bound to ABLE within the apixaban binding site. Thus, by this criterion, ABLE has a specificity within the range seen in natural proteins.

The 39 fragments that bound in ABLE’s apixaban-binding site define a set of chemotypes that favorably interact with ABLE (Extended Data Fig. 1 and 2). The bound fragments have a statistically significant tendency to be slightly larger, more apolar, and richer in aromatic rings when compared to the overall library (Supplementary Fig. 1). The preferred interacting groups found in the complexes show some differences from the interactions that are important for binding the drug apixaban. For example, while apixaban has a primary carboxamide (–CO–NH_2_) that is bound near the surface of ABLE in the drug complex, this polar group was not seen in the bound fragments, all of which bound more deeply into the pocket.

The protein’s structure was nearly identical in all the complexes (sub-Å Cα RMSD). About two-thirds of the fragments bound to a conformer found in the apixaban complex (conformer **A**), that features an aromatic box comprised of the sidechains of Tyr46, Phe79, Phe86, and the backbone of **helix 3** (Fig. 1e). However, a considerable fraction (12/43) of the fragments bound to a second conformer **B**, which highlights a distinct aromatic box comprised of Tyr46 in an alternate sidechain conformation (Fig. 1f). Although there is a relatively modest difference in Tyr46’s sidechain torsional angles between conformers **A** and **B**, this small change propagates to a larger displacement of its long aryl sidechain. In addition to aromatic and apolar interactions, most fragments additionally form polar interactions. For example, Thr112 often engages polar groups in conformer **A**, as does His49 in conformer **B** (Extended Data Fig. 2 and Supplementary Fig. 2). This malleability enables ABLE to adjust to the shape and properties of the fragments^52^.

Often, proteins use conformational sub-states to bind promiscuously to new substrates, and these pre-existing conformational sub-states become locked in place as the protein evolves new specificity for the novel substrate^6^. We were therefore curious whether some of the novel conformers seen in the fragment complexes, but not in the apixaban complex, might pre-exist in multiple crystal structures that we have solved of apo-ABLE. We indeed observed that the critical Tyr46 residue adopted both the **A** and **B** conformers in different structures of apo-ABLE (Fig. 1c). A similar pair of conformations emerged from multiple molecular dynamics simulations of the protein in the drug-free state (Extended Data Fig. 3). Other sources of conformational variability in this region included the sidechains of His49, which also populate more than one rotamer (Supplementary Fig. 3). Collectively, this analysis indicates that key elements of the alternate conformer **B** are already found in the uncomplexed protein, indicating that ABLE does not pay a large reorganization penalty to adopt conformer **B**.

### Designing proteins to bind fluorogenic ligands

We next asked whether the newly discovered site defined by conformer **B** would be a good starting point to design specialized proteins with altered binding specificity. Site **B** bound the fragment 7-hydroxycoumarin (Fig. 2a), which represents the core of a large class of fluorophores. We therefore asked whether we might use the coordinates of the ABLE complexes to bind the larger, brighter and environment-sensitive, turn-on fluorophore Cou485 (1, 7-(dimethylamino)-4- (trifluormethyl)-coumarin, Fig. 3a)^53^. Procedurally, this task involved expanding the top of the **B** site to accommodate the additional trifluoromethyl and dimethylamino groups while filling the lower, unoccupied section of the pocket with well-packed hydrophobic amino acids (Fig. 2a). Using a previously described protocol ^54,55^, we selected 43 of ABLE’s 126 residues to vary during redesign. Fig. 2a illustrates the gradual migration of the binding site, which is accompanied by binding to the larger ligand. OmegaFold ^56^ was used to determine whether the new designs would be preorganized in a conformation capable of binding Cou485; other selection metrics are described in the Methods.

**Fig. 2.**
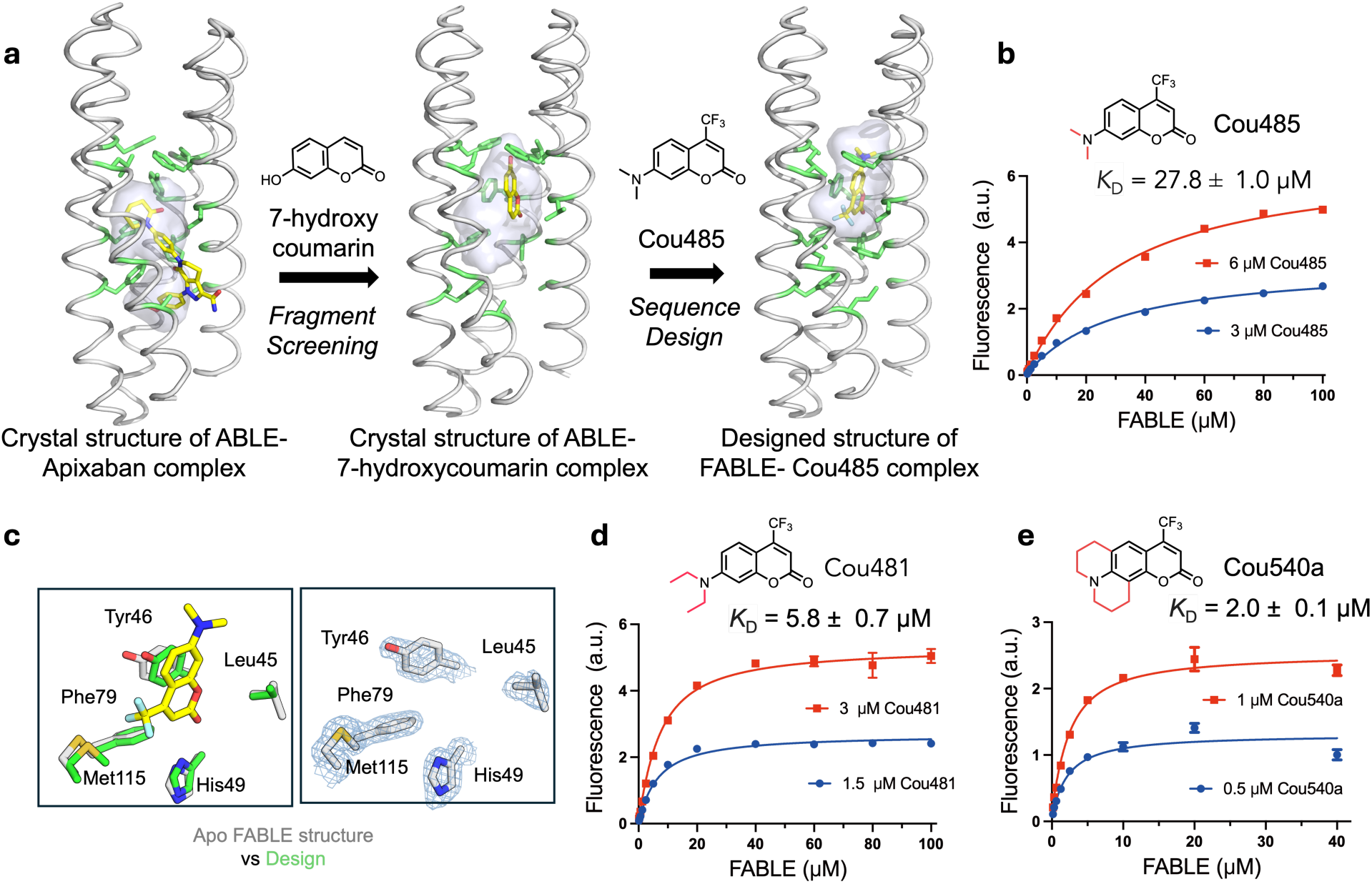
Design of a fluorescent ABLE (FABLE) binding to fluorogenic coumarins. **a,** 7-hydroxycoumarin was found from crystallographic fragment screening to bind ABLE. The **B** box (Fig. 1f) was used to design five sequences for binding to Cou485. The positions of the apixaban-interacting residues (green sticks) and the binding cavity (grey surface) show how the size and location of a binding site change as the site transitions from binding apixaban, to a small coumarin, and then to a Cou485. **b**, A titration of FABLE into Cou485 shows a single-site binding isotherm. The dissociation constant was obtained by globally fitting a single-site binding model to data obtained by titrating FABLE into Cou485 at two fixed fluorophore concentrations, 3 and 6 μM. **c**, Left: Overlay of the designed FABLE-cou485 complex with the uncomplexed FABLE crystal structures (PDB: 9DWC, comparison with 9DWA and 9DWB are in Supplementary Fig.8). Right: 2mF_O_-DF_C_ electron density (1 σ) contoured around the active site residues of FABLE (PDB: 9DWC). **d-e**, The binding constants for FABLE’s interaction with Cou481 **(d**) and Cou540a **(e)** were obtained at the indicated fixed fluorophore concentrations and variable FABLE concentrations as in **b**. These two were identified after screening a set of 37 fluorophores, including coumarin derivatives and other fluorophore scaffolds. Error bars in **b**, **d**, and **e** represent standard deviations of three independent measurements.

**Fig. 3.**
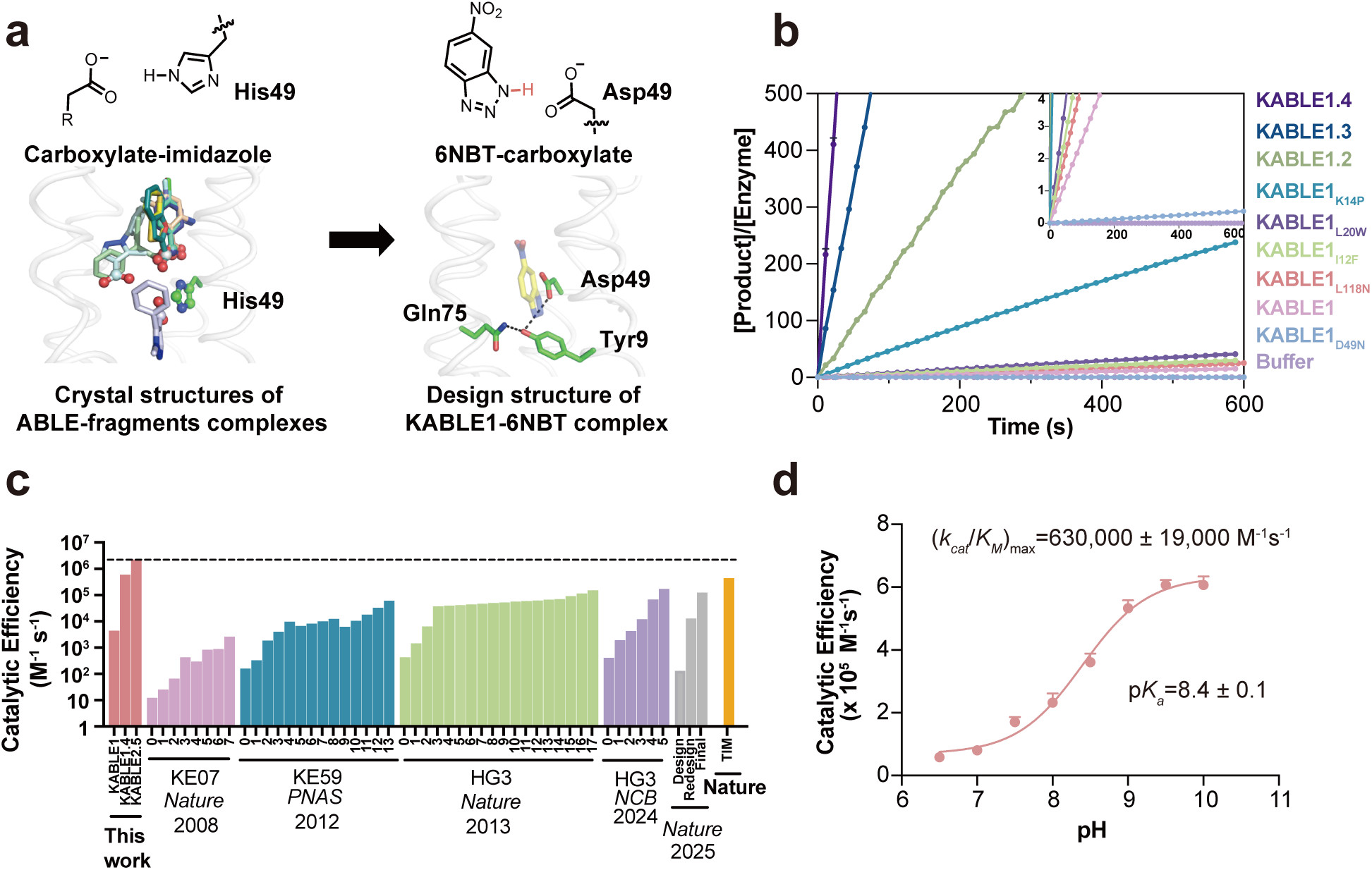
Design of an efficient Kemp eliminase ABLE (KABLE) and its directed evolution. **a**, Fragment-inspired design of KABLE. This carboxylate-to-imidazole interaction is analogous to the carboxylate-to-isoxazole interaction at the active site of a kemp eliminase, suggesting that an active catalyst might be designed by changing His49 into an Asp. Left: Nine of the fragments contain carboxylates that bind to His49. Right: The active site was introduced into the ABLE backbone by substituting the His49 sidechain with the most frequently observed rotamer of Asp^107^. 6NBT (sticks with yellow carbons) was positioned in the saddle point geometry of the transition state for the carboxylate-catalyzed reaction as determined by quantum mechanical calculations^60^. Sequence design was next carried out with LigandMPNN^76^ and Rosetta FastRelax^55^. The resulting Asp triad (Asp49-Tyr9-Gln75) is shown in sticks. **b,** Catalytic activity for the Kemp elimination reaction for the original sequence, designated KABLE1. Mutants arising from combining variants from site-saturation mutagenesis show a steady progression as more sites are included. The data are expressed as the number of turnovers (Y-axis) per unit time (X-axis). The mutations on KABLE1 are expressed as a subscript. KABLE1.2 combines Lys14Pro and Leu20Trp substitutions; KABLE1.3 combines Ile12Phe, Lys14Pro, and Leu20Trp. KABLE1.4 combines Ile12Phe, Lys14Pro, Leu20Trp, and Leu118Asn. The reaction was measured at pH 8 with an initial substrate concentration of 240 μM **c**, Catalytic efficiency of designed and evolved Kemp eliminases during experimental optimization, in comparison with TIM, a natural enzyme with a similar mechanism. For each protein, each step involved in the iterative improvement is indicated. The first on the left reflects the activity of the initial design; the subsequent bars reflect the activities after saturation mutagenesis (KABLE, this work), combining favorable mutants after multiple rounds of directed evolution (KE07, KE59, HG3)^60–63,108^ or computational redesign^71^. TIM refers to triosphosphate isomerase^38^. **d,** pH-dependence of *k*_cat_/*K*_M_ for KABLE1.4. *k*_cat_/*K*_M_ obtained from individual pH was used to fit the (*k*_cat_/*K*_M_)_max_ and p*K*_a_. Error bars in **d** represent the standard errors of the mean from three independent measurements.

Five of the designed proteins with sequence pairwise identity ranging from 77% to 81% were selected for experimental characterization (Supplementary Fig. 4). All five were helical, thermostable to 90 °C and bound Cou485 (Extended Data Fig. 4). The intensity of the emission spectra of Cou485 increases markedly and shifts towards lower wavelengths when bound in a rigid, hydrophobic binding site^57^. Similarly, the fluorescence of Cou485 (6 μM) increased by 10 to 99-fold in the presence of 40 μM of the five different proteins (Supplementary Fig. 5). Global analysis of the titration curves obtained at two fixed Cou485 concentrations and variable protein concentrations indicated that all five proteins bound Cou485 with *K*_D_ < 100 μM, and three exhibited *K*_D_ ranging from 28 μM to 38 μM (Extended Data Fig. 4). The protein showing the most potent *K*_D_ (27.8 ±1.0 μM) was designated as FABLE (Fluorescent ABLE) (Fig. 2b). Attesting to their specificity, the starting ABLE protein showed non-saturable curve, indicative of very weak binding (Supplementary Fig. 5 and 6).

Ligand efficiency is an empirical metric used to assess the effectiveness of a protein’s interaction with a small molecule ligand ^58,59^. Large ligands have more opportunities to interact favorably, so ligand efficiency roughly normalizes for the size of the molecule by dividing the free energy of binding (1 M standard state) by the number of heavy atoms in the ligand. Most FDA-approved small molecule drugs have ligand efficiency around 0.3 kcal/(mol • heavy atom count)^59^. By comparison, ligand efficiencies for the five proteins with Cou485 range from 0.31 to 0.35 kcal/(mol • heavy atom count) (Supplementary Table 1). To address the question of whether FABLE had a well-defined, preorganized structure, we determined crystallographic structures of ligand-free FABLE in three different crystal forms (1.5-1.9 Å resolution). Each of the structures was in excellent agreement (< 1 Å Cα RMSD) with the designed model and the AlphaFold3-predicted structure. (Fig. 2c, Supplementary Figs. 7 and 8, and Supplementary Table 2).

### Experimental exploration of chemical space to probe the specificity of FABLE

Having explored sequence space to create a specialist that is specific for Cou485, we next re-examined chemical space to evaluate FABLE’s specificity for alternative fluorophores. A search of 37 fluorophores that shared molecular features such as hydrophobicity and aromaticity (Supplementary Table 3) identified only two compounds that showed a large (> 4-fold) increase in fluorescence intensity in the presence of FABLE vs. the starting ABLE protein (Extended Data Fig. 5). These compounds, Cou481 and Cou540, were derivatives of cou485 with slightly larger apolar groups at the 7-amino position of the coumarin core, which bound FABLE with *K*_D =_ 5.8 ± 0.7 µM and 2.0 ± 0.1 µM (Figs. 2d and 2e, Supplementary Fig. 9), and ligand efficiencies of 0.35 and 0.34, respectively (Supplementary Table 1). The maximal fluorescence enhancement at saturating concentrations of FABLE was 48-fold for Cou485, 123-fold for Cou481, and 117-fold for Cou540 (Supplementary Fig. 8). Thus, the generalist binding activity of ABLE provided an excellent starting place for a specialist that bound a chemically distinct compound.

### Design of an efficient Kemp eliminase

We next asked whether the generalist binding activity of ABLE could be used to engineer enzymatic function. We noted that many of the fragments that bound to the aromatic boxes were similar in shape and properties to the substrate used in the Kemp elimination, a reaction that has been benchmarked over several decades in the quest to design enzyme-like proteins^60–69^. This reaction involves the removal of a proton from a C–H bond of the substrate (5-nitrobenzisoxazole, 5NBI; Fig. 3a), similar to the carboxylate-mediated abstraction of a proton from a CH bond in the substrates of triosephosphate isomerase and ketosteroid isomerase^38,70^. The success rates and catalytic efficiency of computationally designed Kemp eliminases have been relatively low prior to rational redesign or directed evolution (0 – 200 M^-1^s^-1^)^60,63,64,71–73^. Previous computationally designed Kemp eliminases were based on the redesign of the active sites of natural enzymes. We were curious whether a greater rate might be achieved with a *de novo* scaffold and our knowledge of ABLE’s potential binding interactions.

To design a series of Kemp eliminases based on ABLE (KABLE proteins), we targeted a transition state analogue, 6-nitrobenzotriazole (6NBT, Fig. 3a). We computationally searched^63,64^ for backbone positions of ABLE, in which a Glu or Asp in a low-energy rotamer could position 6NBT in a catalytically competent orientation in the pocket delineated in fragment complexes. Leu108Glu was one site chosen because a low-energy rotamer of Glu108 placed the transition state analog inside the aromatic box of the **A** conformer. We also evaluated His49 as an attractive position for substitution of an Asp residue because histidine has a similar size, shape, and polarity as aspartic acid. Importantly, His49 is located near the bottom of the binding site, where it is a hotspot for interacting with carboxylate-containing fragments (Fig. 3a and Supplementary Fig. 10). Modeling an Asp sidechain at residue 49 reverses the interactions and allows a strong hydrogen bond with 6NBT and the Asp carboxylate’s more basic (syn) lone pairs of electrons^74,75^ (Fig. 3a). With these restraints in place, we used ligandMPNN^76^ and Rosetta FastRelax^55^ to optimize the sequence to bind 6NBT while stabilizing the conformation of either Glu108 or Asp49. To increase their basicity, we introduced no more than one H-bond donor to the Asp49 or Glu108 carboxylates, leaving three potential H-bond accepting sites unsatisfied^77^. Computed models were filtered based on the RMSD between the design model and that predicted for the unliganded protein computed by RaptorX^78^ and ESM2^79^ (to assess preorganization of the substrate-free protein), as well as their pLDDT scores (Supplementary Fig. 11) that assess the likelihood of the predicted structures.

We selected five proteins from each strategy for experimental characterization, with pairwise sequence identity ranging from 26% to 71% compared to ABLE (Supplementary Fig. 4). All of the ten proteins expressed well in soluble form (Supplementary Fig. 12). One of the sequences (designated KABLE0) from the first strategy had modest catalytic activity (*k*_cat_/*K*_M_ = 8 M^-1^s^-1^ at pH 8, Supplementary Fig. 13). A second protein (designated KABLE1) with His49Asp as the base exhibited much greater activity. The pH-rate profile of KABLE1 had a sigmoidal shape typical of a single-site protonation (Supplementary Fig. 14) with a maximal value of *k*_cat_/*K*_M_ of 3800 ± 1400 M^-1^s^-1^ (Fig. 3b and c, Extended Data Tables 1 and 2, Supplementary Table 4). The midpoint of the profile was 9.3 **±** 0.2, much higher than the intrinsic p*K*_a_ of Asp (Extended Data Table 2, Supplementary Fig. 14). Mutation of Asp49 to Asn eliminated the catalytic activity, indicating that this residue was likely the protonatable group (Fig. 3b and Supplementary Fig. 15). The elevated p*K*_a_ for Asp49 is consistent with its placement in a very apolar environment, as seen in a Kemp eliminase based on ketosteroid isomerase, which included an Asp with a similarly elevated _p*K*a68,80._

The residues surrounding Asp49, which were designed using LigandMPNN^76^, were also important for activity. Asp49 is a part of a hydrogen-bonded “Asp Triad” network, which also includes Tyr9, and Gln75 (Fig. 3a). Replacement of either Tyr9 or Gln75 also greatly decreased the eliminase activity of KABLE1 (Supplementary Fig. 15). Thus, the Asp triad plays an essential role for the high activity of KABLE1.

We next used saturation mutagenesis to improve the catalytic efficiency of KABLE1 by 150-fold, to a value of 630,000 ± 19,000 M^-1^s^-1^. (Figs. 4b-d, Extended Data Table 1 and 2, Supplementary Fig. 14). This improvement was accomplished by combining mutations identified by single substitutions of all sidechains within 5.0 Å of the substrate. We also varied Lys14, Pro117, and Leu118, which are near slight kinks in **helices 1** and **4** proximal to the active site. The single mutants Ile12Phe, Lys14Pro, Leu20Trp, and Leu118Asn increased catalytic efficiency by 2- to 5-fold (Fig. 3b and Supplementary Table 5). These substitutions were functionally complementary when two, three or four of the mutants were combined (Fig. 3b, Supplementary Table 5, Extended Data Table 1). The catalytic efficiency was increased by two orders of magnitude in the quadruple mutant, KABLE1.4 (Fig. 3d, Extended Data Table 1 and 2). The improvement in the catalytic efficiency was primarily a result of an increase in *k*_cat_, with no statistically significant change in *K*_M_ (Extended Data Table 1 and 2). The apparent p*K_a_* of the pH-rate profile decreased from 9.3 **±** 0.2 in KABLE1 to 8.4 **±** 0.1 in KABLE1.4, approaching the pH of 8.0 of the buffer used to screen the mutants.

**Fig. 4.**
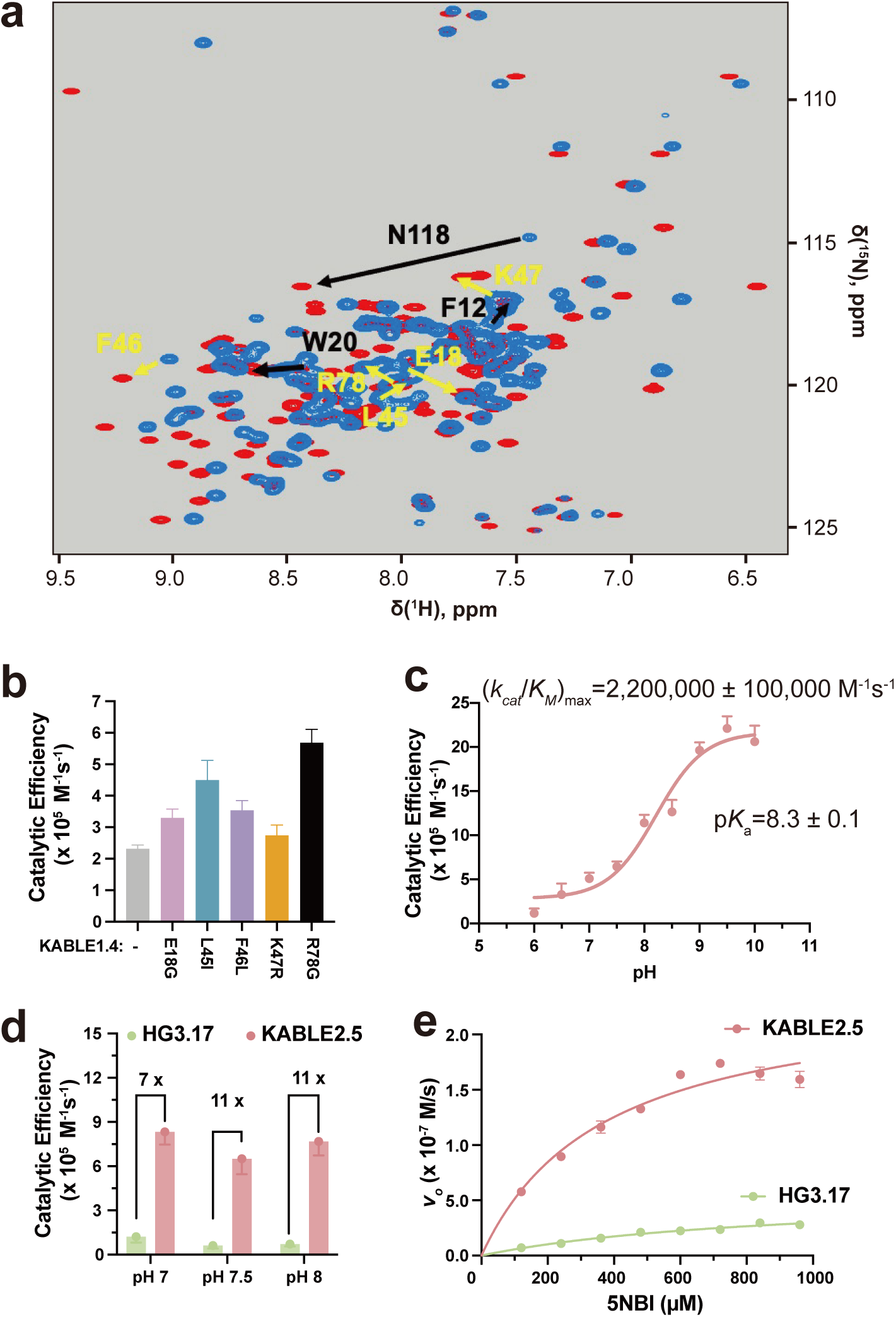
NMR-guided directed evolution of KABLE1.4. **a**, ^1^H-^15^N HSQC spectrum of KABLE1.4 in the absence (blue peaks) and presence (red peaks) of transition-state analogue (5 molar equivalents of 6NBT). Black arrows represent the beneficial residues identified from previous round of active site mutagenesis, and yellow arrows represent residues with high NMR perturbation, where the second set of beneficial mutations were found. **b,** Catalytic efficiency of KABLE1.4 and its beneficial mutants at 25 mM TRIS, pH 8.0, 100 mM NaCl, 1.5% MeCN. **c,** pH-dependence of *k*_cat_/*K*_M_ for KABLE2.5. *k*_cat_/*K*_M_ obtained from individual pH was used to fit the (*k*_cat_/*K*_M_)_max_ and p*K_a_*. **d, Bar plot representation for** catalytic efficiency of KABLE2.5 and HG3.17 at other reported conditions^61^ for HG3.17, including 50 mM sodium phosphate, pH 7.0, 100 mM NaCl, 10% MeOH, and 50 mM Bis-tris propane, pH 8.0,100 mM NaCl, 10% MeOH, and the condition at **e. e,** Michaelis-Menton curve of KABLE2.5 and HG3.17 at optimized condition^61^ for 50 mM Bis-tris propane, pH 7.5, 100 mM NaCl, 10% MeOH. Error bars in **b-e** represent the standard errors of the mean from three independent measurements.

We next improved the activity of KABLE1.4 by introducing substitutions at positions outside of the first shell of contacts using the method of NMR-guided directed evolution^65^. The amide proton resonances of a protein are particularly sensitive to small changes in local structures and can be used to identify residues whose environments are affected by the addition of a transition state analogue. Changes in chemical shift of residues not involved in first-shell interactions are particularly valuable, as they are believed to report structural changes associated with stabilization of the transition-bound state. As expected, the addition of 6NBT to KABLE1.4 induced large changes in its ^1^H-^15^N HSQC spectrum (Fig. 4a). Substitutions at these positions can often result in improved catalytic performance by helping to stabilize the activated complex^65,81^. We therefore performed saturation mutagenesis at 24 positions that show strong changes in chemical shift (Supplementary Fig. 16). Substitutions at five positions led to increases in catalytic efficiency ranging from 1.2 to 2.5 fold (Fig. 4b, Supplementary Table 5). Combining the mutants resulted in KABLE2.5, which had a 3.5-fold increase in *k*_cat_/*K*_M_ relative to KABLE1.4 (Fig. 4c, Extended Data Table 1 and 2). As was the case for KABLE1, we found that substitution of the active site base with an Asn abolished activity, and substitutions to the other members of the catalytic triad (Y9F and Q75M) resulted in over a 10-fold decrease in the value of *k*_cat_/*K*_M_ (Supplementary Fig. 17). *k*_cat_/*K*_M_ for KABLE2.5 showed a pH-dependence with a maximal value of 2,200,000 **±** 100,000 M^-1^s^-1^, and an apparent p*K*_a_ of 8.3 **±**. 0.1(Fig. 4c, Extended Data Table 1 and 2). A 2-fold increase in *k_ca_*_t_ from 290 **±** 70 s^-1^ to 570 **±** 20 s^-1^ contributed to the enhanced activity of KABLE2.5 vs. KABLE1.4 (Extended Data Table 2).

At the suggestion of a reviewer, we compared the activity of KABLE2.5 with that of HG3.17^61^, a protein that was designed starting with HG3^63^, and improved through 17 rounds of directed evolution. Under conditions that had been optimized for HG3.17’s activity at pHs 7, 7.5 and 8, we found that *k_cat_*/*K_M_* was 7 to 11-fold greater for KABLE2.5 than HG3.17 (Fig. 4d and 4e, Extended Data Table 3 and 4, Supplementary Fig. 18).

### Beneficial mutants immobilize and dehydrate the catalytic site

To understand the efficient catalysis of KABLE mutants, we determined structures by X-ray crystallography. The high water-solubility of KABLE variants inhibited crystallization. However, we were able to promote crystallization by introducing substitutions on the surface of the protein^25^, which allowed us to crystallize a surface-modified mutant containing the beneficial substitutions Lys14Pro and Leu20Trp mutants (designated KABLE1.2_cryst_, Supplementary Fig. 19). The crystallographic structure at 1.6 Å resolution solved in the absence and presence of 6NBT confirmed the overall structure of the design (0.8 Å Cα RMSD relative to the designed KABLE1.2; Supplementary Table 6 and Fig. 20).

In the crystal structure, the catalytic Asp triad was arranged as in the design model: the Asp49 carboxylate formed a strong (2.5 Å O – N distance) interaction with the transition state analog’s NH group; Gln75 formed a hydrogen bond to N2 of 6NBT’s trizole ring, and Tyr9 consolidates the triad by simultaneously interacting with Gln75, Asp49 and 6NBT (Fig. 5a). The structure also provided a rationale for the beneficial Lys14Pro and Leu20Trp mutants discovered from saturation mutagenesis. The indole of Leu20Trp formed a weak hydrogen bond (3.3 Å) with the nitro group of 6NBT. Furthermore, the helix-breaking Pro at position 14 locally disrupts the α-helix, enabling a backbone carbonyl to form a strong hydrogen bond to a water molecule, which in turn forms a hydrogen bond to the transition state analogue (Fig. 5b). Similar catalytically important waters have been observed in other Kemp eliminases^61^.

**Fig. 5.**
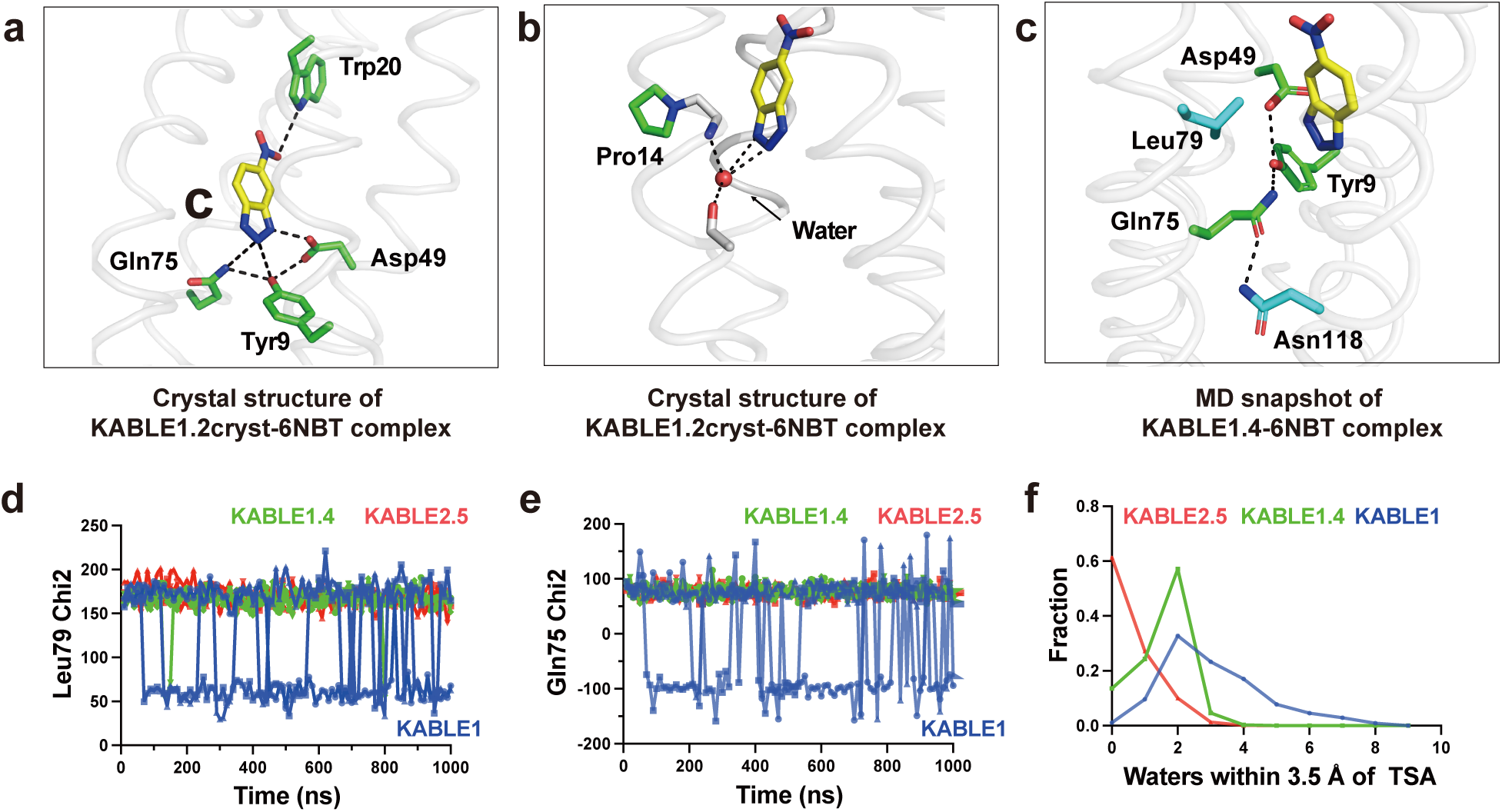
The origin of KABLE’s efficient catalysis probed by X-ray crystallography and molecular dynamics simulation. **a**, Structure of the KABLE1.2cryst complex (PDB: 9N0J) showing the Asp-triad engaging the triazole and a Trp indole NH engaging the nitro group at opposite sides of the bound 6NBT. **b**, Structure of the KABLE1.2_cryst_ complex (PDB: 9N0J). Pro14 disrupts the helical geometry, a backbone amide that binds a water molecule, which, in turn, forms two additional interactions with the triazole. **c**, Asp49-Tyr9-Gln75 triad (green carbons) and key residues, Leu79 and Asn118 (blue carbons). The structure is a representative snapshot from the MD simulation of KABLE1.4 complex with 6NBT (yellow carbons). **d-e**, The fluctuation of Chi2 angle of Gln75 (**d**) and Leu79 (**e**) versus time during MD simulation. KABLE1 sampled two distinct conformers, only one of which is found in the more active KABLE1.4 and KABLE2.5 trajectories. Three independent simulations were shown for each measurement. **f**, The number of waters within 3.5 Å of OD1 or OD2 of Asp49.

To probe the dynamic nature of the improvement in catalytic activity of the quadruple mutant, we performed three independent microsecond molecular dynamics (MD) simulations of the 6NBT complexes of KABLE1, with the corresponding mutants optimized by directed evolution, KABLE1.4 and KABLE2.5. Large changes in dynamics were observed between KABLE1 and KABLE1.4, which were consistent with the large (150-fold) improvement in its activity (Extended Data Table 2). Three active site residues, Gln75, Leu79, and Asn118 had greater mobility in KABLE1 than the more active KABLE1.4 (Figs. 5c-5e, Extended Data Fig. 6): Leu79 interacts with the aromatic ring of 6NBT (Fig. 5d); Gln75 is a member of the catalytic triad (Fig. 5e); and the Asn118 forms a hydrogen bond to Gln75 of the catalytic triad. (Figs. 5c and Extended Data Fig. 6). Thus, the active conformation of the Asp triad is more rigid in KABLE1.4, suggesting that beneficial mutants quench non-productive dynamic fluctuations in KABLE1. The MD simulations also confirmed the role of the two other beneficial mutants seen in the crystal structure of KABLE1.2_cryst_, including a water-mediated interaction near the helical kink caused by Lys14Pro and a hydrogen bond between Leu20Trp and 6NBT (Supplementary Fig. 21). Similar decreases in rigidity were observed comparing KABLE1 versus KABLE2.5.

The mean number of water molecules in contact with 6NBT decreased markedly between KABLE1 to KABLE1.4 (Fig. 5f), which is consistent with previous work showing that dehydration of a carboxylate base is important for Kemp eliminase activity ^82–84^. Interestingly, the mean hydration and heterogeneity of the environment surrounding 6NBT decrease even further in the more active mutant, KABLE2.5. Together, these findings provide strong support for our proposed catalytic mechanism and rationalize the improvements in activity that were observed during the two rounds of mutagenesis.

## Discussion

The evolution of life leveraged the emergence of primordial proteins that could bind, utilize, and create new small molecule metabolites and signaling molecules. These proteins evolve on a landscape that is often portrayed pictorially by projecting the features of sequence space into two dimensions^52,85,86^. Here, we expand this projection to understand binding specificity, by collapsing protein sequence and chemical structure to the two horizontal axes and representing fitness by the vertical height (Fig. 6a). Proteins that bind to and catalyze small molecules evolve along this landscape, as they acquire new functions. The ability to bind to new chemical entities, represented by moves in the axis of chemical structure, is enabled through alteration of their amino acid sequences (represented by the change in the axis of protein sequence). Here, we began with ABLE, which is specific for binding apixaban (sharp peak, Fig. 6a), progressed through an investigation of this same sequence’s low-level chemical promiscuity (broad peak, Fig. 6a), and ultimately used computational design to reach two new sequences KABLE2.5 and FABLE with different activities and specificities (sharp peaks, Fig. 6a).

**Fig. 6.**
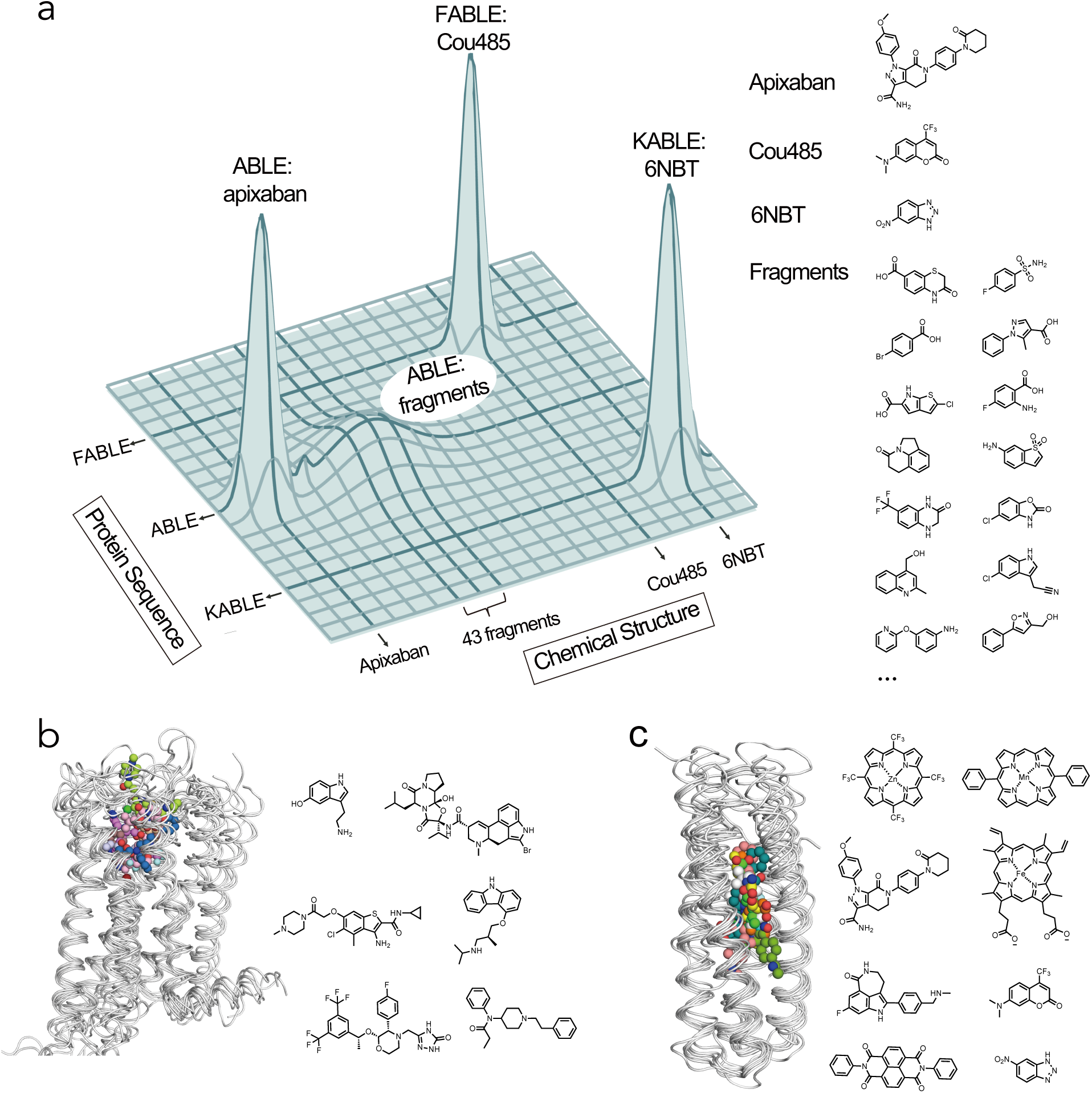
Exploration of sequence and chemical space of *de novo* protein-ligand interactions for designing new functions. **a**, Protein sequence diversity and small molecule chemical diversity are represented on the horizontal axes, and the heights of peaks denote fitness for binding or catalyzing the Kemp elimination reaction. Crystallographic fragment screening revealed ABLE as a generalist for binding 43 chemical fragments, which informed the design of the binding protein FABLE and the enzyme KABLE. **b-c**, Similar to GPCRs, the four-helical bundle scaffold has been used to bind different ligands. **b**, Overlay of GPCR complexes (left) and their diverse ligands (7WC4, 2RH1, 6EM9, 6OIK, 6VMS, 6HLO, 8EF5). **c**, Overlay of *de novo* designed four-helical bundle complexes and their diverse ligands (5GTY&7JH6, 7JRQ, 6W70, 7AH0&8D9P, 8TN6, 9DWC, 9DM3, 9N0J).

Searching broadly and simultaneously through both chemical and sequence space requires the exploration of two vast molecular landscapes. Most experimental work has previously focused on exploring just one of these two variables in isolation: drug discovery involves an extensive search of chemical space to discover small molecules or molecular fragments that bind to a target protein with a fixed sequence^-84^, while methods such as yeast display, phage display or directed evolution explore sequence space to discover new protein sequences for a given small molecule ^85,86^. *De novo* protein design provides an alternative approach to explore sequence space. Although *de novo* design of small molecule-binding proteins is still considered a significant challenge^90^, a number of proteins have been designed to bind drugs and cofactors^19,25,31,36,90–93^. However, it was previously unknown whether designed small molecule-binding proteins are more or less specific than natural proteins. *De novo* proteins have great conformational stability^25^ and their designed substrate-binding interactions are generally idealized. Together, these features might increase their specificity, making it difficult for them to act as generalists that can evolve new functions. Alternatively, they might have relatively low specificity because their binding sites have not been subject to the negative selective pressures that enforce specificity in a natural context. Here, we found that the *de novo* protein ABLE had promiscuity similar to natural proteins and that its promiscuous low-level binding could be used as a starting point for designing specific binding and catalytic functions.

In drug discovery, chemotypes are often discovered that bind to a target’s active site despite their bearing little resemblance to the target protein’s cognate substrate. Such hits can guide the design of novel inhibitors^87^. Here, we use fragment chemotype discovery to instead steer the design of proteins with novel activities. While we have used crystallographic fragment screening^34,50^, there are many experimental and computational methods for fragment discovery^87,88^ that could be brought to bear in future approaches to protein design.

The facile emergence of Kemp eliminase activity in the ABLE scaffold is noteworthy. Kemp eliminases have previously been elaborated starting with the active sites of natural proteins, but the attainment of high activity was achieved only after many rounds of directed evolution^60,63,64,72,73,81,71^(Fig. 4e). The differences in the ease of obtaining a novel enzymatic reaction from a *de novo* versus a natural protein as a starting point might reflect the fact that natural enzymes have been selected to not only promote their cognate reactions, but also avoid catalyzing off-target reactivities that might be detrimental to an organism’s survival. With a *de novo* scaffold, we begin with an open book without an evolutionary history predisposing towards one specific reactivity. It also freed us to match the scaffold with the reaction of interest^94–96^. The active site carboxylate of the KABLE1 series of proteins is placed within a deep cavity of the protein, which enhances the dehydration and increases the basicity of its active site carboxylate^82^. The enhancement of reactivity occurs at the expense of the stability of the protein, as the burial of a carboxylate into a hydrophobic environment is thermodynamically unfavorable. The kinetically-derived p*K*_a_ of Asp49 is four units higher than the unperturbed p*K*_a_ of Asp in water, corresponding to a free energy cost of 5.5 kcal/mol. Natural proteins are marginally stable, and hence cannot as easily withstand multiple modifications required to invince an entirely new function; by contrast, the extreme stability of *de novo* proteins^11,97^ enables such modifications. For example, KABLE1’s melting temperature is in excess of 95 °C (Supplementary Fig. 22).

Moreover, during improvement by saturation mutagenesis, KABLE1’s high thermodynamic stability also allowed the incorporation of multiple substitutions that were functionally beneficial but structurally destabilizing: Lys14Pro introduced a substitution that destabilizes alpha-helices by approximately 3 kcal/mol^98^; Leu20Trp introduced a steric clash in KABLE1 that required conformational rearrangements; and Leu118Asn replaced a hydrophobic interaction for an H-bonded interaction, which tends to be energetically unfavorable in analogous systems^71^. Despite these changes KABLE1.4 remained highly stable (Supplementary Fig. 22).

In pioneering work, Kuhlman and coworkers designed a Zn(II)-binding protein that was serendipitously found to catalyze ester hydrolysis^28^. Hilvert and coworkers evolved this metalloprotein to improve its hydrolytic activity^29^ and also to catalyze a Zn-promoted Diels-Alder reaction^30^. It is hard to directly compare these studies, because their catalytic activity was evoked by a collaboration between the protein matrix and the metal ion cofactor. Nevertheless, it is striking that our protein remained much closer in structure to the original design^19,28–30^, and the optimization of the FABLE and KABLE proteins was facilitated by the higher stability of the starting scaffold.

Nature has long used helical bundles as frameworks for enzymes that catalyze some of the most essential reactions for life, such as O_2_ formation and utilization^47,99^. They are also the most widely used small molecule binding and sensing devices. G-coupled protein receptors (GPCRs) bind small molecules of widely varying polarity, charge and hydrophobicity in a site that is bounded by helical bundles^43^ (Fig. 6b). Although their sequences are highly diverse, their structures are remarkably well conserved (Supplementary Table. 7)^100–102^. Similarly, *de novo* helical bundles represent a privileged scaffold for synthetic recognition of a wide range of small molecules, which include hemes^103–105^, synthetic cofactors^42,54,106^, and small molecule drugs^25^, in addition to the molecules studied here (Fig. 6c). Indeed, there is high similarity between the structures of ten *de novo* bundles designed in several different labs to bind a diverse array of small molecules (Supplementary Table. 8). Thus, in *de novo* design as in Nature, form follows function.

## Conclusion

This work advances at once the *de novo* design of proteins and our understanding of how proteins evolve from generalists to specialists. Here, we used: 1) information from fragment screening; 2) a deep mechanistic understanding of the reaction catalyzed^60,68,81,95^, and 3) state-of-the-art computational tools^55,76,78^ as full partners at each step of our work, from conception to realization of novel function. Thus, blending physical understanding with modern methods for computation design provides a powerful strategy for the design of sensors and catalysts.

## Methods

### ABLE crystallization and fragment Screening

Lyophilized ABLE was dissolved in water to a concentration of 18 mg/ml. Crystals were grown using sitting drop vapor diffusion in SwissCI 3-well plates (HR3-125, Hampton) with 200 nL protein and 200 nL reservoir (220 mM sodium malonate pH 5, 20% PEG 3350 and 3% DMSO). Needle-shaped crystals grew overnight at 19°C. We screened the 320 compound Enamine Essential fragment library against ABLE. Fragment solutions (40 nL of 100 mM stocks prepared in DMSO) were added to crystal drops using an acoustic liquid handler (Echo 650, Beckman Coulter)^109^. Crystals were incubated with fragments for 2-4 hours before being looped and vitrified in liquid nitrogen. X-ray diffraction data were collected at beamline 8.3.1 of the Advanced Light Source (ALS) and beamlines 12-1 and 12-2 of the Stanford Synchrotron Radiation Lightsource (SSRL). Data collection strategies are summarized in Supplementary File 1.

Diffraction images were indexed, integrated, and scaled with XDS and merged with Aimless^110,111^. In total we mounted 447 crystals soaked with fragments and were able to obtain high quality datasets from 321 crystals (resolution limit <2.2 Å based on CC_1/2_ < 0.3, *R*_free_ of initial model <35%). Replicate soaks were performed for some fragments: the total number of unique fragments with high-quality datasets was 242 (Supplementary File 1). We initially refined a model of ABLE against a dataset collected from a crystal soaked only in 10% DMSO. Phases were obtained by molecular replacement using Phaser^112^ and chain A of the previously published ABLE X-ray crystal structure as the search model (PDB: 6W6X)^36^. The initial model was improved by iterative cycles of refinement with *phenix.refine* (version 1.21.1-5286)^113^ and manual model building with COOT^114^. Waters were placed automatically in early-stage refinement using *phenix.refine* into peaks in the F_O_-F_C_ difference map >3.5 σ. In the later stages of refinement, waters were manually added or deleted, and hydrogens were refined with parameters constrained by those of the non-hydrogen atom (the riding hydrogen model). Data collection and refinement statistics are reported in Supplementary File 1. Coordinates and structure factor amplitudes were deposited in the PDB with accession code 9DW2. Refinement of the fragment-soaked datasets was performed using a pipeline based on Dimple as described previously using the DMSO-soaked ABLE model (PDB code 9DW2).^115,116^ The pipeline consisted of initial rigid body refinement with refinement followed by two cycles of restrained refinement in Refmac: the first with harmonic distance restraints (jelly-body restraints, four cycles) and the second with restrained refinement (eight cycles).

Fragment binding was detected using the PanDDA algorithm^35^ packaged in CCP4 version 7.0^117^. The background electron density map was calculated from 30 datasets (Supplementary File 1): six datasets were from crystals soaked in 10% DMSO and the remaining datasets came from fragment-soaked crystals where no fragment was detected. We collected multiple datasets for selected fragments. If fragments were detected in multiple datasets, we modeled the fragment with the highest occupancy based on the PanDDA event map and 1-BDC (Background Density Correction) value. Fragments were modeled into PanDDA event maps (Supplementary File 1) using COOT (version 0.8.9.2)^114^ and changes in protein conformation and solvent near the fragments were modeled. Fragment restraint files were generated with *phenix.elbow*^118^ from 3D coordinates generated by LigPrep (Schrödinger, version 2022-1) or Grade 2 (Global Phasing Ltd.). Structure refinement was performed with *phenix.refine* (version 1.20.1-4487) starting from the ABLE DMSO reference structure (PDB: 9DW2). Alternative conformations were modeled for residues when the RMSD exceeded 0.15 Å from the apo structure. The apo conformation was assigned alternative location (altloc) A and the fragment-bound conformation was assigned altloc B. Two fragments bound with alternative conformations that overlapped with the Tyr46 sidechain (PDB code 7HJK and 7HJX). For these fragments, residues 44-48 were modeled with a second conformation (altloc C). The multi-state models were initially refined with five *phenix.refine* macrocycles without hydrogens, followed by 10 macrocycles with riding hydrogens. Occupancy of the fragment- and apo-states were refined at either 2*(1-BDC) ^35,119^ or at fragment occupancy values from 10-90% at 10% increments. Fragment occupancy was determined by inspection of mF_O_-DF_C_ difference map peaks after refinement. The occupancy at 2*(1-BDC) was appropriate for 41/43 fragments (Supplementary File 1). Coordinates and structure factor amplitudes have been deposited in the PDB with accession codes 7HIY, 7HIZ, 7HJ0, 7HJ1, 7HJ2, 7HJ3, 7HJ4, 7HJ5, 7HJ6, 7HJ7, 7HJ8, 7HJ9, 7HJA, 7HJB, 7HJC, 7HJD, 7HJE, 7HJF, 7HJG, 7HJH, 7HJI, 7HJJ, 7HJK, 7HJL, 7HJM, 7HJN, 7HJO, 7HJP, 7HJQ, 7HJR, 7HJS, 7HJT, 7HJU, 7HJV, 7HJW, 7HJX, 7HJY, 7HJZ, 7HK0, 7HK1, 7HK2, 7HK3 and 7HK4. Structure factor intensities (unmerged, merged, and merged/scaled), PanDDA input and output files including Z-map and event maps in CCP4 format, and refined models including the fragment-bound state extracted from multi-state models were uploaded to Zenodo (DOI: 10.5281/zenodo.13913848).

### Computational design of fluorescent ABLE (FABLE)

With (7-hydroxylcoumarin)-ABLE structure (PDB: 7HIY) available from fragment-screening, we superimposed Cou485 to 7-hydroxycoumarin fragment based on shared heavy atoms to obtain the initial pose, which contains the Tyr46 packing with 2H-1-benzopyran ring of coumarin and hydrogen bond between His49 and ketone of coumarin. This initial pose was used for Rosetta flexible backbone sequence design, using a previously reported script in our lab^54^ (Supplementary Note, Computational design of FABLE). During the sequence design, the side chain conformation of His49 and Tyr46 were fixed and the geometry constraint for the His49 interaction with ketone of Cou485 was applied. Out of the total 126 residues, 43 residues in the vicinity of apixaban and 7-hydroxylcoumarin binding site were chosen to be designed to reshape the binding pocket towards Cou485 while keeping the portion of the packing core of ABLE that is not involved in binding ligands unchanged. We selected the 100 best-scoring designs (lowest Rosetta energy score, using the Rosetta Ref2015 energy function) from 1000 design outputs. We next carried out structure prediction using OmegaFold^56^ and ESMFold^79^ to further filter out the designs whose predicted structures do not agree with the designed structures. Sequences with a less than 1.0 Å RMSD difference between the design model and prediction as well as a plDDT score over 90 were picked for detailed structural evaluation. Final sequences were selected by additional criteria: 1) No more than 2 unsatisfied side-chain hydrogen bonds in the interior of design; 2) Pre-organized binding pocket from the prediction, especially the side-chain conformation of Tyr46 and His49, which is quantified by the sub-Å RMSD of CA, CB, CG between the designed model and AF3 prediction. AF3 server (https://alphafoldserver.com) was used to predict the structures for selected designs (Supplementary Fig. 6), although the program was not available at the time that the sequences were designed.^120^

### Computational design of KABLE (Kemp eliminase)

Our goal was to use the information from the fragment complexes to inform the design of the active site. As the substrate was a bicyclic aromatic compound, it was logical to position it into the aromatic boxes of the A site, hosting 8 bicyclic fragments. This was followed by searching for a Glu or Asp sidechain that was well positioned to abstract a proton from the substrate to catalyze the elimination reaction (Strategy 1). The second strategy centered instead on using the information from fragment complexes to first identify a catalytically favorable position for a Glu/Asp transition state complex, followed by redesign of ABLE’s binding site to stabilize the bicyclic ring, primarily through hydrophobic, aromatic interactions and van der Waals interactions.

In each case we considered theoretical suggestions that emphasize the location of a binding site relative bulk solvent. As early as 1967 Perutz stated: “The non-polar interior of enzymes provides the living cell with the equivalent of the organic solvents used by the chemists. The substrate may be drawn into a medium of low dielectric constant in which strong electric interactions between it and specific polar groups of the enzyme can occur.” Thus, it is important to consider placement of the base in a pre-organized, solvent-inaccessible location well below the surface of the protein. This principle has been supported by a body of theoretical and computational studies, as well as through careful examination of evolved Kemp eliminases^61,63,64,83^. We satisfied Perutz’s this principle by choosing designs in which the carboxylate base and the substrate were fully inaccessible to a water-sized probe, the surrounding sidechains should be primarily hydrophobic^63,71^ and the geometry of the carboxylate should be close to the saddle point for the computed transition state of the reaction.

#### Strategy 1: Aromatic box

Inspired by the aromatic boxes of conformer A, we used a similar procedure for design of a Kemp Eliminase. For the aromatic box in conformer A, we used 7HJQ as the starting structure. The plane of the ring of 6NBT was oriented as in 7HK4. The next step was to identify a position for incorporation of the active site Asp or Glu to satisfy the following criteria: **1)** When in a low-energy rotamer, its carboxylate should form a hydrogen bond to N3 of 6-nitrobenzotriazole (6NBT) in a geometry computed from previous quantum calculations^60,121,122^ **2)** The benzisoxazole ring of the substrate should be fully buried (to increase the basicity of the carboxylate). Thus, the Asp/Glu carboxylate should be solvent-inaccessible. We also filtered out any examples where the Cα of the acidic residue is located at a surface position, but the sidechain reaches into the protein interior, because the burial of the carboxylate would need to compete with solvation. **3)** The introduced sidechain should not clash with the backbone or sidechains of the residues comprising the aromatic box. This left 3 positions to explore, namely 108, 112 and 115. After searching through low-energy sidechain rotamers of Asp and Glu at these locations, Glu at position 108 was found to uniquely fulfill these criteria. The resulting complex was used as an input structure for sequence design with either an 1) an iterative round of LigandMPNN^76^ and Rosetta FastRelax^55^ or a 2) classical flexible backbone Rosetta sequence design.

1. For iterative rounds of LigandMPNN and Rosetta FastRelax, residue 46 was fixed to either Y or W and residue 86 was fixed to F to maintain the aromatic packing interactions discovered in fragment screening. In the Rosetta FastRelax step, geometric constraints from the reported crystallographic conformation sampling were applied (Supplementary Note, constrain for active site 1 of kemp eliminase).^123^
2. The second route using the flexible backbone Rosetta sequence design proceeded with the same constraint applied (Supplementary Note, constrain for active site 1 of kemp eliminase) following the reported protocol from our lab^54^.

Structure prediction with RaptorX was carried out for all designed sequences^78,124^, to ensure that the designed structures would be preorganized to adopt the same active site geometry of the design model. Of 2000 sequences evaluated on 1), we chose designs (1, 4 and 5). Of 2000 sequences evaluated on 2), we chose designs 2 and 3. The selection criteria were the sub-Å backbone RMSD between designed complex and the predicted unliganded structures and also agreement of sidechain conformation of active site (residue 108, 46, 79, 86) between design and prediction.

#### Strategy 2: Hotspot residue 49

In strategy 2 we focused on identifying a site that was well-positioned to accommodate a Glu/Asp as the general base. His49 was selected as a potentially ideal position, because it was situated at a central location near the bottom of the binding site where it donated a hydrogen bond to a carbonyl of apixaban in ABLE. His49 also formed hydrogen bonds to 35% fragments binding at the pocket of ABLE, and it formed imidazole-carboxylate salt bridges in 9 of the complexes, as well as with acetate in 6W6X (Supplementary Fig. 9). We reasoned that it should be possible to reverse the polarity of the interaction by replacing the His with an Asp, which is nearly isosteric to histidine, as this would place the sidechain in a good environment to form a strong hydrogen bond. 6NBT was docked to the Asp49 of ABLE-fragment crystal structures (PDB: 7HJV) based on previously reported geometric constraints from DFT calculations, considering the more basic (syn) lone pairs of electrons on either OD1 or OD2 for proton abstraction^60^.

All 3 resulting poses (Supplementary Fig. 9) were used as input structures for sequence design with 3 iterative rounds of LigandMPNN and Rosetta FastRelax, resulting in 1000 designs per input pose. In the Rosetta FastRelax step, geometric constraints from the reported crystallographic conformation sampling were applied (Supplementary Note, constrain for versatile residue 49 of kemp eliminase)^123^. Structure prediction with RaptorX was carried out for all designed sequences ^78,124^. Five designs were picked based on the goodness of fit between the predicted model of the uncomplexed structure versus the design model for the complex as described in the section above.

### Molecular dynamics (MD) simulation

#### General MD procedure

Proteins and ligands were parameterized using Gaussian09 and AmberTools’s Antechamber program in Amber22^125,126^. Ligand Partial charges for ligand were determined by first optimizing its geometries at the B3LYP/6- 31G* level^126^, and then calculating its electrostatic potentials using the Merz-Singh-Kollman method in Gaussian09^126^. Charge fitting was then performed using Antechamber’s RESP program within AmberTools^127,128^. All other small molecule parameters were assigned by Antechamber based on the GAFF2 database^129,130^. The starting structures for the MD simulations are described in the following section. The simulation box was built by solvating protein with OPC-modeled waters ^131^ in a box with 8 Å padding from the protein, and sodium and chloride ions were added to reach a charge-neutral 137 mM NaCl concentration to match experimental conditions. Simulations began with 1,000 restrained steepest-descent minimization steps before switching to a maximum of 6,000 steps in conjugate gradient steps. The system was then heated up to 278 K over 50 ps in the NVT ensemble with Langevin thermostat control of temperature and 1 fs timestep. The simulation was then switched to the NPT ensemble, and pressure was maintained at 1 atm using the Monte Carlo barostat ^132^. Throughout equilibration steps, protein and ligand heavy atoms were initially restrained with harmonic potentials at 10 kcal/(mol·Å2) and ramped down to 0 kcal/(mol·Å2) over 9 equilibration steps, totaling 1 nanosecond. Each simulation was then carried out for an unrestrained production run under periodic boundary conditions with 2 fs timesteps. The SHAKE algorithm^133,134^ was used to restrain hydrogens, short-range non-bonded electrostatic and Lennard-Jones interactions were cut off at 10 Å, and the Particle Mesh Ewald method (Ref*X13*) was used for long-range electrostatics.

#### MD for ABLE

The starting structures for the simulations were derived from the crystal structures of ABLE complexed with apixaban (PDB 6W70, chain B), and apo ABLE (PDB 6W6X, chain A)^36^. Both systems were prepared using the epsilon nitrogen-protonated state of H49 (“HIE”). The simulation was carried out at 278K for 2 independent runs with each run of 500 ns. The final trajectory was extracted with a 5 ns interval, which resulted in 100 states for each run of 500 ns. Chi1 and Chi2 angles of Tyr46 were extracted with an in-house script (Supplementary Note, extract_chi1_chi2.py) and manual inspection was also done to verify the correct calculation of Chi2 angle due to the symmetry of tyrosine sidechain. Chi1 and Chi2 of His49 were extracted with the same script. Contours were generated from the Chi1 and Chi2 angles seen at the simulation using an in-house script (Supplementary Text, generate_contour_from_tyr_chi1and2.py), which utilizes the Gaussian kernel density estimate methods from scipy.stats module of SciPy Python Packages.

#### MD for KABLEs

The starting structure for the KABLE1 simulations was derived from the design structures of KABLE1, in complex of 6NBT. For the simulation of KABLE1.4 with four mutations, Rosetta FastRelax was used to incorporate these mutations and the relaxed structures were used for MD simulation. The simulation was carried out at 300 K for 3 independent runs with Berendsen barostat and each run lasted for 1000 ns. The final trajectory was extracted with 1 ns interval, which resulted in 1000 states for each run of 1000 ns. Chi1 and Chi2 of Gln75 and Leu79 were extracted using the script (Supplementary Note, extract_chi1_chi2.py). The number of waters within 3.5 Å of 6NBT was calculated using the script (Supplementary Note, count_water.py).

## Acknowledgments

We are grateful for helpful discussions with Sam Schneider and Hyunil Jo. We thank members of the DeGrado lab and Fraser lab for support. The synchrotron X-ray diffraction data used to determine crystal structures reported in this work were collected at beamline 8.3.1 of the Advanced Light Source (ALS) and beamlines 12-1 and 12-2 of the Stanford Synchrotron Radiation Lightsource (SSRL). We acknowledge the use of the Wynton high-performance compute cluster at UCSF.

## Author contributions

Y.C., S.B., N.F.P., J.S.F., and W.F.D. formulated the project; L.B., G.J.C.and J.B. perfomed the fragment-screening, Y.C. performed the computational design and experimental characterization of FABLE; Y.C. conducted the computational design of KABLE. S.B. and Y.C. carried out experimental optimization and characterization of KABLEs. S.B. conducted the substrate specificity study of KABLE1.4, L.B., G.J.C. and Y.C. obtained the crystal structure of the FABLEs and KABLEs. Y.C. and S.T. perfomed MD simulation. K.H., L.L. and I.B. assisted with computational design. Y.C., S.B., G.J.C., K.H., N.F.P., J.S.F., and W.F.D wrote the paper with input from all authors.

## Data and materials availability

All data are available in the main text or the supplementary materials.

Supplementary Information is available for this paper.

Correspondence and requests for materials should be addressed to James S. Fraser, and William F. DeGrado.

