## Extended Data for "Emergence of specific binding and catalysis from a designed generalist binding protein"

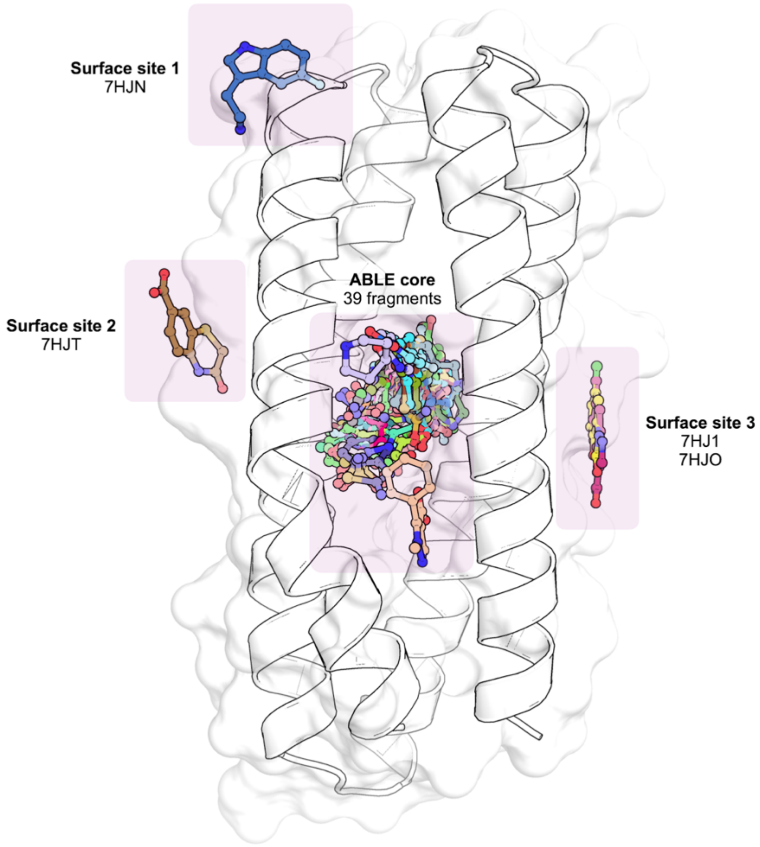


**Extended Data Fig. 1 |** X-ray crystal structure of ABLE showing binding sites for 43 fragments. The structure of apo ABLE (PDB: 9DW2) with white cartoon and transparent white surface.

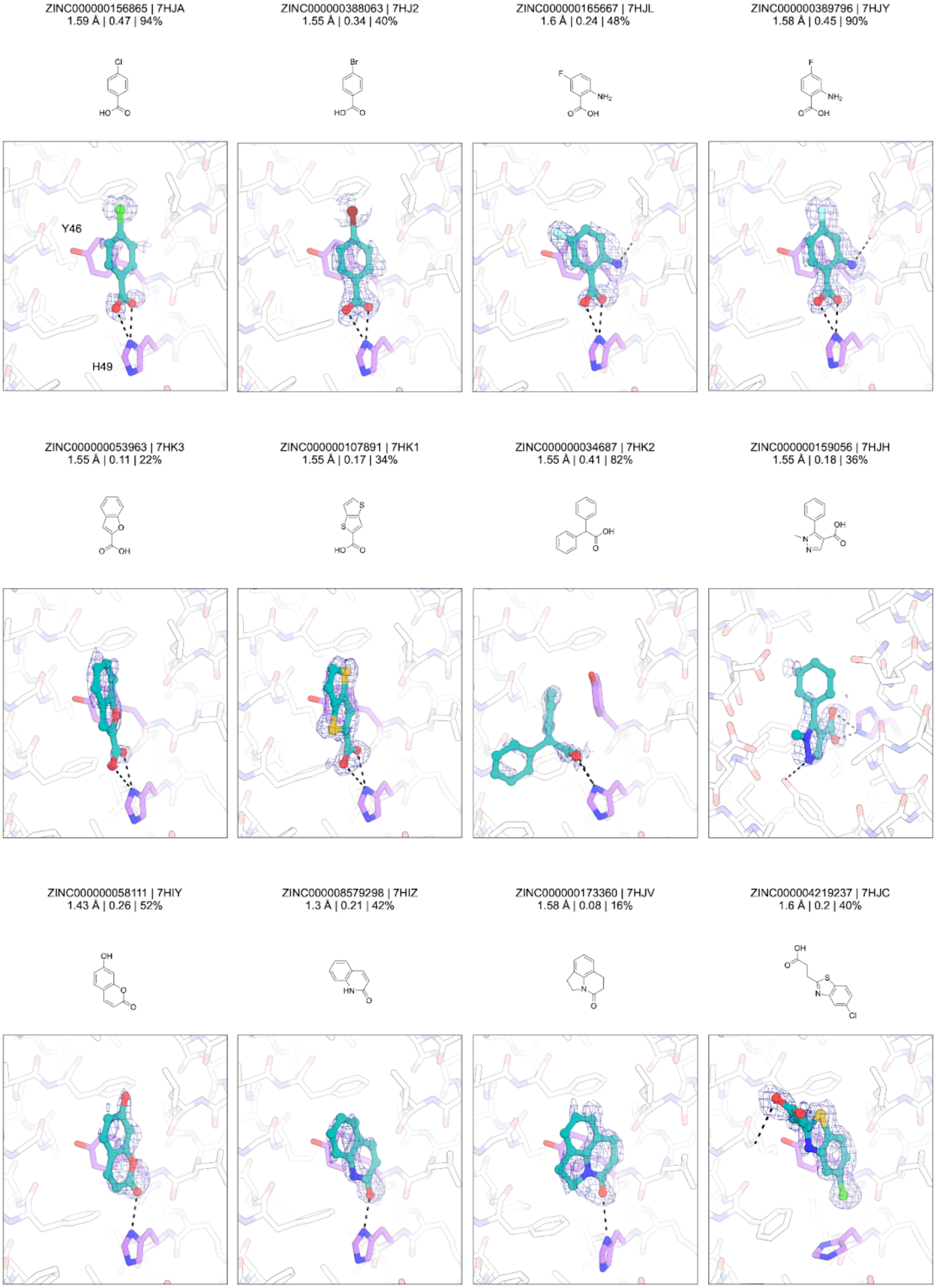

ZINC000000156865 | 7HJA
1.59 Å | 0.47 | 94%
ZINC000000388063 | 7HJ2
1.55 Å | 0.34 | 40%
ZINC000000165667 | 7HJL
1.6 Å | 0.24 | 48%
ZINC000000389796 | 7HJY
1.58 Å | 0.45 | 90%
Y46
H49
ZINC000000053963 | 7HK3
1.55 Å | 0.11 | 22%
ZINC000000107891 | 7HK1
1.55 Å | 0.17 | 34%
ZINC000000034687 | 7HK2
1.55 Å | 0.41 | 82%
ZINC000000159056 | 7HJH
1.55 Å | 0.18 | 36%
ZINC000000058111 | 7HIY
1.43 Å | 0.26 | 52%
ZINC000008579298 | 7HIZ
1.3 Å | 0.21 | 42%
ZINC000000173360 | 7HJV
1.58 Å | 0.08 | 16%
ZINC000004219237 | 7HJC
1.6 Å | 0.2 | 40%

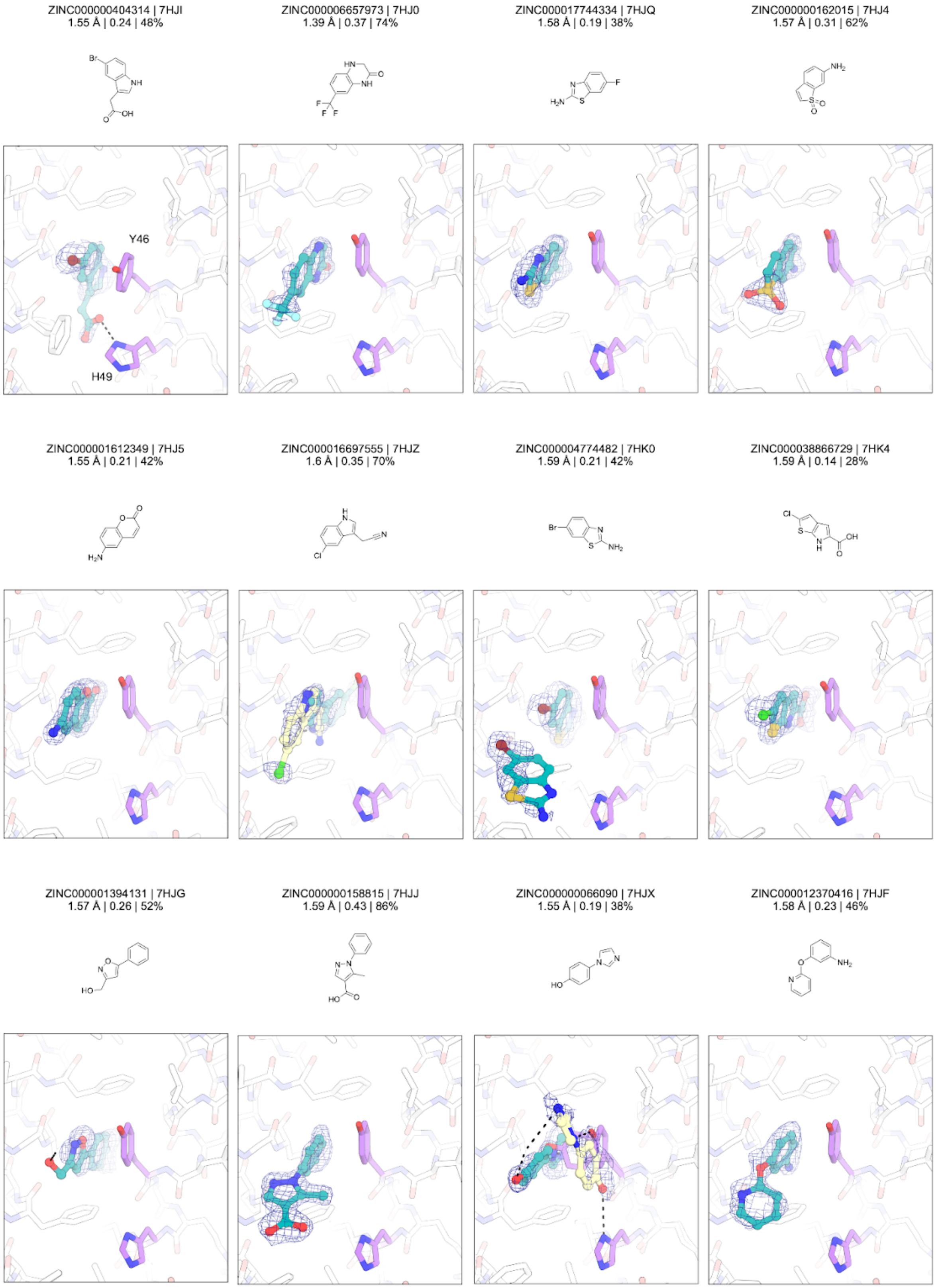

ZINC000000404314 | 7HJI
1.55 Å | 0.24 | 48%
ZINC000006657973 | 7HJ0
1.39 Å | 0.37 | 74%
ZINC000017744334 | 7HJQ
1.58 Å | 0.19 | 38%
ZINC000000162015 | 7HJ4
1.57 Å | 0.31 | 62%
Y46
H49
ZINC000001612349 | 7HJ5
1.55 Å | 0.21 | 42%
ZINC000016697555 | 7HJZ
1.6 Å | 0.35 | 70%
ZINC000004774482 | 7HK0
1.59 Å | 0.21 | 42%
ZINC000038866729 | 7HK4
1.59 Å | 0.14 | 28%
ZINC000001394131 | 7HJG
1.57 Å | 0.26 | 52%
ZINC000000158815 | 7HJJ
1.59 Å | 0.43 | 86%
ZINC000000066090 | 7HJX
1.55 Å | 0.19 | 38%
ZINC000012370416 | 7HJF
1.58 Å | 0.23 | 46%

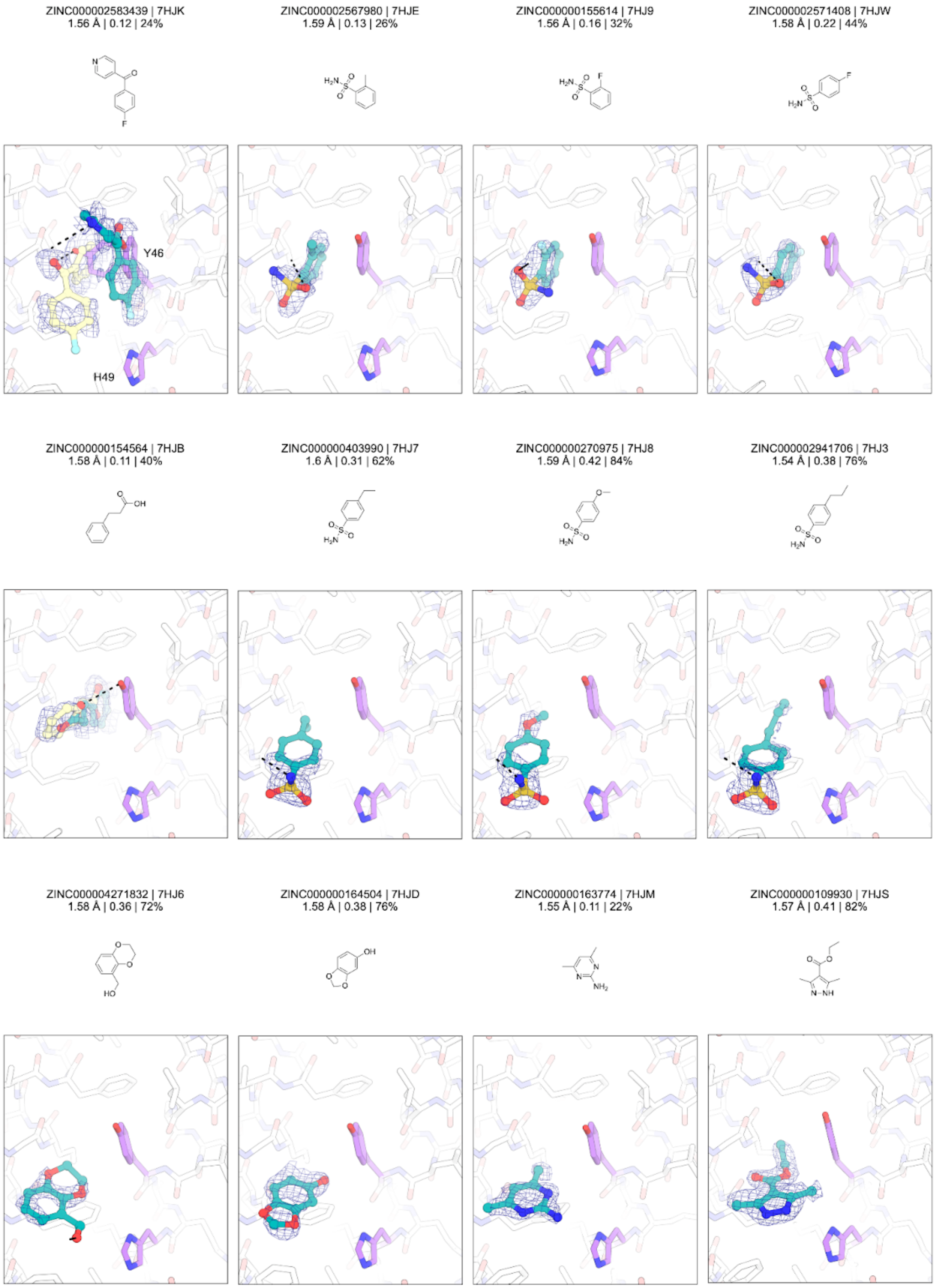
ZINC000002583439 | 7HJK
1.56 Å | 0.12 | 24%
Y46
H49
ZINC000002567980 | 7HJE
1.59 Å | 0.13 | 26%
ZINC000000155614 | 7HJ9
1.56 Å | 0.16 | 32%
ZINC000002571408 | 7HJW
1.58 Å | 0.22 | 44%
ZINC000000154564 | 7HJB
1.58 Å | 0.11 | 40%
ZINC000000403990 | 7HJ7
1.6 Å | 0.31 | 62%
ZINC000000270975 | 7HJ8
1.59 Å | 0.42 | 84%
ZINC000002941706 | 7HJ3
1.54 Å | 0.38 | 76%
ZINC000004271832 | 7HJ6
1.58 Å | 0.36 | 72%
ZINC000000164504 | 7HJD
1.58 Å | 0.38 | 76%
ZINC000000163774 | 7HJM
1.55 Å | 0.11 | 22%
ZINC000000109930 | 7HJS
1.57 Å | 0.41 | 82%

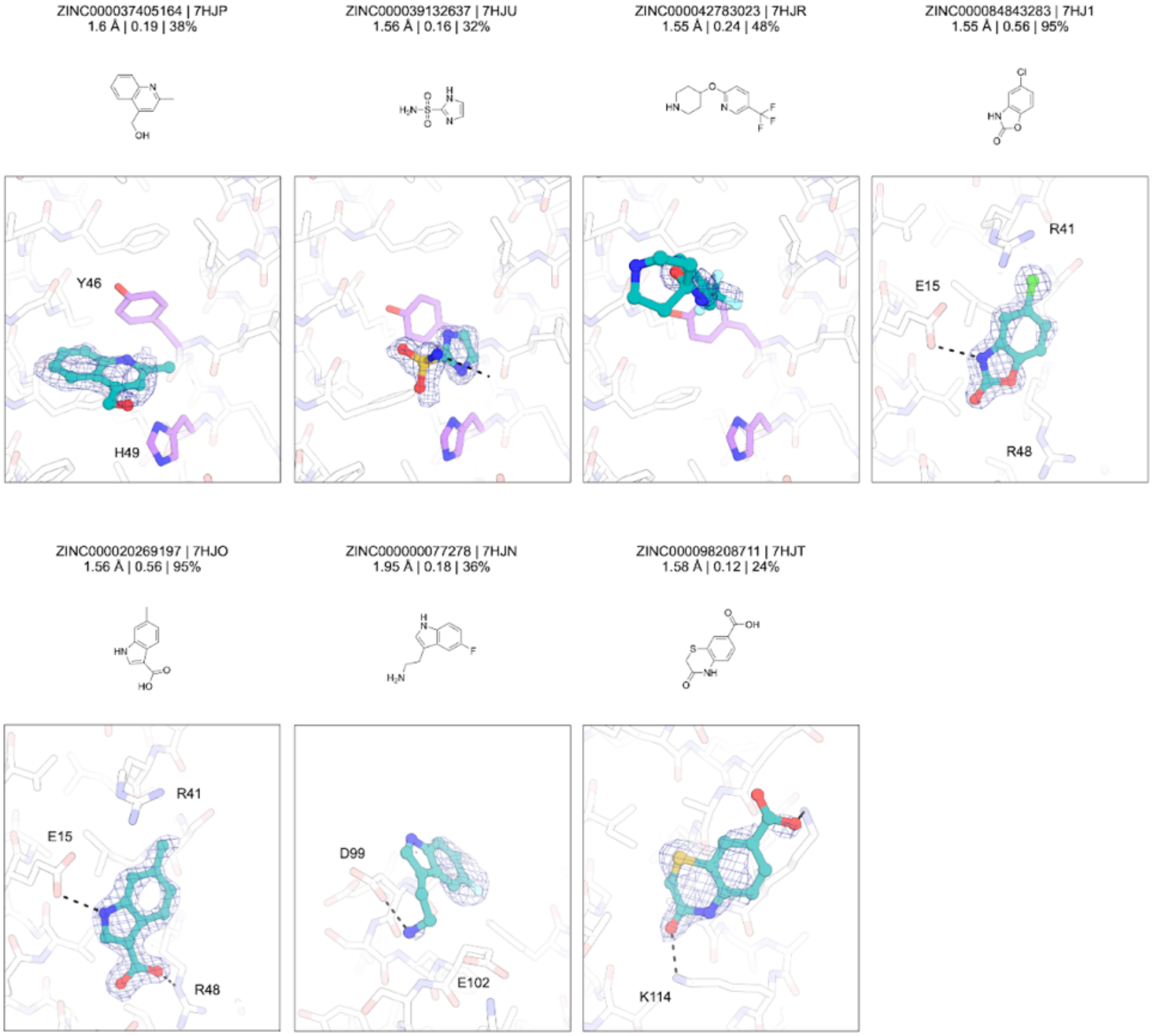


**Extended Data Fig. 2 | Chemical structures and electron density maps for the 43 fragment structures reported in this work**. PanDDA event maps (blue mesh, 2 σ) are contoured around fragments (teal/yellow sticks). The sidechains of residues Tyr46 and His49 are shown with purple sticks. Fragment names, PDB codes, resolution, 1-BDC value and refined occupancies are indicated. Hydrogen bonds are shown with dashed black lines. For clarity, residues 10-23 and 105-118 are hidden for fragments binding in the ABLE core. His49-mediated polar interaction could be found in the majority of fragments at site B.

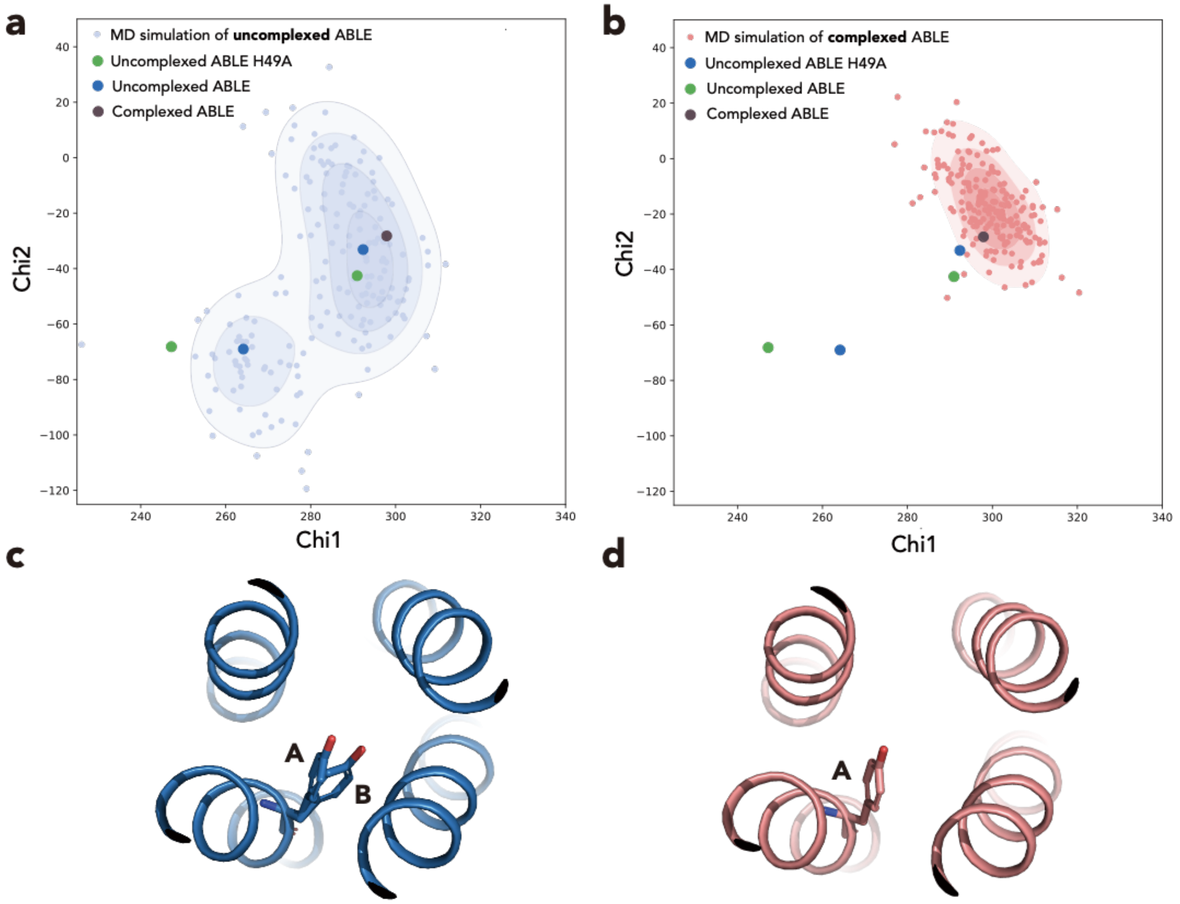


**Extended Data Fig. 3 | Contour map of Tyr46 side chain conformation of ABLE from molecular dynamics simulation.** (**a&b**) Dynamics of uncomplexed ABLE (PDB: 6W6X) and ABLE-apixaban (PDB: 6W70) were explored by doing molecular simulation at 278 K for 500 ns with Amber. Chi1 and Chi2 from each state of these simulations and reported crystal structures of ABLEs (6W6X: uncompleted ABLE; 6W70: ABLE-Apixaban Complex; 6X8N: uncomplexed ABLE His49Ala) were extracted for plotting. Contours were generated from the points of uncomplexed ABLE (**a**) and ABLE-apixaban complex contours (**b**) were generated using an in-house script that utilizes the Gaussian kernel density estimate methods from scipy.stats module of SciPy Python Packages. (**c&d**) **A** and **B** conformations of Tyr46 sidechain were present in the uncomplexed ABLE crystal structure (**c**, PDB: 6W6X) while only A conformation of Tyr46 sidechain conformations was present at ABLE-apixaban complex (**d**, PDB: 6W70).

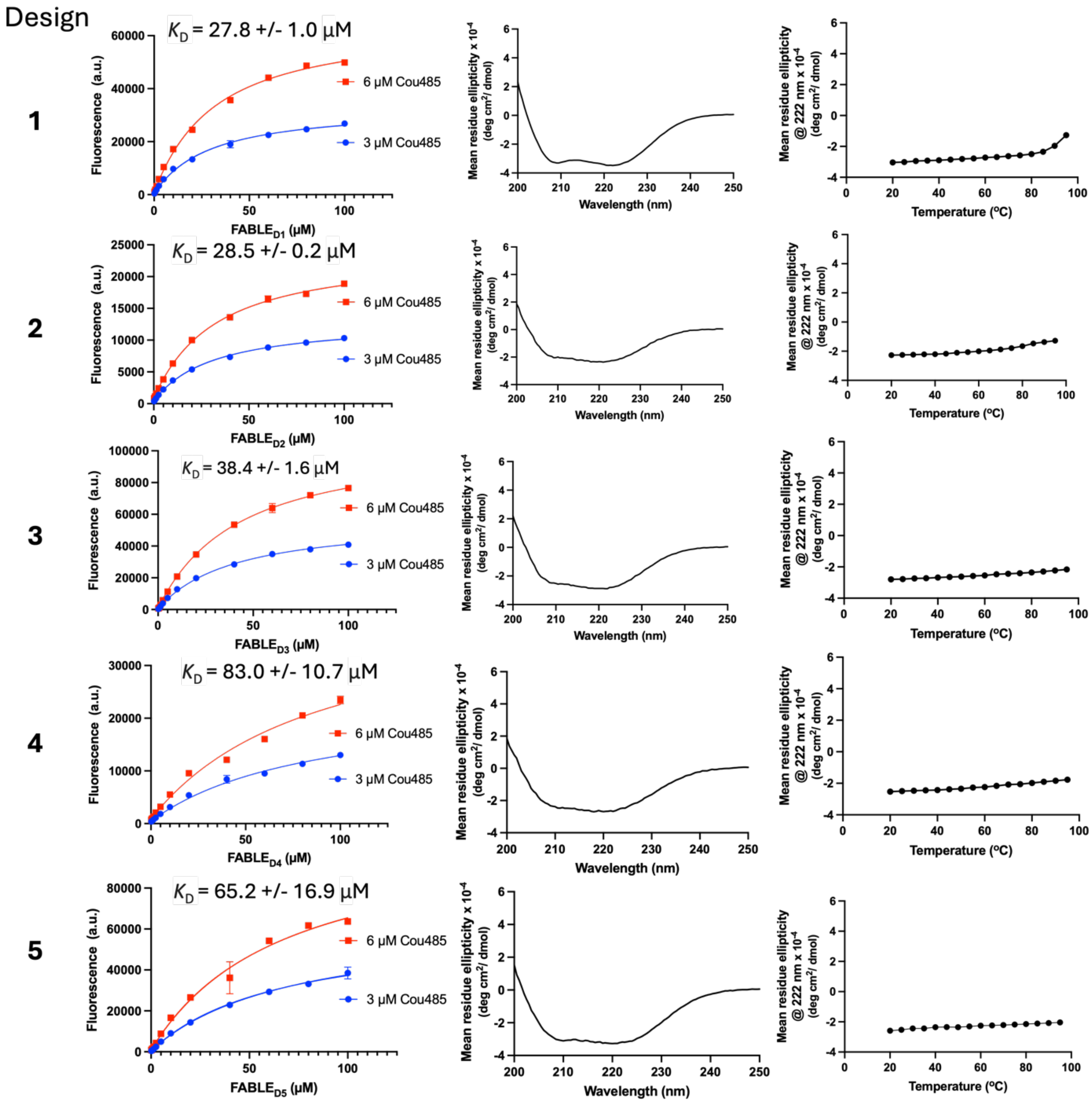


**Extended Data Fig. 4 | Characterization of five FABLE Designs.** Left column: Fluorescence of Cou485 at two fixed concentrations of 3 μM and 6 μM were measured in the presence of increasing amounts of each protein. The dissociation constant was obtained by globally fitting a single-site binding model to data, as described at Supplementary Methods. The error bars represent standard deviations of three independent measurements. Middle Column: circular dichroism spectra shows that all the designs are helical proteins. Right column: temperature-dependent circular dichroism signals measured at 222 nm show that all designs are thermostable. The design 1 was designated as FABLE.

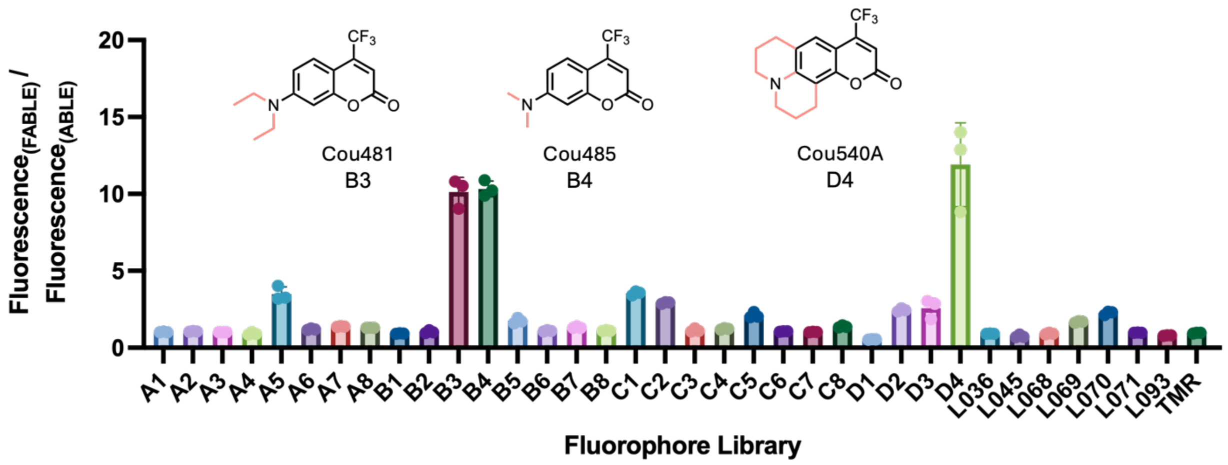


**Extended Data Fig. 5 | Exploring the chemical space of FABLE.** The excitation and emission of fluorophores are in Supplementary Table 3. The error bars represent standard deviations.

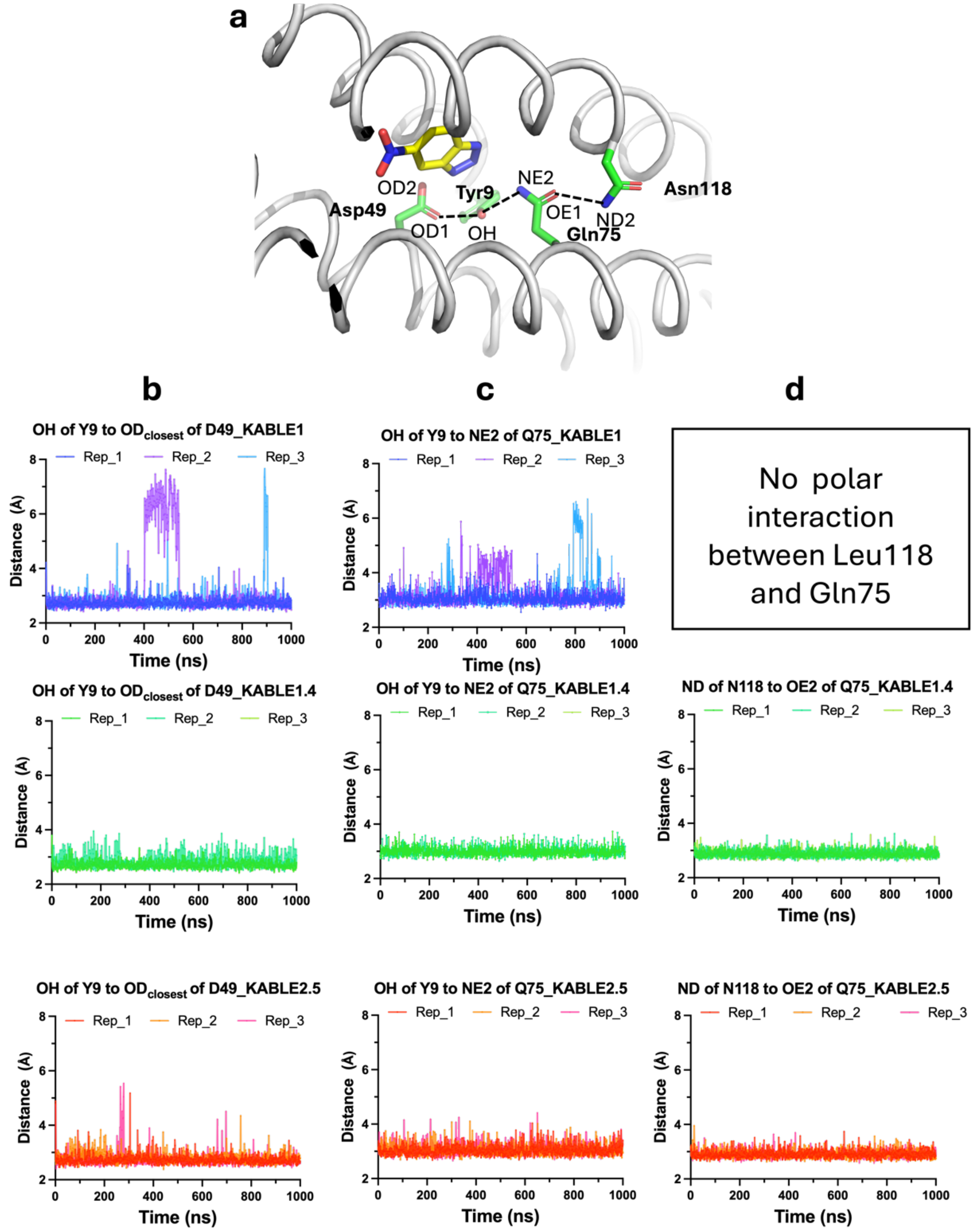


**Extended Data Fig. 6 | Stabilization of triad by Leu118Asn.**

**a,** A Representative snapshot of molecular dynamic simulation of KABLE1.4-6NBT complex showing the triad consisting of Asp49, Tyr9, Gln75 and its stabilization residue Asn118, whose sidechain are shown in green sticks. The hydrogen bonding between these 4 residues are annotated with dotted line and their distance are within 2.8-3.2 Å. **b-d,** The fluctuation of indicated distances between indicated atoms at the process of 1000 ns MD simulation of KABLE1.4-6NBT (green) complex, KABLE2.5-6NBT (red) complex, in comparison with KABLE1-6NBT (blue) complex. The distances are extracted using the *label bond* function of VMD. Three independent MD runs are show as three separate curves.

**Extended Data Table 1 | pH-dependence of kinetic parameters for KABLEs[a].**

| Protein | pH | $k_{cat}$, $s^{-1}$ | $K_M$, mM | ($k_{cat}/K_M$), $M^{-1}s^{-1}$ |
|---|---|---|---|---|
| KABLE1 | 7.0 | 0.010 ± 0.0006 | 0.15 ± 0.04, | 68 ± 17 |
| | 7.5 | 0.030 ± 0.0007 | 0.17 ± 0.02, | 176 ± 17 |
| | 8.0 | 0.09 ± 0.001, | 0.20 ± 0.01 | 460 ± 20 |
| | 8.5 | 0.11 ± 0.001 | 0.22 ± 0.01 | 480 ± 20 |
| | 9.0 | 0.39 ± 0.03 | 0.12 ± 0.05 | 3,300 ± 130 |
| | 9.5 | 1.3 ± 0.2 | 0.50 ± 0.1 | 2,600 ± 800 |
| | 10 | 1.3 ± 0.2 | 0.35 ± 0.16 | 3,700 ± 1,800 |
| KABLE1.4 | 6.5 | 10 ± 0.7 | 0.18 ± 0.05 | 58,000 ± 16,000 |
| | 7.0 | 30 ± 1.6 | 0.37 ± 0.05 | 80,000 ± 12,000 |
| | 7.5 | 65 ± 5 | 0.38 ± 0.03 | 170,000 ± 20,000 |
| | 8.0 | 99 ± 5 | 0.43 ± 0.05 | 230,000 ± 32,000 |
| | 8.5 | 180 ± 6 | 0.49 ± 0.03 | 360,000 ± 28,000 |
| | 9.0 | 210 ± 4 | 0.40 ± 0.02 | 530,000 ± 26,000 |
| | 9.5 | 270 ± 3 | 0.44 ± 0.01 | 610,000 ± 16,000 |
| | 10 | 290 ± 5 | 0.49 ± 0.02 | 610,000 ± 28,000 |
| KABLE2.5 | 6.5 | 36 ± 3 | 0.11 ± 0.04 | 330,000 ± 130,000 |
| | 7.0 | 140 ± 5 | 0.25 ± 0.03 | 510,000 ± 67,000 |
| | 7.5 | 210 ± 6 | 0.30 ± 0.03 | 640,000 ± 58,000 |
| | 8.0 | 230 ± 5 | 0.20 ± 0.02 | 1,100,000 ± 92,000 |
| | 8.5 | 340 ± 10 | 0.27 ± 0.03 | 1,300,000 ± 140,000 |
| | 9.0 | 470 ± 7 | 0.24 ± 0.01 | 2,000,000 ± 93,000 |
| | 9.5 | 544 ± 10 | 0.25 ± 0.01 | 2,200,000 ± 140,000 |
| | 10 | 570 ± 20 | 0.27 ± 0.02 | 2,100,000 ± 180,000 |

[a]Error bars represent the standard errors of the mean from three independent measurements.

**Extended Data Table 2 | Analysis of pH-dependence of kinetic parameters for KABLEs[a]**

| **Protein** | pH range | $k_{cat}$, $s^{-1}$ | ($k_{cat}/K_M$), $M^{-1}s^{-1}$ | p$K_a$ [b] |
|---|---|---|---|---|
| KABLE1 | 7.0-10 | 1.7 ± 0.2 | 3,800 ± 1,400 | 9.3 ± 0.2 |
| KABLE1.4 | 6.5-10 | 290 ± 70 | 630,000 ± 19,000 | 8.4 ± 0.1 |
| KABLE2.5 | 6.5-10 | 570 ± 20 | 2,200,000 ± 100,000 | 8.3 ± 0.1 |

[a]Error bars represent the standard errors of the mean from three independent measurements. Maximum values from fitting the pH-dependence of kinetic parameters were used.
[b]Values were from fitting pH-dependence of $k_{cat}$.

**Extended Data Table 3 | Kinetic parameters[a] of KABLE2.5 and HG3.17.**

| Protein | $k_{cat}$, $s^{-1}$ | $K_M$, mM | ($k_{cat}/K_M$), $M^{-1}s^{-1}$ |
|---|---|---|---|
| HG3.17[b] | 106 ± 17 | 0.51 ± 0.25 | 121,700 ± 40,500 |
| KABLE2.5[b] | 508 ± 23 | 0.51 ± 0.06 | 832,700 ± 86,300 |
| HG3.17[c] | 56.3 ± 9.0 | 0.91 ± 0.25 | 62,100 ± 19,700 |
| KABLE2.5[c] | 274 ± 14 | 0.37 ± 0.06 | 651,000 ± 105,600 |
| HG3.17[d] | 30.4 ± 2.7 | 0.42 ± 0.09 | 72,300 ± 15,500 |
| KABLE2.5[d] | 291 ± 13 | 0.38 ± 0.04 | 768,500 ± 95,700 |

[a]Error bars represent the standard errors of the mean from three independent measurements. Assay conditions are specified by the superscript.
[b-d]Assay conditions reported for measuring the activity of HG3.17[61].
[b]Assay conditions: 50 mM sodium phosphate, pH 7.0; 100 mM NaCl; 10% MeOH, 27 ºC.
[c]Assay conditions: 50 mM sodium phosphate, pH 7.5; 100 mM NaCl; 10% MeOH, 27 ºC.
[d]Assay conditions: 50 mM Bis-tris propane, pH 8.0; 100 mM NaCl; 10% MeOH, 27 ºC.

**Extended Data Table 4 | Kinetic parameters of reported proteins catalyzing the Kemp elimination of 5-nitrobenzisoxazole with base-mediated mechanism.**

| Protein | Description | $k_{cat}/K_M$, $M^{-1}s^{-1}$ | $k_{cat}$, $s^{-1}$ | $K_M$, mM | Amino acid efficiency[a] | Ref |
|---|---|---|---|---|---|---|
| 34E4 | Catalytic antibody | 5500 | 0.66 | 1.2 | 26 | 135 |
| BSA | Natural lipid carrier | 6500 | 6.02 | ND | 11 | 136 |
| KE07 | Computational redesign | 12.2 | 0.018 | 1.4 | 0.05 | 60 |
| KE59 | Computational redesign | 163 | 0.29 | 1.8 | 0.66 | 60 |
| KE70 | Computational redesign | 78 | 0.16 | 2.1 | 0.31 | 60 |
| KE07 R7 10/11G | 7 round DE of KE07 | 2590 | 1.37 | 0.54 | 10 | 77 |
| KE59 R13_3/11H | 13 round DE of KE59 | 60,430 | 9.53 | 0.16 | 240 | 108 |
| KE70 R8 15/11E | 8 round DE of KE70 | 34,900 | 5.3 | 0.15 | 140 | 137 |
| HG3 | Computational redesign and rational engineering | 425 | 0.68 | 1.6 | 1.4 | 63 |
| HG3.17 | 17 round DE of HG3 | 150,000 | 604 | 3.5 | 500 | 61,62 |
| HG3.R5 | 5 round DE of HG3 | 170,000 | 702 | 4.8 | 560 | 62 |
| HG4 | Ensemble-based redesign of HG3 | 103,000 | ND | ND | 340 | 66 |
| HG649 | Ensemble-based redesign of HG3 | 32,000 | ND | ND | 110 | 123 |
| AlleyCat | Single mutation of Calmodulin | 5.8 | ND | ND | 0.08 | 64 |
| AlleyCat10 | NMR-guided DE of AlleyCat7 | 4378 | 21.2 | 4.8 | 59 | 65 |
| Lysozyme L99A/M102H | Double mutation of Lysozyme | 1.8 | ND | ND | 0.01 | 138 |
| GNCA4 W229D F290W | Double mutation of GNCA4 | 5497 | 10 | ND | 21 | 67 |
| V4 | DE of GNCA4 W229D F290W | 200,000 | 635 | 3.14 | 710 | 139 |
| KE15 | Computational redesign | 35 | 0.022 | 0.63 | 0.14 | 60 |
| KE15 Tyr167Lys+ R4 | Electrical field-guided design of KE15 | 403 | 0.31 | 0.77 | 1.6 | 73 |
| tKSI | Natural enzyme | 2.5 | ND | ND | 0.02 | 68 |

| | | | | | | |
|---|---|---|---|---|---|---|
| D38N tKSI | Single mutation of tKSI | 17,000 | ND | ND | 140 | [68] |
| Des27 | Computational design | 130 | 0.07 | 0.5 | 0.52 | [71] |
| Des27.7 | Computational redesign | 12,700 | 2.85 | 0.21 | 51 | [71] |
| F113L Des27.7 | Single mutation on Des27.7 | 123,000 | 30 | ND | 490 | [71] |
| **KABLE1** | De Novo Design | **3,800** | **1.7** | **0.45** | **33** | This work |
| **KABLE1.4** | Quadruple mutant of KABLE1 | **630,000** | **290** | **0.46** | **5,400** | This work |
| **KABLE2.5** | NMR-guided DE of KABLE1.4 | **2,200,000** | **570** | **0.26** | **19,000** | This work |

[a]Amino acid catalytic efficiency, which is the quotient of catalytic efficiency ($k_{cat}/K_M$, $M^{-1}s^{-1}$), by the total number of amino acid for each protein.
