## Supplementary Material for "Emergence of specific binding and catalysis from a designed generalist binding protein"

**Affiliations:**

#### Supplementary Methods

##### ABLE expression and purification

Plasmid DNA encoding $His_6$-tagged ABLE with a Tobacco Etch Virus (TEV) protease cleavage site for tag removal (pET11a) was transformed into BL21 (DE3) *Escherichia coli*[1]. After overnight growth at 37°C (Luria-Bertani (LB)-agar, 100 µg/ml carbenicillin), a single colony was used to inoculate small-scale cultures (10 ml LB, 100 µg/ml carbenicillin). Following overnight growth at 37°C, large-scale cultures (1 L LB, 100 µg/ml carbenicillin) were inoculated and grown at 37°C until an OD600 = 0.7, at which point protein expression was induced with 0.2 mM IPTG for 4 hours at 37°C. Cells were harvested by centrifugation (4000 g, 20 min, 4°C) and cell pellets were briefly washed with phosphate-buffered saline before being re-centrifuged (4000 g, 20 min, 4°C). The cell pellets were frozen at -80°C until further purification. Thawed cell pellets were resuspended in 25 ml lysis buffer (50 mM sodium phosphate pH 7.4, 150 mM NaCl, 20 mM imidazole) and 1/2 tablet of protease inhibitor (EDTA-free complete, Roche). Cells were lysed by sonication (2 min) and cell debris was collected by centrifugation (30,000 g, 30 min, 4°C). The supernatant was filtered (0.45 µm cellulose acetate) and incubated with Ni-NTA beads (pre-washed with lysis buffer) for 4 hrs at 4°C with gentle rolling. Beads were thoroughly washed with lysis buffer before protein was eluted from the beads with elution buffer (50 mM Tris pH 8.0, 100 mM NaCl, 0.5 mM EDTA, 1 mM DTT, 300 mM imidazole). Eluted protein was supplemented with TEV protease (mass ratio: 1:10 = TEV: ABLE) and dialyzed against TEV reaction buffer (50 mM Tris pH 8.0, 100 mM NaCl, 0.5 mM EDTA, 1 mM DTT) overnight at 4°C. After tag cleavage, the protein was incubated with Ni-NTA beads (pre-washed with TEV reaction buffer) for 2 hrs and the cleaved ABLE was eluted with TEV reaction buffer. The eluted fractions were concentrated to 4 ml and further purified via HPLC (C4 reverse phase column, 30-70% acetonitrile in water). Fractions containing protein were collected and lyophilized for 72 hrs. Lyophilized ABLE was stored at -80°C.

##### Expression and purification for designed FABLE proteins

The designed FABLE proteins, with additional N-terminal His tag and TEV cleavage site, were subjected to codon optimization for *E. Coli.* using Integrated DNA Technologies (IDT) codon optimization tool. The optimized genes with additional sequences of (GTTTAACTTTAAGAAGGAGATATACAT) at 5' and (CTCGAGTGAGATCCGGCTGCTAAC) at 3' of gene were ordered as "clonal genes" from Twist Biosciences, which were assembled into pET29b vector digested with Nde1 and Xho1 using Gibbson assembly [2].

The plasmids were introduced into *Escherichia coli* BL21(DE3) cells (C2527H, New England Biolabs) via chemical transformation. The transformed cells were cultured in a 5 mL starter culture of LB medium supplemented with 50 $\mu$g ml$^{-1}$ kanamycin at a temperature of 37°C. The starter culture was used for protein expression at 1 L LB supplemented with 50 $\mu$g ml$^{-1}$ kanamycin. Protein expression was induced with 1 mM isopropyl β-D-1-thiogalactopyranoside (IPTG) (Apex Bioresearch Products) when the optical density at 600 nm ($A_{600}$) reached 0.8. After overnight expression at 30°C, the pellets were harvested by centrifugation, and their pellets were stored at -80°C.

For protein extraction, the cell pellets were resuspended in PBS buffer (119-069-13, Quality Biological) and lysed via sonication. The resulting cell lysates were clarified by centrifugation at 14,000 g for 30 min. The supernatants after centrifugation were purified with nickel-nitrilotriacetic acid (Ni-NTA) resin (Invitrogen), following the manufacturer's instruction. After buffer exchange to PBS using PD-10 column (17085101, Cytiva), proteins were analyzed using sodium dodecyl-sulfate polyacrylamide gel electrophoresis (SDS-PAGE) and fast protein liquid chromatography (FPLC).

**Screening fluorophore library with FABLE**

Fluorophores dissolved in DMSO were added to FABLE in PBS to a final concentration of 38 μM FABLE and 50 μM coumarin fluorophores, including A1 to D4, whose individual structures are summarized in Supplementary Table 3. Besides fluorophores with coumarin scaffolds, we also evaluated the interaction of FABLE with other environment-sensitive fluorophores, including L036 to TMR. Those fluorophores dissolved in DMSO were added to FABLE in PBS to a final concentration of 38 μM FABLE and 1 μM fluorophore. The fluorescence of each fluorephore was recorded with BioTek Synergy Neo2 Reader after incubating at room temperature for 30 minutes. The exact wavelength of excitation and emission for these fluorophores are in Supplementary Table 3.

**FPLC analysis**

To confirm the coumarin dyes were firmly bound to the FABLE, we prepared the FABLE-coumain complexes and showed that the complexes were stable during chromatographic separation. To prepare the ligand-FABLE complex, 100 μM FABLE was mixed with 200 μM of Cou485, Cou481, and Cou540a, respectively. After incubation at room temperature for 30 min, the resulting complex was injected to the Fast Protein Liquid Chromatography (FPLC) equipped with a Superdex 75 Increase 5/150 analytical column. The ligand was monitored by their absorbance indicated in the in Supplementary Fig. 9.

**Determining binding constants**

We used fluorescence titration experiments to determine the binding dissociation constants of FABLE for Cou485, Cou481, and Cou540a with BioTek Synergy Neo2 Reader.
The excitation and emission wavelength with the largest shift upon FABLE interaction was used to quantify the the bound ligand, which is 395 nm (excitation) and 470 nm (emission) for Cou485, 410 nm (excitation) and 470 nm (emission) for Cou481, 440 nm (excitation) and 510 nm (emission) for Cou540a. For each ligand, FABLE in PBS was serially diluted while fixing the concentration of ligand at indicated values. Global fitting with 2 fixed ligand concentrations was used to fit $K_D$ following the single-site binding model (Equation 1, N=1, L=indicated concentration, shared $K_D$) using GraphPad Prism. $b_{free}$ was obtained by measuring the fluorescence of the ligand in PBS after serial dilution.

**Equation 1**

$$Fluorescence = b_{bound}[Ligand_{bound}] + b_{free}[Ligand_{free}]$$

- $Ligand_{total:}$ total concentration of ligand
- $Ligand_{free:}$ free ligand
- $Ligand_{bound}$: ligand binding to protein
- $Ligand_{free} = ligand_{total} - ligand_{bound}$
- $b_{free} =$ brightness (fluorescence) of free ligand per μM
- $b_{bound} =$ brightness (fluorescence) of bound ligand per μM

**Equation 2**

$$\text{bound ligand} = \frac{1}{2}\left[K_D + [Ligand_{\text{total}}] + \frac{[Protein_{\text{total}}]}{N} - \sqrt{\left(K_D + [Ligand_{\text{total}}] + \frac{[Protein_{\text{total}}]}{N}\right)^2 - 4\,[Ligand_{\text{total}}]\frac{[Protein_{\text{total}}]}{N}}\right]$$

- $Protein_{total}$: total protein concentration
- N: binding stoichiometry
- $K_D$: binding constant of protein-ligand interation

**FABLE crystallization and structure determination**

FABLE crystals were grown by sitting drop vapor diffusion using SwissCI 3-well crystallization plates with reservoir solutions containing: i) 100 mM Tris pH 8.0, 20% (v/v) MPD, ii) 200 mM sodium chloride, 100 mM CHES pH 9.5, 50% (v/v) PEG 400 or iii) 100 mM CHES pH 9.5, 30% (v/v) PEG 600. Crystal drops were set up with 200 nl reservoir and 200 nl protein. Crystals were looped and vitrified in liquid nitrogen without further cryoprotection. X-ray diffraction data were collected at SSRL beamline 12-2 (MPD condition) or ALS beamline 8.3.1 (PEG 400 and PEG 600 condition) with the strategy and data collection statistics summarized in Supplementary File 1. Data were reduced and models refined using the same procedure as described for ABLE, except phases were obtained for the MPD structure using molecular replacement and the previously published ABLE structure (PDB code 6W6X), while phases for the PEG 400 and PEG 600 structures were obtained using the structure from the MPD condition (PDB code 9DWA). Refinement statistics are reported in Supplementary File 1 and coordinates and structure factor amplitudes have been deposited in the PDB with accession codes 9DWA (MPD condition), 9DWB (PEG 400 condition) and 9DWC (PEG 600 condition).

**Thermal stability**

Proteins were prepared in 10 μM concentrations in PBS buffer. CD spectra were collected on a Jasco J-810 CD spectrometer in a 0.1 cm path-length quartz cuvette. The parameters for full spectra collections were set up as follows: the bandwidth was set to 2 nm, the scanning speed was set to 50 nm/min, and the average of accumulations was set to 3. Temperature-dependent data were collected at 222 nm from 20 to 95 °C with an interval of 5 °C and an increase rate of 2 °C/minute.

**Library design**

The plasmid encoding the KABLE1 gene was used as the template for site-specific saturation mutagenesis, which was achieved using megaprimer PCR protocol with primer sets (Integrated DNA Technologies) that overlapped the 5' position of the randomized position, together with a flanking primer (T7 reverse). Saturation mutagenesis was performed using NNK codons (where N can be any base and K represents a mixture of G and T nucleotides) that cover the 20 genetically encoded amino acids (primer sequences were shown in Supplementary Table 9). There are 20 libraries with NNK codon at indicated residue. The size of the PCR product was checked using agarose gel (1%) electrophoresis. Product DNA was digested with DpnI (New England Biolabs) at 25 °C for 12-14 hours to eliminate the parental clone. The sample was transformed into *E. coli* NEB5α cells (C2987H, New England Biolabs) and plated on an LB agar plate containing kanamycin (50 μg $ml^{-1}$). Colonies obtained from the plate upon incubation at 37 °C for 12-14 hours were grown in LB media with kanamycin at 37 °C for 2-3 hours. Cells were collected by centrifugation at 6,500*g* for 10 min. Plasmids were extracted using a DNA plasmid extraction kit following the manufacturer's protocol (Monarch, New England Biolabs). Library quality was confirmed by Sanger sequencing (Azenta Life Sciences).

**Library screening**

Around 160 randomly picked independent colonies were screened for each library. The plasmid encoding the NNK library was transformed into *E. coli* BL21 cells and plated on LB agar containing kanamycin. Individual colonies were inoculated in LB media (200 µl) containing kanamycin in a 96-well plate. Cultures were incubated at 37 °C, 220 rpm for 5-6 hours. A replica deep 96-well plate (Costar) was generated, where cultures (20 µl) were inoculated in LB media (400 µl) containing kanamycin and IPTG (0.5 mM). Cultures were then allowed to grow at 30 °C, 220 rpm for 20 h. Cells were collected by centrifugation at 4 °C, 2,000*g*. Pellets were resuspended in buffer A (25 mM Tris, pH 8.0, 20 mM imidazole, 300 mM NaCl), centrifuged again, and the supernatant was discarded. Cells were lysed using a buffer containing 25 mM Tris, pH 8.0, 100 mM NaCl, 0.5% triton X and centrifuged to separate the supernatant. Kemp eliminase activity of the clones was tested using a 96-well plate in the buffer (20 mM Tris, pH 8.0, 100 mM NaCl) containing 1.5% acetonitrile. Reactions were initiated upon mixing cell lysate with 100 µM substrate in a 1:1 ratio (final substrate concentration of 50 µM). Product formation was observed by measuring absorbance at 380 nm at 25 °C on a plate reader for 10 min. The activities of the clones showing notable improvement in activity over the initial design were confirmed by rescreening them in triplicate. Plasmids extracted from the colonies demonstrating enhanced activity were sequenced (Azenta Life Sciences) to determine the identities of beneficial mutations. Sequences of the design (KABLE1) as well as its mutants (KABLE1.4 and KABLE2.5) are provided in Supplementary Table 10.

**Expression and purification of KABLEs**

The plasmids encoding KABLE0, KABLE1, and its mutants, cloned into pET29b(+) vector with NdeI and XhoI restriction sites, were transformed into *E. coli* BL21(DE3) (C2527H, New England Biolabs) and plated on LB agar plate with 50 µg $ml^{-1}$ kanamycin (Thermo Scientific). This same kanamycin concentration was used throughout the experiment. Single colonies were inoculated into 15 mL LB medium containing kanamycin and grown at 37 °C, 220 rpm for 5-6 hours. 10 ml of starter culture was then diluted into 1 L of LB with kanamycin and allowed to grow at 37 °C until $A_{600}$ reached 0.6-0.8. The culture was induced upon the addition of 0.5 mM isopropyl-ß-D-1-thiogalactopyranoside (IPTG) (Apex Bioresearch Products) and grown at 30 °C for 20 hours. Cells were harvested by centrifugation (Sorvall Legend RT+, Thermo Scientific) at 4 °C, 4, 3500 rpm for 15 min., flash frozen in liquid nitrogen, and stored at -80 °C.

For protein purification, 200 mL cells were resuspended in 40 mL buffer A (25 mM Tris, pH 8.0, 20 mM imidazole, 300 mM NaCl). Cell lysis was done upon sonication (Sonic Dismembrator Model 500, Fisher Scientific) for 10 min. (20 s pulse, 20 s rest) at an amplitude of 30%. To isolate the soluble fraction, centrifugation was performed at 4 °C, 20,000*g* for 30 min. The lysate was loaded onto a Ni-NTA gravity column (Hispur, Thermo Scientific) pre-equilibrated with buffer A, washed, and eluted using buffer B (25 mM Tris, pH 8.0, 250 mM imidazole, 300 mM NaCl). Protein fractions identified by Bradford assay were combined, concentrated to 3 ml using 10K MWCO spin concentrator (Amicon Ultra, Millipore Sigma), and exchanged into TEV cleavage buffer (50 mM Tris, pH 8.0, 75 mM NaCl) using desalting column (BioRad, 10 DG). TEV protease (at protein-to-protease absorbance ($A_{280}$) ratio of 100:1) along with EDTA (0.5 mM) and DTT (1 mM) was added to cleave the N-terminal $His_6$-tag. Protein was incubated at 37 °C for 12-14 hours and exchanged into buffer C (20 mM HEPES, pH 7.0, 100 mM NaCl). Protein fractions were then passed through Ni-NTA column to remove the protease and any un-cleaved KABLE protein. Final purification was performed on GE Healthcare AKTA FPLC system with Superdex 75 Increase 10/300 column in buffer C. Protein concentrations were determined by measuring the absorbance at 280 nm using extinction coefficient determined by Expasy ProtParam tool (https://web.expasy.org/protparam/).

**Kinetic characterization of KABLEs**

The initial rates for Kemp elimination for all the kinetic assays were corrected for the background rates in the appropriate buffers without the enzyme. Product (2-hydroxy-5-nitro-benzonitrile) concentration was determined using an extinction coefficient of 15,800 $M^{-1}cm^{-1}$ at 380 nm. Substrate 5-nitrobenzisoxazole (5NBI, AK Scientific) concentration ranging from 100 µM to 1 mM were used. The substrate 5NBI was prepared in acetonitrile as 100 mM stock, which was further diluted in buffer to prepare working solutions of various substrate concentrations. The reactions were initiated by mixing the enzyme in buffer and solutions of 5NBI in a 1:3 ratio in a 96-well plate (Genesee Scientific). The final reaction mixtures contained enzyme (1-1000 nM, depending on the activity of the mutant, Supplementary Table 4) in buffer, 1.5% acetonitrile (to maintain the constant final concentrations of the substrate), and variable concentrations of substrate. To maintain the pH at different values, different buffers[1] were used at 20 mM in the presence of 100 mM NaCl (MES at pH 6.0 and 6.5, HEPES at pH 7.0 and 7.5, Tris at pH 8.0 and 8.5, CHES at pH 9.0 and 9.5, and CAPS at pH 10.0). Product formation was monitored on a plate reader (BioTek Synergy neo2, Serial: 22052011) at 25 °C for 10 min. with the minimum kinetic interval determined by manufacturer provided software (Gen 5, version 3.12). The initial rates, under a series of substrate concentrations in a specified buffer, were used to fit the $k_{cat}$ and $K_M$ using the Michaelis-Menton equation in GraphPad Prism. The dependence of $k_{cat}/K_M$ and $k_{cat}$ on pH was fitted using the equation $k_{\mathrm{cat}}/K_{\mathrm{M}} = (k_{\mathrm{cat}}/K_{\mathrm{M}})_{\mathrm{EH}} \times (1-\mathrm{f}_{\mathrm{E-}}) + (k_{\mathrm{cat}}/K_{\mathrm{M}})_{\mathrm{E-}} \times \mathrm{f}_{\mathrm{E-}}$, in which $\mathrm{f}_{\mathrm{E-}}$ is the fraction of the protein in the anionic state and is related to pH and the p$K_a$ of its base by: $\mathrm{f}_{\mathrm{E-}} = 1/(1+10^{\mathrm{pKa-pH}})$. The $(k_{\mathrm{cat}}/K_{\mathrm{M}})_{\mathrm{E-}}$ was used as the maximum $k_{\mathrm{cat}}/K_{\mathrm{M}}$. Similarly, $k_{\mathrm{cat}} = (k_{\mathrm{cat}})_{\mathrm{EH}} \times (1-\mathrm{f}_{\mathrm{E-}}) + (k_{\mathrm{cat}})_{\mathrm{E-}} \times \mathrm{f}_{\mathrm{E-}}$.

**KABLE crystallization and structure determination**

LigandMPNN[1] chose very polar and flexible surface residues for KABLE, which impeded crystallization. To improve the crystallization of KABLE1.2, we introduced 23 surface mutations to make KABLE12$_{\mathrm{cryst}}$. Because ABLE crystallizes easily, we transplanted its surface residues to KABLE12$_{\mathrm{cryst}}$ as indicated in Supplementary Fig. 19. Following the previous expression and purification procedures, both the histag-cleaved version and histag-uncleaved version at PBS buffer were used to screen crystals and prepare the complex with 6NBT. 100 mM 6NBT dissolved in DMSO was added to 2.5 mM KABLE12$_{\mathrm{cryst}}$ for the final 6NBT concentration of 10 mM.
Apo crystals of KABLE12$_{\mathrm{cryst}}$ with an intact His-tag were grown by sitting-drop vapor diffusion using SwissCI 2-well crystallization plates with reservoir solution containing 0.2 M sodium bromide, 0.1 M Bis-Tris propane pH 6.5, 20% PEG 3350. Crystals of KABLE12$_{\mathrm{cryst}}$ in complex with 6NBT grew using His-tag cleaved protein from a reservoir containing 0.2 M zinc acetate, 0.1 M sodium cacodylate pH 6.5, 18% PEG 8000. Crystals were looped and vitrified in liquid nitrogen without further cryoprotection. X-ray diffraction data were collected at ALS beamline 8.3.1 with the strategy and data collection statistics summarized in Supplementary File 1. Data were reduced and models refined using the same procedure as described for ABLE. For the structure in complex with 6NBT, ligand restraints were generated with *phenix.elbow* (for 6NBT) and Grade2 (for

cacodylate) -. In addition to the 6NBT molecule present in the KABLE core, two additional 6NBT molecules were identified in complex with three zinc molecules and a cacodylate ion at a crystal packing interface (Supplementary Fig. 20). The metal identity was supported by the calculation of an anomalous difference density map showing large peaks for the zinc atoms and no peak for the cacodylate arsenic (Supplementary Fig. 20). B-factors for the zinc atoms were refined anisotropically to account for the observed disorder (Supplementary Fig. 20).

**Expression of $^{15}N$ and $^{13}C$-labelled Proteins**

For the expression of isotopically labelled KABLE variants, the plasmid was transformed into *E. coli* BL21(DE3) cells and plated on LB agar plate containing kanamycin (50 µg/ml). A single colony was inoculated with 2 ml of LB medium with kanamycin of same concentration mentioned above. The culture was grown at 37 ºC for 5-6 h, then transferred to 18 ml of unlabelled M9 minimal medium with kanamycin and grown further at 37 ºC for an additional 5-6 h. The resulting 20 ml starter culture was diluted with 1 l of M9 medium made with $^{15}NH_4Cl$ as $^{15}N$ and [$^{13}C_6$]-glucose (Cambridge Isotope Laboratories), supplemented with kanamycin, and grown at 37 ºC until $A_{600}$ reached 0.6-0.8. The culture was then induced by 0.5 mM IPTG and grown at 30 ºC for 20 h. The cells were harvested by centrifugation and the isotopically labelled protein was purified as discussed above.

**Nuclear Magnetic Resonance (NMR) spectroscopy**

The NMR samples contained 1.5-1.6 mM of U-[$^{13}C$,$^{15}N$] KABLE1.4 in 20 mM HEPES 100 mM NaCl pH 6.9, 0.02% $NaN_3$ and 10% $D_2O$ for the lock. All NMR spectra were acquired at 298 K on a Bruker Avance III HD 800 MHz spectrometer equipped with a TCI cryoprobe. The assignments of KABLE1.4 backbone and CβHβ resonances in the free, inhibitor-bound, and $CH_3CN$ reference forms were obtained from a standard set of 3D BEST HNCACB, HN(CO)CACB, HNCO, HN(CA)CO, and HBHA(CO)NH experiments. For the reference (containing 7.4% v/v of $CH_3CN$) and inhibitor-bound (containing 5 molar equivalents of 6-NBT and 7.4% v/v of $CH_3CN$) forms, the assignments were extended to the side-chain methyl resonances using the 3D (H)CCH TOCSY experiment. The NMR data were processed in TopSpin 3.7 (Bruker) or NMRPipe[6], and analyzed in CCPNMR[7]. The assigned $^{1}H$, $^{13}C$ and $^{15}N$ chemical shifts have been deposited in the Biological Magnetic Resonance Bank (http://www.bmrb.wisc.edu/) under the accession number XXX.

The chemical shift perturbations (CSPs) were obtained by subtracting the peak positions in the $CH_3CN$ reference spectra from the corresponding resonances of the 6-NBT bound sample. For the backbone amides, the average CSP ($\Delta\delta_{avg}$) was calculated as $\Delta\delta_{avg} = (\Delta\delta_X^2/n + \Delta\delta_H^2/2)^{0.5}$, where $\Delta\delta_X$ and $\Delta\delta_H$ are the binding shifts of nitrogens or carbons ($\Delta\delta_X$) and protons ($\Delta\delta_H$), and n = 50 or 9 for the nitrogen or carbon atoms, respectively. For each observed resonance, the Z score was calculated as $Z = (\Delta\delta_{avg} - \mu)/\sigma$, where μ and σ are, respectively, the average and the standard deviation of $\Delta\delta_{avg}$ values for a given backbone amide.

**Landscape generation (Fig. 6a)**

To generate a conceptual landscape representing the sequence-chemical space exploration in this work, we used two axes on the horizontal level to represent protein sequence diversity and small molecule chemical diversity. The exact coordinates for the protein sequence (FABLE, KABLE) and chemical diversity (fragments, cou485, and 6NBT) were manually assigned to represent their differences relative to ABLE and apixaban. The heights of the four peaks denote the fitness for binding or catalyzing the Kemp elimination reaction, calculated by applying a Gaussian function to four sets of horizontal coordinates. Parameters of the individual Gaussian functions were chosen to reflect their experimental observations, with the amplitude representing the efficiency of activity and sigma representing the specificity of the primary function. For example, a lower and broader peak was used for the ABLE-fragments interaction, considering their weak and promiscuous interaction. The script used to generate the landscape is included as a supplemental note (generate_landscape.py), which contains all the above information.

**Alignment of four-helical bundles and GPCRs (Fig. 6b-c)**

For GPCRs in Fig. 6b, its PDB strctures were firstly cleaned by only keeping the first chain of GPCR. All the other cleaned GPCR structures were aligned to 2RH1 using TMalign. The terminal flexible regions of 6EM9 (residue 31-37 and 2001-2106) and 7WC4 (residue 67-74) are hidden to highlight the shared region of all these GPCRs.

For four-helical bundles in Fig. 6c, chain A of deposited PDB structures was used except for cou485-binder and NDI-binder, where only apo structures were deposited. For these two structures, their design structures, in good agreement with the corresponding apo structure, were used to indicate the designed ligand binding site. All the structures are aligned to 5TGY using TMalign, except for 7JH6. To accommodate the larger size and double ligand binding sites of 7JH6, the backbones atoms of its axial histidine residue and second shell threonine residue were used to superposition with the corresponding atom of 5TGY to generate the alignment. The extended structure of 7JH6 (residue 4-11, 80-98,167-177) and 8D9P (residue 2-5,24-73,127-171) are hidden to highlight the shared region of all of these four-helical bundles.

### Supplementary Figures

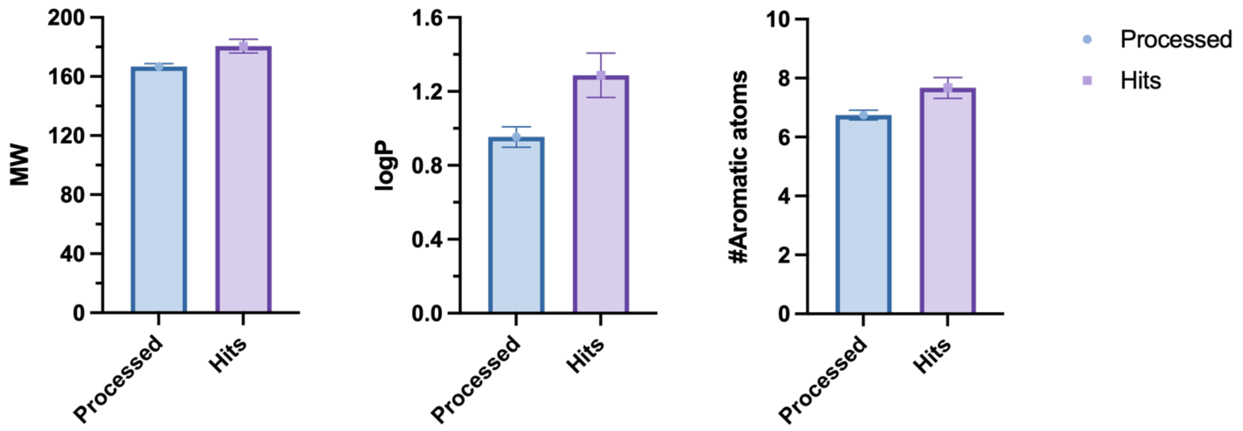


**Supplementary Figure 1 | Chemical properties of fragments in the processed diffraction dataset and hits binding at apixaban-binding pocket of ABLE.** For each chemical, its MW, logP and the number of aromatic atoms were generated using "calculate properties" function of DataWarrior. The error bars represent standard error of mean. Processed: n=242; Hits: n=39. Z scores are all over 3.

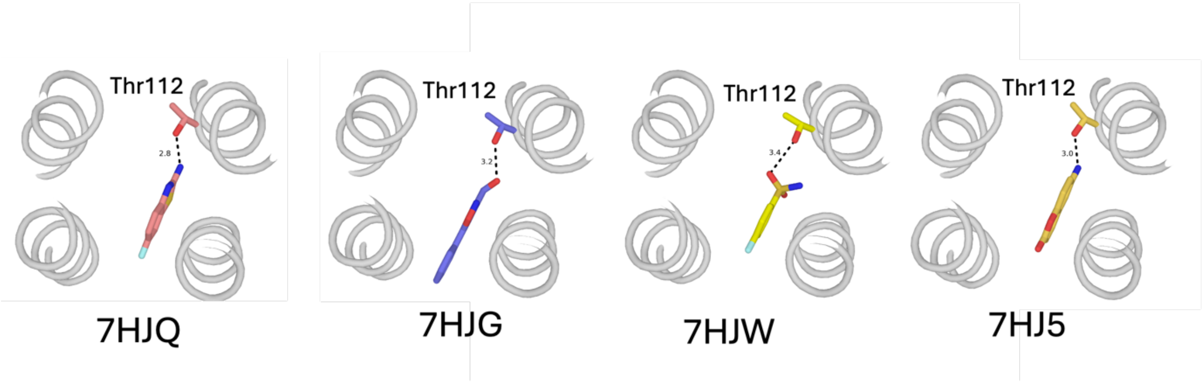


**Supplementary Figure 2 | Thr112-mediated polar interactions in many fragments at site A**. Their PDB codes are labeled at the bottom.

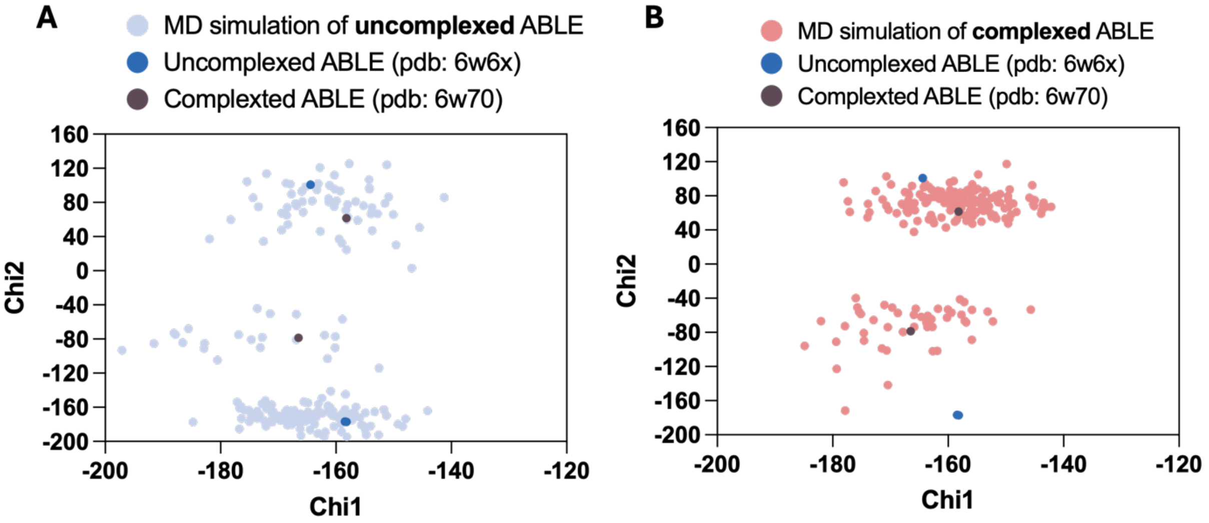


**Supplementary Figure 3 | Contour map of His49 side chain conformation of ABLE from molecular dynamics simulation.** (**A**-**B**) Dynamics of uncomplexed ABLE (PDB: 6W6X, **A**) and complexed ABLE (PDB: 6W70, **B**) were explored by doing molecular simulation at 278 K for 500 ns with amber. Chi1 and Chi2 of His49 from each state of these simulations and reported crystal structures of ABLE were extracted for plotting.

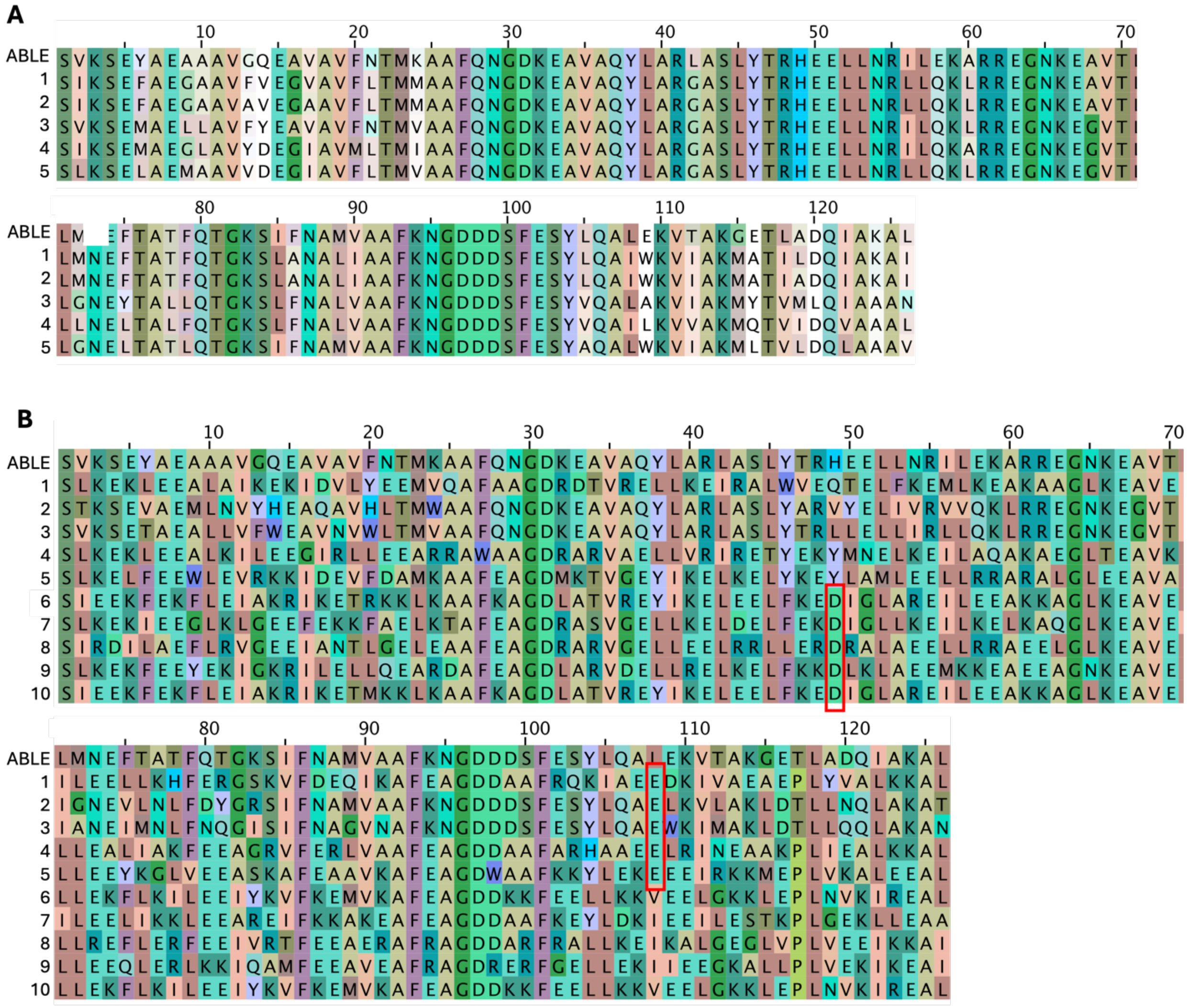

**Supplementary Figure 4 | Sequence comparison between ABLE and FABLEs/KABLEs. (A)** Amino acid sequence of all 5 designed fluorescent ABLE (FABLE$_{D1}$ to FABLE$_{D5}$), with ABLE shown on the top. FABLE$_{D1}$ wss denoted as FABLE. (**B**) Amino acid sequence of all 10 Kemp eliminase ABLE designs (KABLE$_{D1}$ to KABLE$_{D10}$), with ABLE shown on the top. The designed base Glu108 in the first strategy (KABLE$_{D1}$ to KABLE$_{D5}$) and Asp49 in the second strategy (KABLE$_{D6}$ to KABLE$_{D10}$) are highlighted in the red box. KABLE$_{D5}$ and KABLE$_{D9}$ were denoted as KABLE0 and KABLE1, respectively.

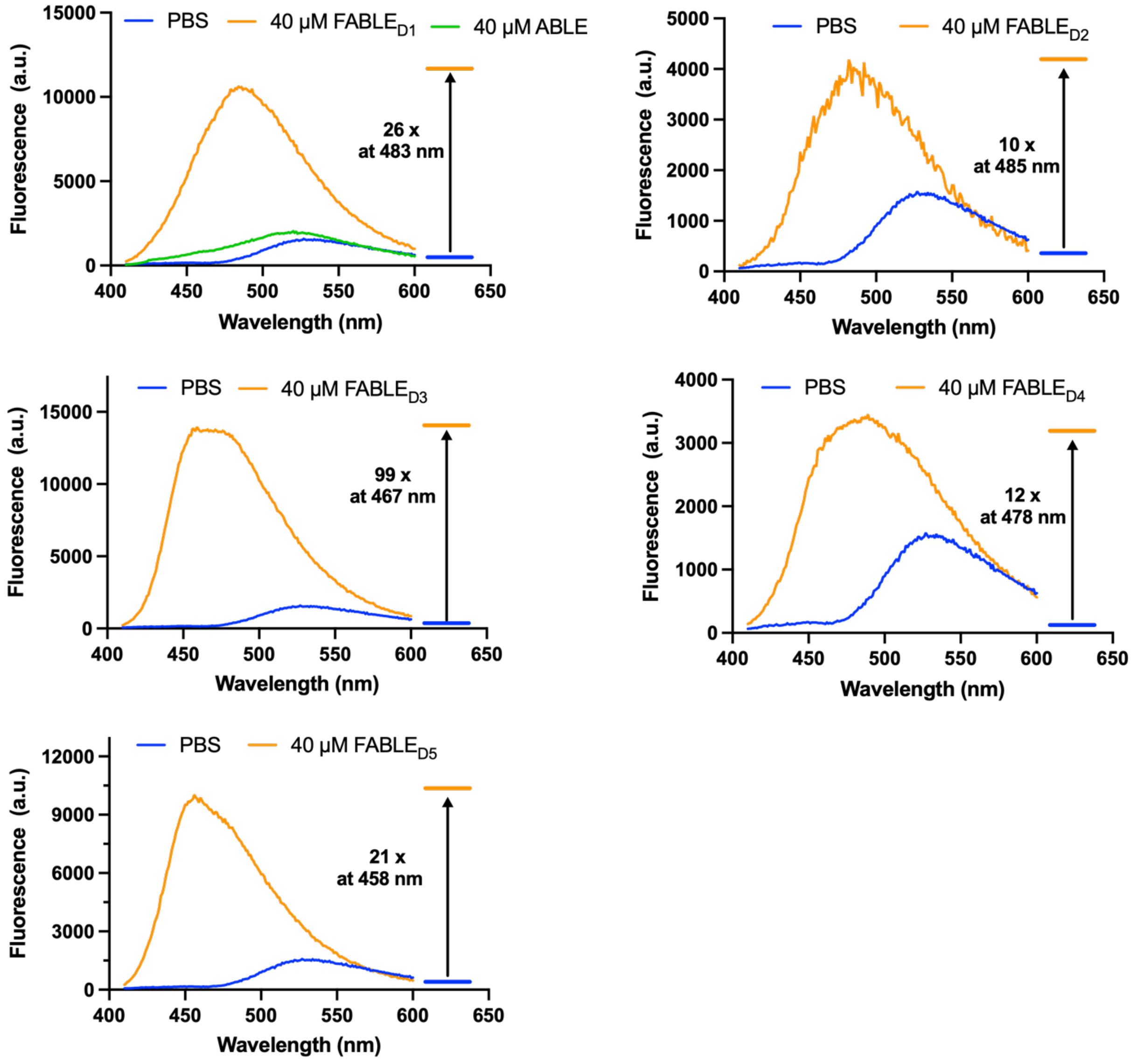


**Supplementary Figure 5 | Emission Spectra of Cou485 in the presence of FABLE protein derivatives.** The fluorescence emission spectrum of 6 µM Cou485 was recorded using BioTek Synergy Neo2 Reader with an excitation of 395 nm. The indicated FABLEs or ABLE (control) were present at 40 µM.

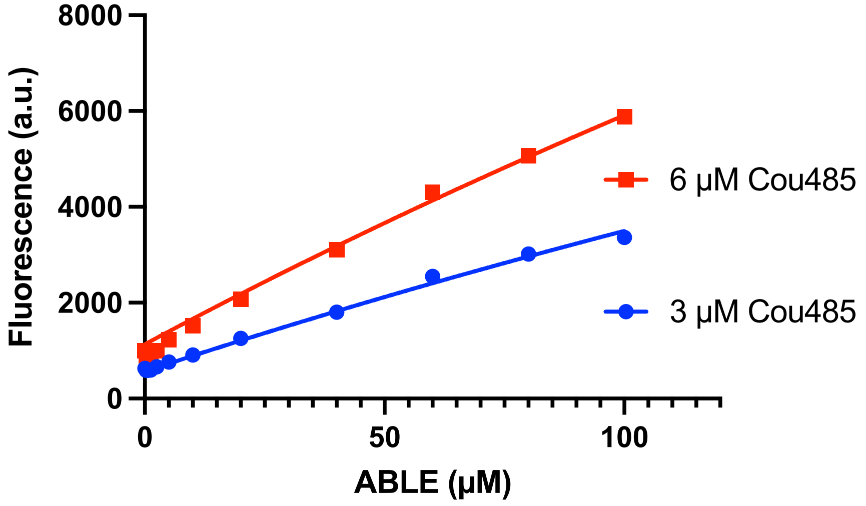


**Supplementary Figure 6 | : Fitting Cou485 fluorescence at varying concentrations of ABLE did not yield $K_D$ < 500 uM.** Unlike the quadratic binding curves observed with varying amounts of FABLEs, the fluorescence of Cou485 increased linearly with increasing concentrations of ABLE.

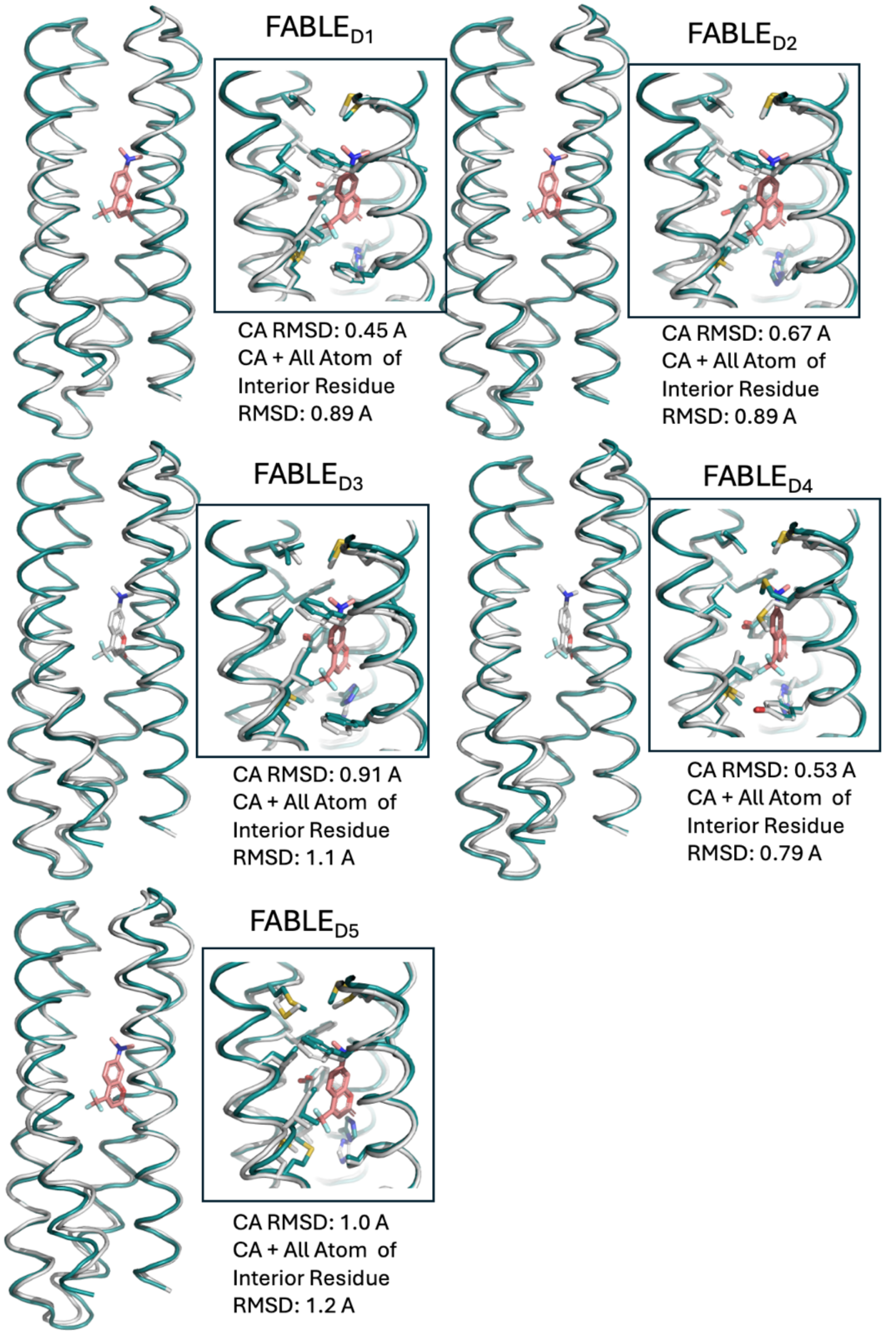


**Supplementary Figure 7 | AF3 prediction agrees with all 5 FABLE designs.** Left: Designed structure (protein in teal cartoon and ligand in pink stick) and predicted structure (protein in grey cartoon). Right: Zoom in view of active site with carbon in designed structures shown as the teal stick and carbon in predicted structures shown as a grey stick.

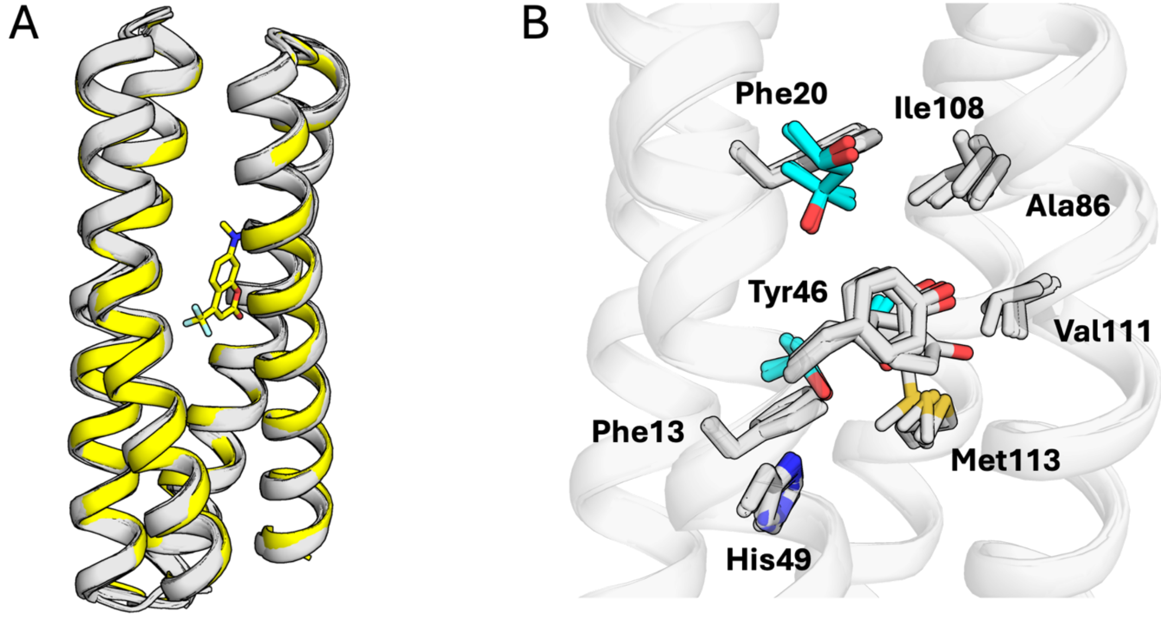


**Supplementary Figure 8 | The crystal structure of apo FABLE agrees with the designed structure. (A)** Overlay of designed structure (yellow carbon) with structures from the individual asymmetric unit in the three crystal structures (9DWA, 9DWB, 9DWC) of FABLE (grey carbon). The Cou485 in the designed structure are shown in yellow sticks. The low solubility of Cou485 under crystallization conditions limited our ability to obtain a co-crystal structure. However, the un-complexed structures are in excellent agreement with the model of the complex (with sub-Å of Cα RMSD, Supplementary Table 2). (B) Active site view of structures from the individual asymmetric units in three crystal structures (9DWA, 9DWB, 9DWC) of FABLE. The sidechains of active site residues are shown in sticks. Two methylpentanediol molecules from the crystallization buffer, which occupy the binding site in 9DWA, are shown in blue carbons. Most residues are very well organized, including Met23. Tyr46 shows some variability in the different crystal environments.

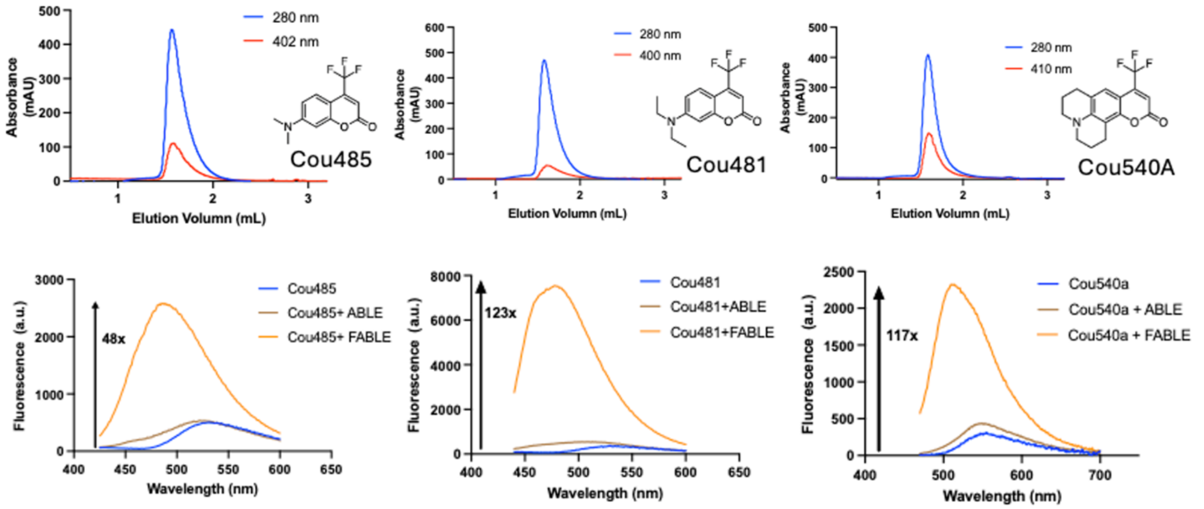


**Supplementary Figure 9 | FABLE interacts with Cou485, Cou481 and Cou540a as evidenced by SEC and an increase in fluorescence intensity by 48-123 fold.** SEC analysis of 100 µM FABLE mixed with 200 µM indicated coumarin revealed a monomeric peak for FABLE, with the protein coeluting with the coumarin at the same time. Incubation of 1.5 µM Cou485 and Cou481 with 40 µM FABLE increased fluorescence intensity at 470 nm by 48 and 122 fold, respectively. Similarly, incubation of 0.5 µM Cou540a with 6 µM FABLE resulted in a 117-fold increase in fluorescence at 510 nm.

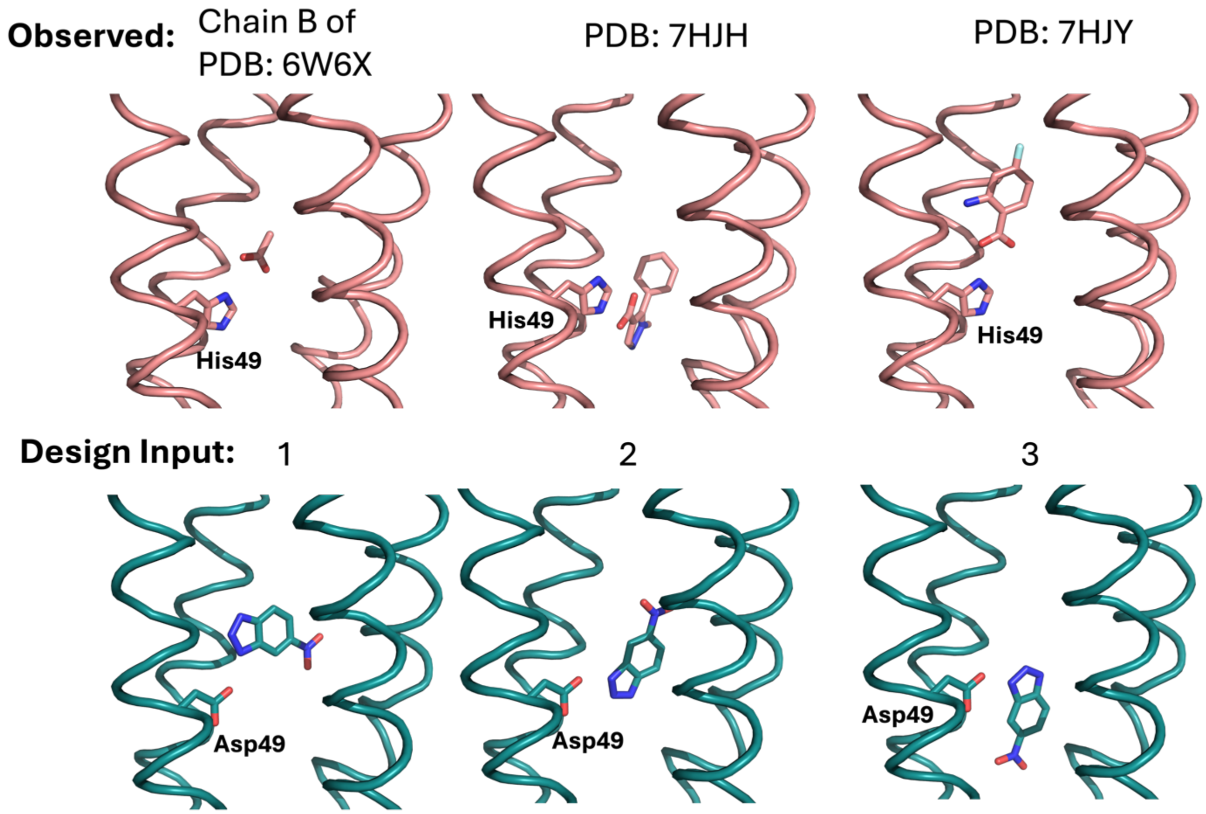


**Supplementary Figure 10 | Observed His49-carboxylic interactions in ABLE-fragment complexes and the Asp49-6NBI interactions used as starting points to design KABLE1.** Top: Examples of observed ABLE-fragment interactions with indicated PDB code, carbon of His49 and fragments are shown in pink sticks. Bottom: three initial input structure for designing KABLE1 with each utilizing one distinctive lone pair of electron on either OD1 or OD2 of Asp49 (see see section 1.11.2 of SI). Carbon of Asp49 and fragments are shown in teal sticks. Similar observed orientations of residue 49 and fragment are used as input to design KABLE1.

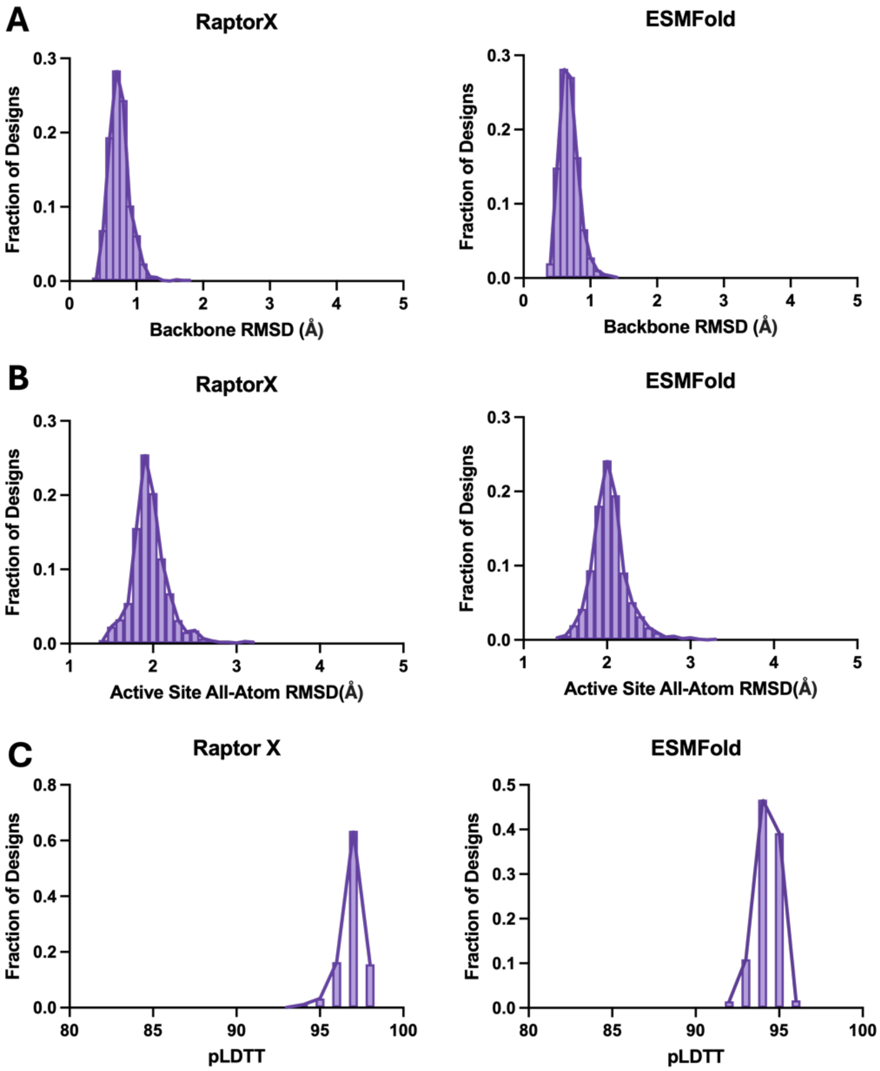


**Supplementary Figure 11 | Structure prediction metrics of KABLE designs bearing Asp49.** All 1000 sequences were subjected to structure prediction with both RaptorX and ESMFold. (**A**) Backbone RMSD between either prediction and design shows most of designs are within 1 Å backbone RMSD. (**B**) All-atom RMSD of active site residues (< 3.5 Å of TSA) between either prediction and design shows the median of active site RMSD at 2 Å. (**C**) pLDTT scores from both prediction methods show all the designs are with pLDTT score over 90.

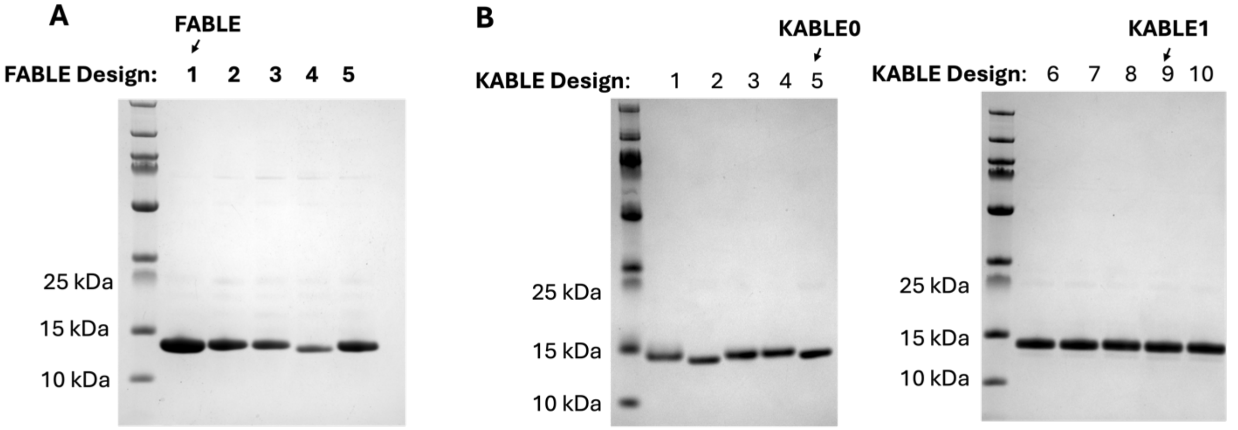


**Supplementary Figure 12 | SDS-PAGE Characterization of all designed proteins**. **(A)** SDS-PAGE analysis of the purified form of all 5 FABLE designs ($FABLE_{D1}$-$FABLE_{D5}$), with the $FABLE_{D1}$ designated as FABLE. **(B)** SDS-PAGE analysis of all 10 KABLE designs from the first design strategy of aromatic box ($KABLE_{D1}$-$KABLE_{D5}$) and the second design strategy of versatile residue 49 ($KABLE_{D6}$-$KABLE_{D10}$), with the $KABLE_{D5}$ highlighted as KABLE0 and $KABLE_{D9}$ highlighted as KABLE1.

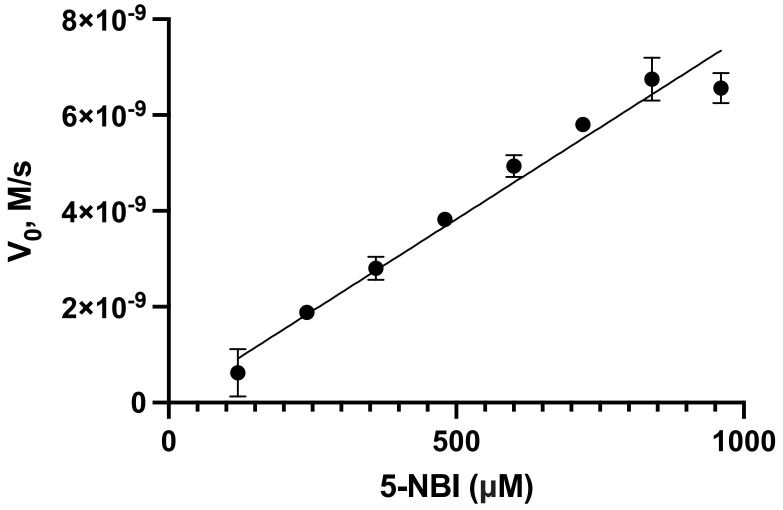


**Supplementary Figure 13 | Substrate concentration dependence versus rate data for KABLE0.** The reaction was measured with 1 μM KABLE0 in pH 8 buffer. Background reaction at the same condition without enzyme was subtracted and data were analyzed by least squares analysis to the linear portion of the Michaelis-Menten model ($v_0 = (k_{cat}/K_M)[E_0][S]$), and $k_{cat}/K_M$ was deduced from the slope. Error bar stands for the standard deviation.

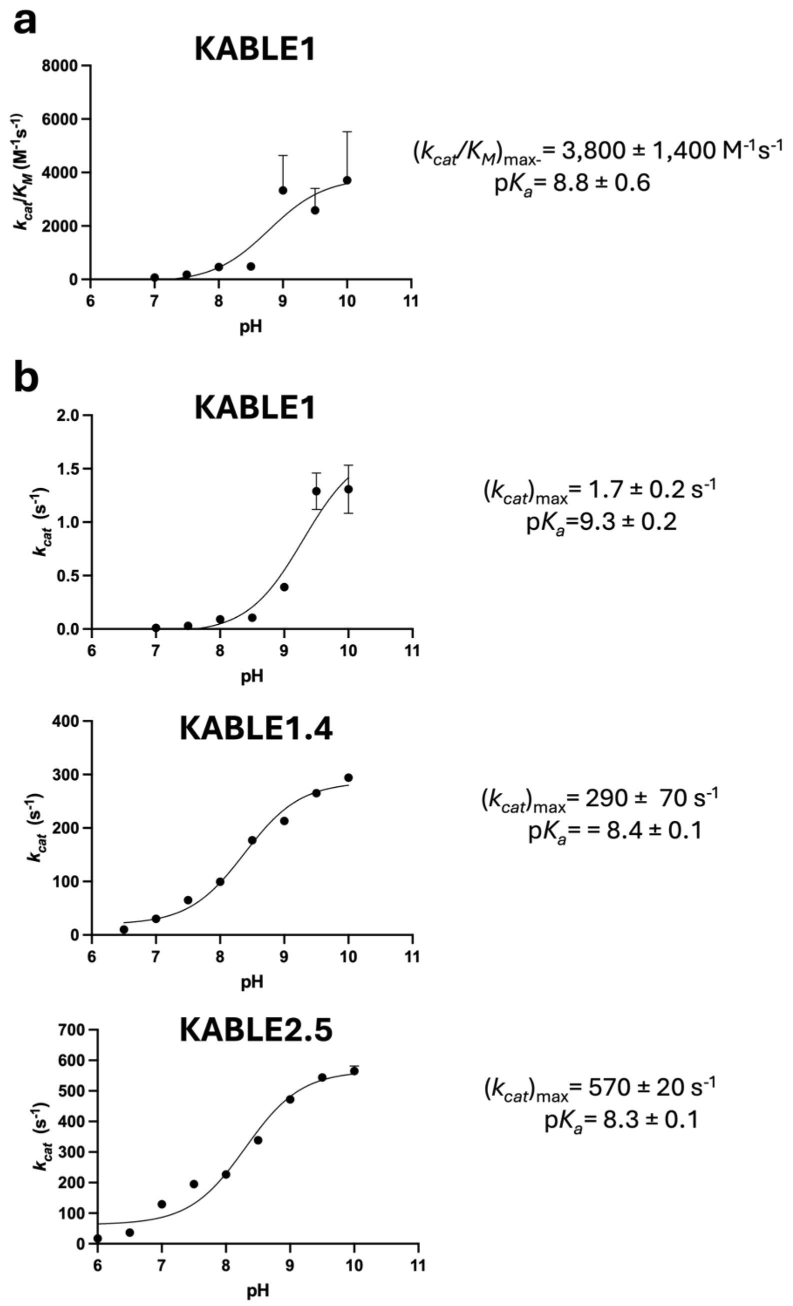


**Supplementary Figure 14 | Dependence of $k_{cat}/K_M$ and $k_{cat}$ on pH.** (a) pH-dependence of $k_{cat}/K_M$ for KABLE1. (b) pH-dependence of $k_{cat}/K_M$ for KABLE1, KABLE1.4, and KABLE2.5, showing a decreasing trend in p$K_a$ as the activity improves.

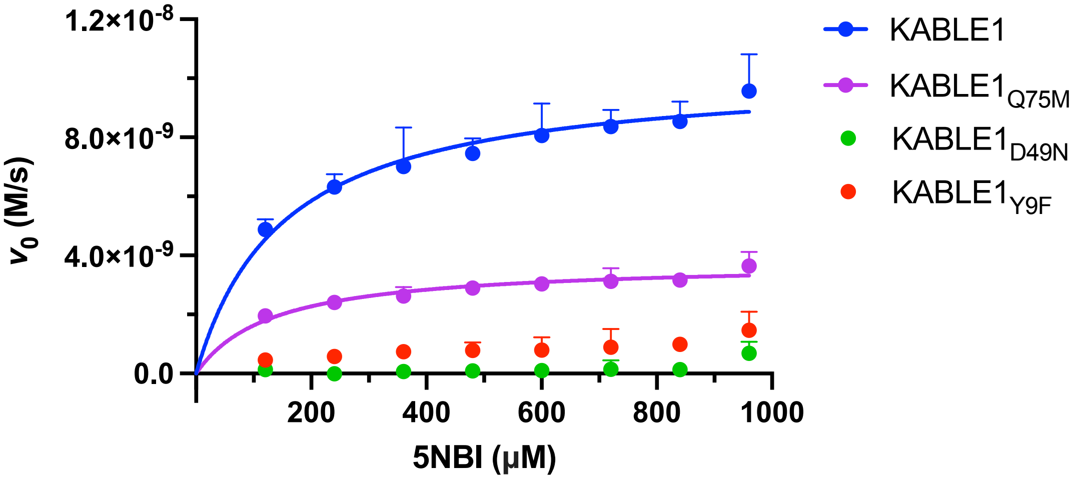


**Supplementary Figure 15 | Substrate concentration versus rate data for KABLE1 mutants at Q75 and Y9 and D49.** Reactions were measured at pH 7.0 buffer (20 mM HEPES,100 mM NaCl) with 1 μM KABLE1, 2.5 μM $KABLE1_{Q75M}$, 2.5 μM $KABLE1_{Y9F}$ and 2.5 μM $KABLE1_{D49N}$, respectively. Fitting the data with Michaelis-Menten model yielded KABLE1 ($k_{cat}/K_M$ = 68 ± 28 $M^{-1}$ $s^{-1}$, $k_{cat}$ = 1.0 ± 0.10 x$10^{-2}$ $s^{-1}$, $K_M$ = 0.15 ± 0.023 $mM^{-1}$) and $KABLE1_{Q75M}$ ($k_{cat}/K_M$ = 11 +7.3 $M^{-1}$ $s^{-1}$, $k_{cat}$ = 0.15 ± 0.013 x$10^{-2}$ $s^{-1}$, $K_M$ = 0.13 ± 0.083 $mM^{-1}$). For $KABLE1_{Q75M}$ and $KABLE1_{D49N}$, the fitting was not able to return reliable parameters. Error bar represents standard deviation.

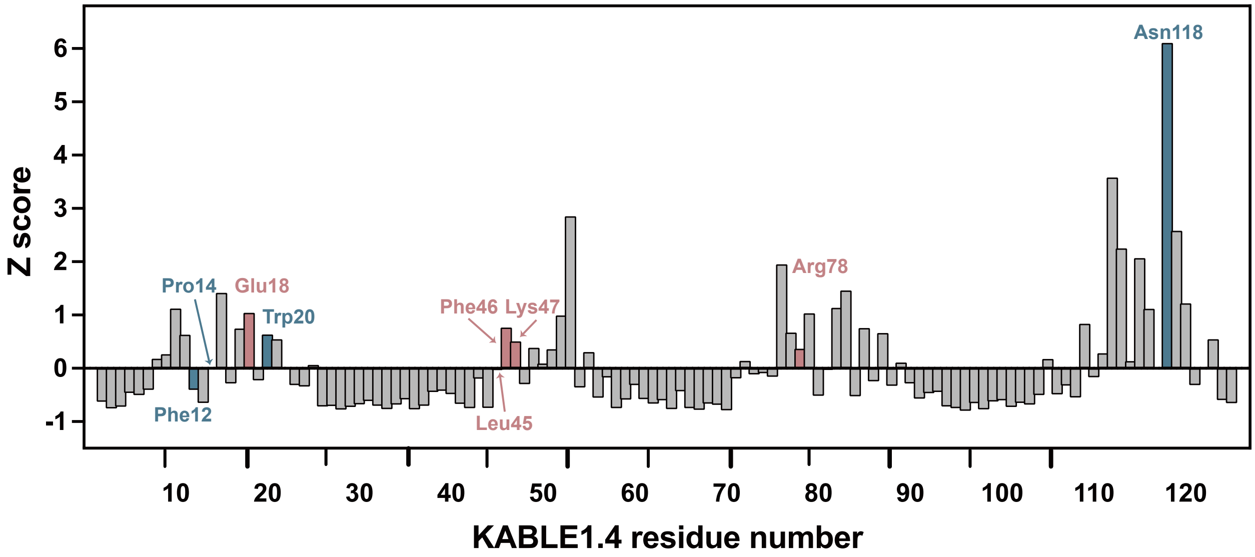


**Supplementary Figure 16 | Backbone amide chemical shift perturbations (CSPs) of KABLE1.4 upon addition of 5 molar equivalents of 6NBT.** The blue and red bars indicate the positions where beneficial mutations were identified through site saturation mutagenesis and NMR-guided directed evolution, respectively.

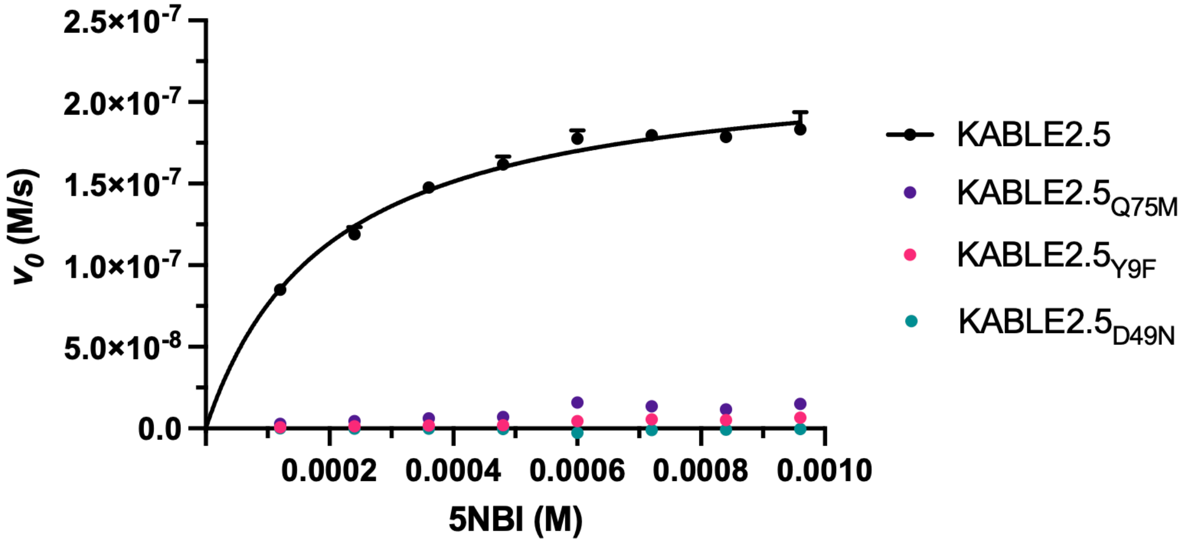


**Supplementary Figure 17 | Michaelis-Menten curve of KABLE2.5 and its negative mutants at 25 mM TRIS, pH 8.0, 100 mM NaCl, 1.5% MeCN.**

50 mM sodium phosphate, pH 7.0,
100 mM NaCl, 10% MeOH

| Protein | $k_{cat}$ (s-1) | $K_M$ (mM) | $k_{cat}/K_M$ ($M^{-1}s^{-1}$) |
|---|---|---|---|
| HG3.17 | 106 ± 17 | 0.87 ± 0.25 | 122,000 ± 40,000 |
| KABLE2.5 | 508 ± 23 | 0.61 ± 0.06 | 833,000 ± 86,000 |

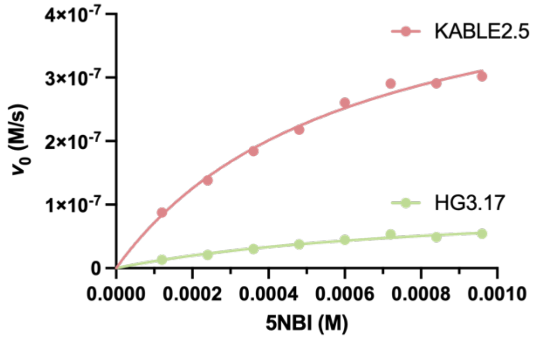


50 mM Bis-tris propane, pH 8.0,
100 mM NaCl, 10% MeOH

| Protein | $k_{cat}$ (s-1) | $K_M$ (mM) | $k_{cat}/K_M$ ($M^{-1}s^{-1}$) |
|---|---|---|---|
| HG3.17 | 30 ± 3 | 0.42 ± 0.09 | 72,000 ± 17,000 |
| KABLE2.5 | 291 ± 13 | 0.38 ± 0.04 | 768,000 ± 96,000 |

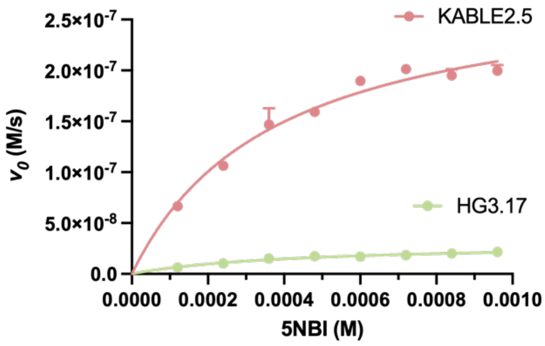


**Supplementary Figure 18 | Michaelis-Menten curve of HG3.17 and KABLE2.5 at indicated buffer conditions.**

**A**

```
ABLE              SVKSEYAEAAAVGQEAVAVFNTMKAAFQNGDKEAVAQYLARLASLYTRHEELLNRILEKA 60
KABLE1.2          SLKEKFEEYEKIGPRILELWQEARDAFEAGDLARVDELLRELKELFKKDLKLAEEMKKEA 60
KABLE1.2.crystal  SLKEKFAEYEAIGPRILELWQAARNAFEAGDLARVANLLAELKELFKKDLNLANAMAAEA 60

ABLE              RREGNKEAVTLMNEFTATFQTGKSIFNAMVAAFKNGDDDSFESYLQALEKVTAKGETLAD 120
KABLE1.2          EEAGNKEAVELLEEQLERLKKIQAMFEEAVEAFRAGDRERFGELLEKIIEEGKALLPLVE 120
KABLE1.2.crystal  AEAGNKEAVALLAEQLERLKKIQAMFAAAVNAFRAGDREAFGALLEAIINEGKALLPLVE 120

ABLE              QIAKAL 126
KABLE1.2          KIKEAI 126
KABLE1.2.crystal  AIKEAI 126
```

**B**

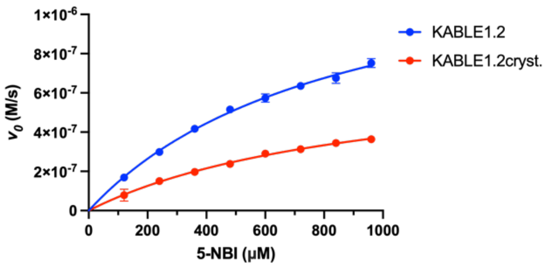


**Supplementary Figure 19 | Substrate concentration versus rate data for KABLE1.2 and KABLE1.2$_{cryst}$. (A)** Amino acid sequence alignment of ABLE with KABLE1.2 and KABLE1.2$_{cryst}$, shows the surface residues of ABLE were grafted to KABLE1.2$_{cryst}$. **(B)** Reactions were measured at pH 8.0 buffer (20 mM Tris,100 mM NaCl) with 0.1 μM KABLE1.2 and KABLE1.2$_{cryst,}$ respectively. Error bars represent standard deviations.

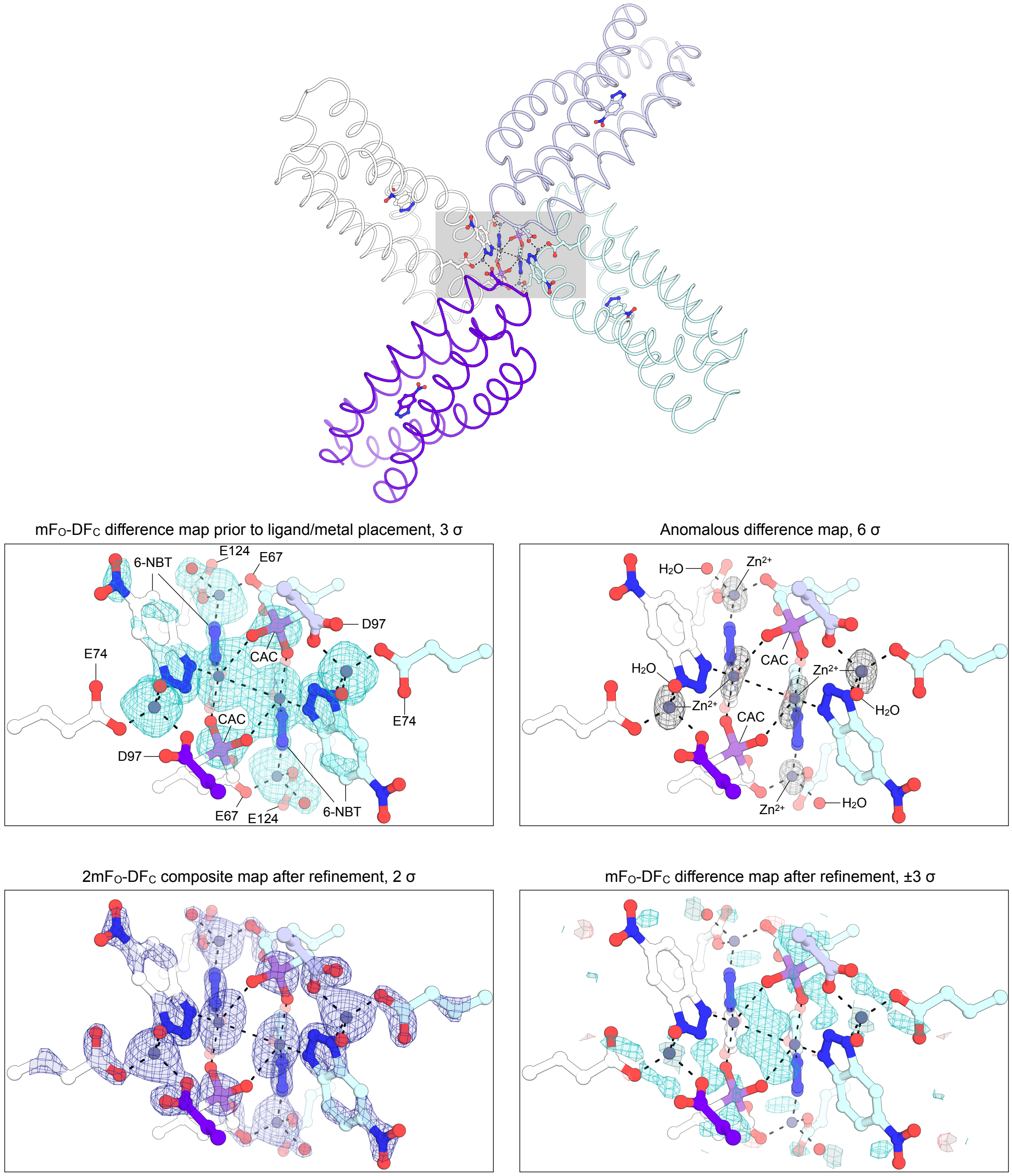


**Supplementary Figure 20 | Stabilization of the KABLE12$_{cryst}$ crystal lattice by a 6NBT/zinc/cacodylate complex.** The interface between four symmetry-related molecules is shown in the top panel. The gray shaded region is shown in the bottom panels with electron density maps used for assignment and modeling. The modeling of zinc (Zn) complexed by cacodylate (CAC) is supported by the anomalous difference map that shows strong peaks corresponding to the zinc positions and no peaks at the arsenic position, as expected for anomalous scattering for the two elements at 11111 eV. Although all features are supported by the mF$_O$-DF$_C$ difference map prior to complex modeling and the 2mF$_O$-DF$_C$ composite map after refinement, there are residue positive peaks in the mF$_O$-DF$_C$ difference map after refinement.

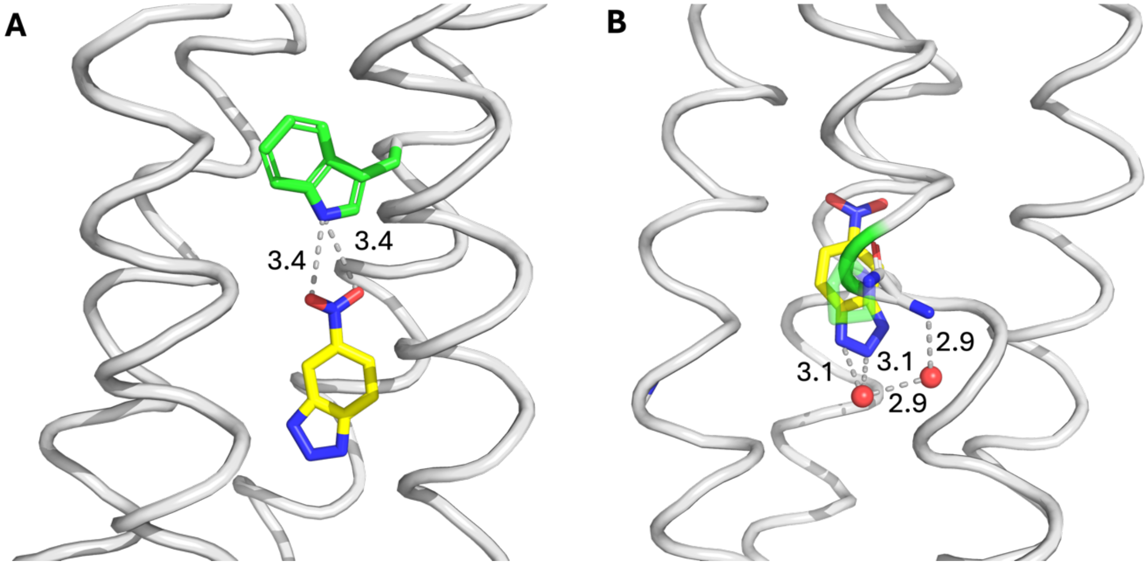


**Supplementary Figure 21 | Representative MD snapshots showing the hydrogen bond introduced by Leu20Trp and the active site water introduced by Lys14Pro**. **(A)** Representative MD snapshot showing the hydrogen bond between Trp20 and 6NBT. Trp20 is shown in green stick and 6NBT is shown in yellow stick. **(B)** Representative MD snapshot showing the water-mediated interaction to 6NBT, introduced by Pro14-mediated backbone kink. Distances are shown in Å with dashed lines.

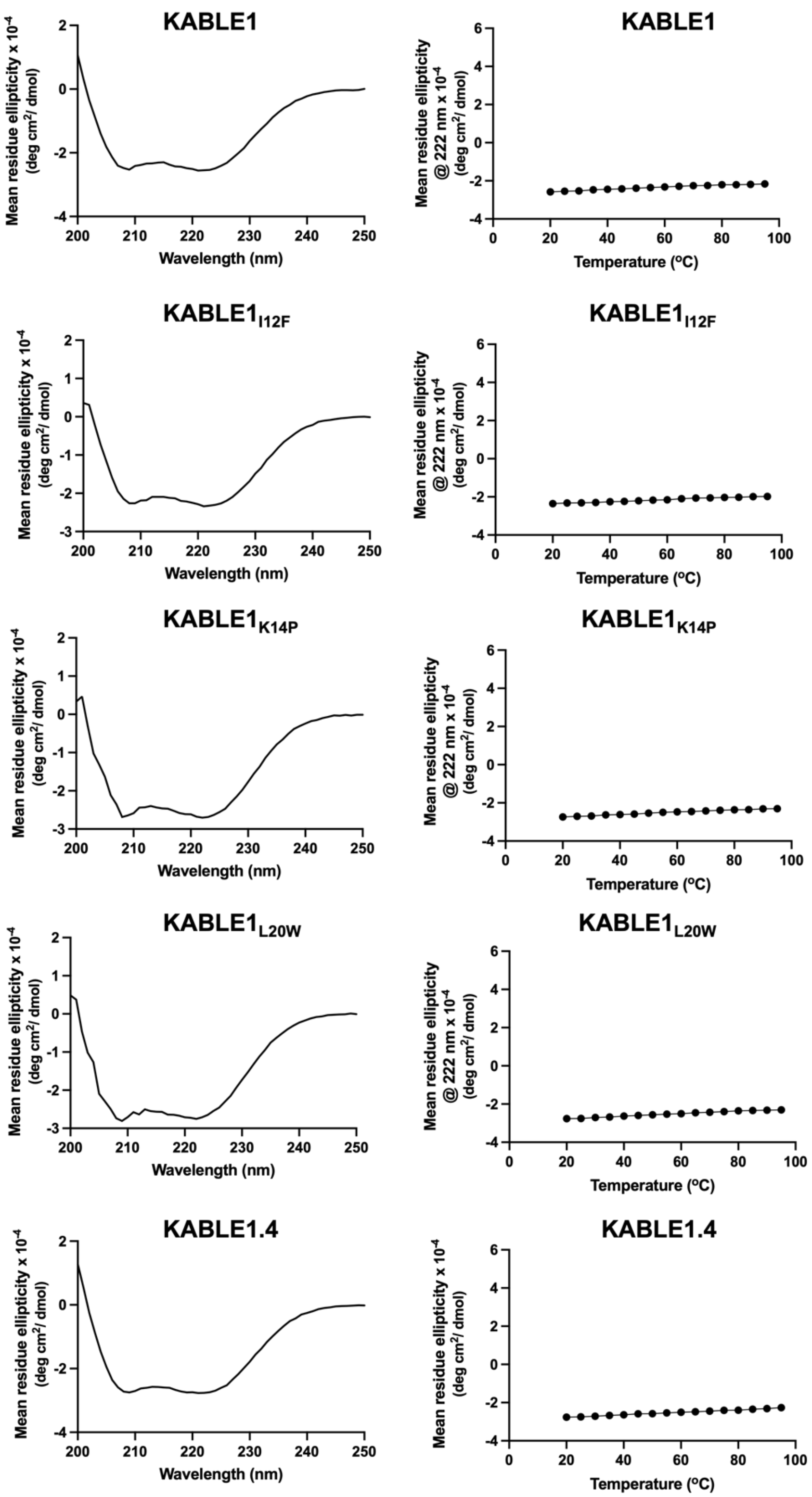


**Supplementary Figure 22 | Thermostability of KABLEs.** Proteins were prepared in 10 μM concentrations in PBS buffer. CD spectra were collected on a Jasco J-810 CD spectrometer in a 0.1 cm path-length quartz cuvette. Temperature-dependent data were collected at 222 nm from 20 to 95 °C with an interval of 5 °C and an increase rate of 2 °C/minute.

### Supplementary Tables

**Supplementary Table 1 | Ligand efficiency for starting ABLE-apixaban and FABLE-coumarin interactions.**

| | Protein | Ligand | ΔG (kcal/mol) | #Heavy Atoms | Ligand Efficiency |
|---|---|---|---|---|---|
| 1 | ABLE | Apixaban | -7.23 | 34 | 0.213 |
| 2 | $FABLE_{D1}$[a] | Cou485 | -6.21 | 18 | 0.345 |
| 3 | $FABLE_{D2}$ | Cou485 | -6.19 | 18 | 0.344 |
| 4 | $FABLE_{D3}$ | Cou485 | -6.02 | 18 | 0.334 |
| 5 | $FABLE_{D4}$ | Cou485 | -5.56 | 18 | 0.309 |
| 6 | $FABLE_{D5}$ | Cou485 | -5.70 | 18 | 0.317 |
| 7 | $FABLE_{D1}$[a] | Cou481 | -7.14 | 18 | 0.357 |
| 8 | $FABLE_{D1}$[a] | Cou540a | -7.77 | 18 | 0.353 |

[a]$FABLE_{D1}$ was designated as FABLE.

**Supplementary Table 2 | Cα RMSD (Å) of FABLE design with AF3 prediction and each monomer of its 3 crystal structures. (**PDB: 9DWA, 9DWB, 9DWC)

| | Design | 9DWA_A | 9DWA_B | 9DWB_A | 9DWB_B | 9DWC_A | 9DWC_B | 9DWC_C | 9DWC_D | AF3 |
|---|---|---|---|---|---|---|---|---|---|---|
| Design | | 0.78 | 0.79 | 0.93 | 0.91 | 0.91 | 0.99 | 1.01 | 0.93 | 0.61 |
| 9DWA_A | 0.78 | | 0.11 | 0.49 | 0.53 | 0.46 | 0.66 | 0.66 | 0.47 | 0.55 |
| 9DWA_B | 0.79 | 0.11 | | 0.51 | 0.54 | 0.47 | 0.67 | 0.67 | 0.48 | 0.57 |
| 9DWB_A | 0.93 | 0.49 | 0.51 | | 0.19 | 0.32 | 0.52 | 0.53 | 0.34 | 0.69 |
| 9DWB_B | 0.91 | 0.53 | 0.54 | 0.19 | | 0.32 | 0.49 | 0.50 | 0.36 | 0.70 |
| 9DWC_A | 0.91 | 0.46 | 0.47 | 0.32 | 0.32 | | 0.43 | 0.42 | 0.13 | 0.74 |
| 9DWC_B | 0.99 | 0.66 | 0.67 | 0.52 | 0.49 | 0.43 | | 0.14 | 0.44 | 0.88 |
| 9DWC_C | 1.01 | 0.66 | 0.67 | 0.53 | 0.50 | 0.42 | 0.14 | | 0.42 | 0.89 |
| 9DWC_D | 0.93 | 0.47 | 0.48 | 0.34 | 0.36 | 0.13 | 0.44 | 0.42 | | 0.78 |
| AF3 | 0.61 | 0.55 | 0.57 | 0.69 | 0.70 | 0.74 | 0.88 | 0.89 | 0.78 | |

**Supplementary Table 3 | Fluorophore library used to probe the specificity of FABLE.**

| Location | Name | Excitation | Emission | Structure |
|---|---|---|---|---|
| A1 | Coumarin440 | 350 | 440 | 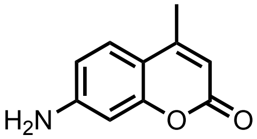 |
| A2 | Coumarin445 | 355 | 445 | 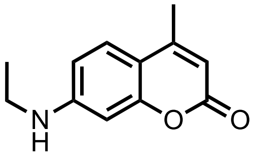 |
| A3 | Coumarin450 | 360 | 450 | 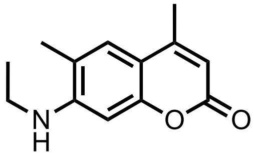 |
| A4 | Coumarin456 | 366 | 456 | 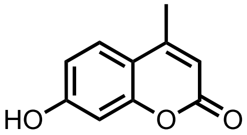 |
| A5 | Coumarin460 | 370 | 460 | 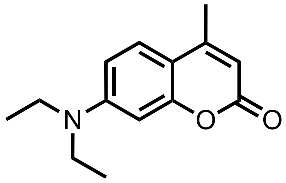 |
| A6 | Coumarin461 | 371 | 461 | 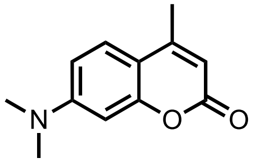 |
| A7 | Coumarin466 | 376 | 466 | 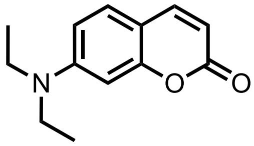 |
| A8 | LD473 | 383 | 473 | 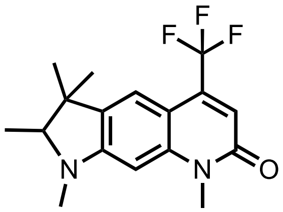 |

| | | | | |
|---|---|---|---|---|
| B1 | Coumarin478 | 388 | 478 | 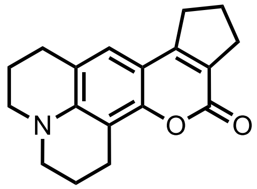 |
| B2 | Coumarin480 | 390 | 480 | 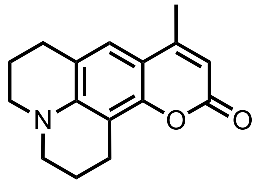 |
| B3 | Coumarin481 | 391 | 481 | 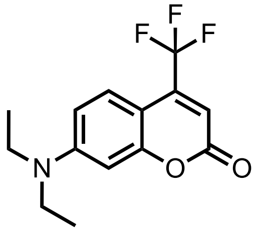 |
| B4 | Coumarin485 | 395 | 485 | 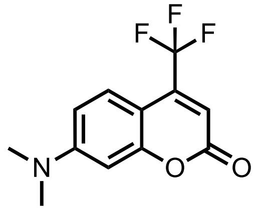 |
| B5 | LD489 | 399 | 489 | 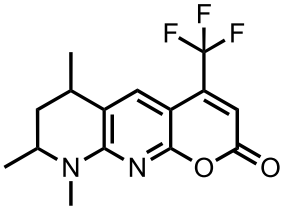 |
| B6 | Coumarin490 | 400 | 490 | 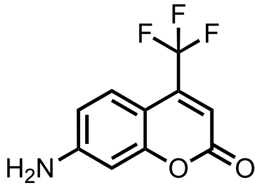 |
| B7 | LD490 | 400 | 490 | 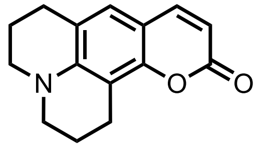 |
| B8 | Coumarin498 | 408 | 498 | 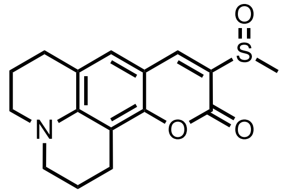 |

| | | | | |
|---|---|---|---|---|
| C1 | Coumarin500 | 410 | 500 | 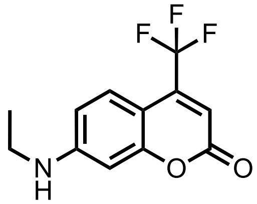 |
| C2 | Coumarin503 | 413 | 503 | 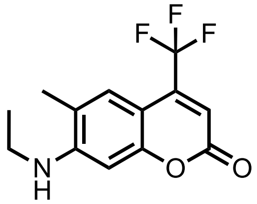 |
| C3 | Coumarin504 | 414 | 504 | 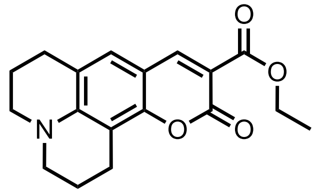 |
| C4 | Coumarin510 | 420 | 510 | 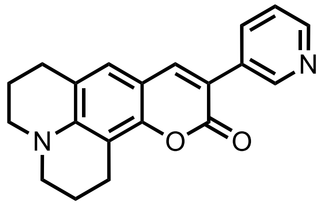 |
| C5 | Coumarin515 | 425 | 515 | 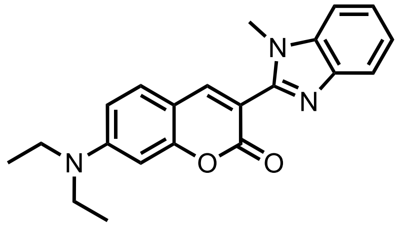 |
| C6 | Coumarin519 | 429 | 519 | 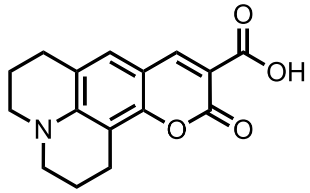 |
| C7 | Coumarin521 | 431 | 521 | 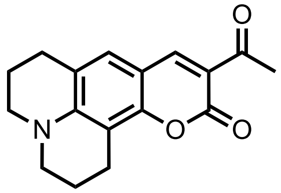 |
| C8 | Coumarin523 | 433 | 523 | 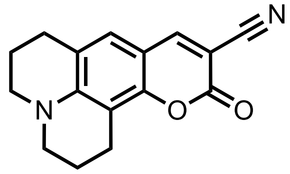 |

| | | | | |
|---|---|---|---|---|
| D1 | Coumarin525 | 435 | 525 | 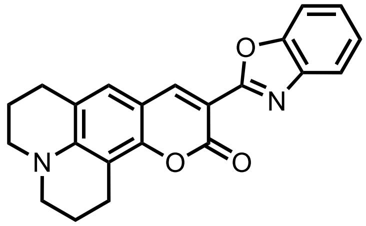 |
| D2 | Coumarin535 | 445 | 535 | 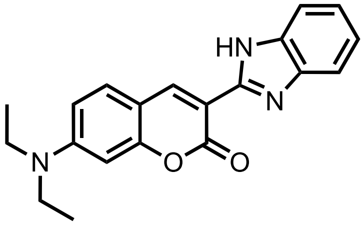 |
| D3 | Coumarin545 | 455 | 545 | 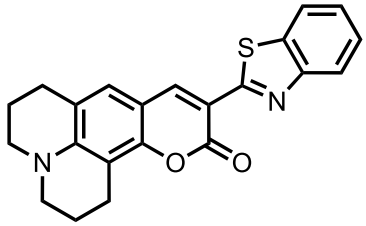 |
| D4 | Coumarin540a | 460 | 540a | 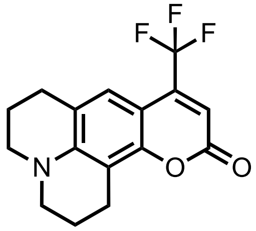 |
| L036 | Pyrromethene556 | 492 | 518 | 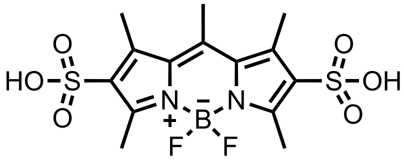 |
| L045 | Fluorol555 | 460 | 559 | 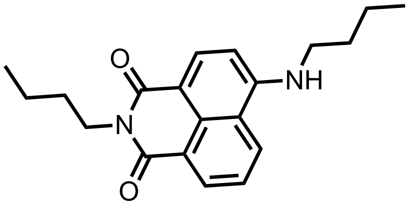 |
| L068 | Phenoxazone680 | 586 | 670 | 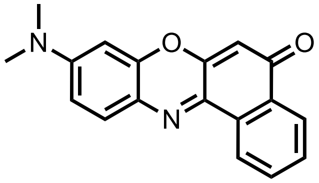 |
| L069 | CresylViolet670Perchlorate | 592 | 628 | 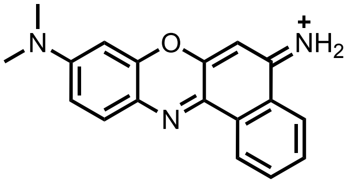 |

| | | | | |
|---|---|---|---|---|
| L070 | NileBlue690Perchlorate | 636 | 682 | 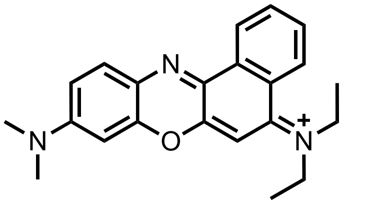 |
| L071 | LD690Perchlorate | 616 | 638 | 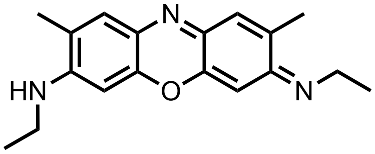 |
| L093 | HICl | 539 | 569 | 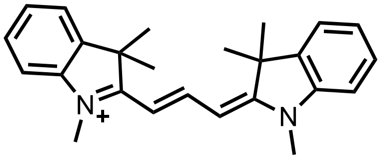 |
| TMR | TMR | 550 | 575 | 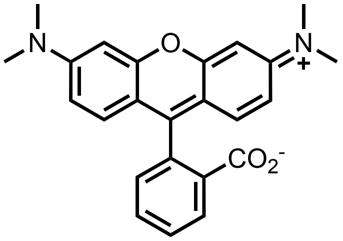 |

**Supplementary Table 4 | Protein concentrations (nM) used in kinetic assay of Kemp eliminase.**

| pH | KABLE1 | KABLE1$_{I12F}$ | KABLE1$_{K14P}$ | KABLE1$_{L20W}$ | KABLE1$_{L118N}$ | KABLE 1.2 | KABLE 1.3 | KABLE 1.4 | KABLE 2.5 |
|---|---|---|---|---|---|---|---|---|---|
| 6 | | 1000 | | 1000 | 1000 | 100 | 100 | 5 | |
| 6.5 | | 1000 | 100 | 1000 | 1000 | 100 | 100 | 5 | 1 |
| 7 | 1000 | 1000 | 100 | 1000 | 1000 | 10 | 10 | 5 | 1 |
| 7.5 | 1000 | 1000 | 100 | 1000 | 1000 | 10 | 10 | 5 | 1 |
| 8 | 1000 | 1000 | 100 | 1000 | 1000 | 10 | 10 | 5 | 1 |
| 8.5 | 1000 | 1000 | 100 | 1000 | 1000 | 10 | 10 | 5 | 1 |
| 9 | 100 | 100 | 10 | 100 | 100 | 10 | 10 | 5 | 1 |
| 9.5 | 100 | 100 | | 100 | 100 | 10 | 10 | 5 | 1 |
| 10 | 100 | 100 | | 100 | 100 | 10 | 10 | 5 | 1 |

**Supplementary Table 5 | Kinetic parameters of KABLEs at pH 8.**

| Protein | $k_{cat}$, $s^{-1}$ | $K_M$, mM | ($k_{cat}/K_M$), $M^{-1}s^{-1}$ |
|---|---|---|---|
| KABLE1 | 0.09 ± 0.001 | 0.20 ± 0.01 | 460 ± 20 |
| KABLE1$_{I12F}$ | 0.28 ± 0.01 | 0.37 ± 0.03 | 740 ± 60 |
| KABLE1$_{K14P}$ | 16 ± 4 | 4.3 ± 1 | 3,800 ± 1,400 |
| KABLE1$_{L20W}$ | 0.33 ± 0.01 | 0.24 ± 0.03 | 1,300 ± 200 |
| KABLE1$_{L118N}$ | 0.20 ± 0.01 | 0.33 ± 0.02 | 600 ± 40 |
| KABLE1.2 | 12 ± 1 | 0.64 ± 0.1 | 19,000 ± 5,000 |
| KABLE1.3 | 62 ± 3 | 0.9 ± 0.09 | 68,000 ± 8,000 |
| KABLE1.4 | 99 ± 5 | 0.43 ± 0.05 | 230,000 ± 32,000 |
| KABLE1.4$_{E18G}$ | 174 ± 6 | 0.53 ± 0.04 | 330,000 ± 28,000 |
| KABLE1.4$_{F46L}$ | 280 ± 10 | 0.55 ± 0.07 | 450,000 ± 60,000 |
| KABLE1.4$_{K47R}$ | 175 ± 6 | 0.49 ± 0.04 | 350,000 ± 30,000 |
| KABLE1.4$_{L45I}$ | 150 ± 8 | 0.55 ± 0.06 | 270,000 ± 30,000 |
| KABLE1.4$_{R78G}$ | 154 ± 4 | 0.27 ± 0.02 | 570,000 ± 40,000 |

[a]Error bars represent the standard errors of the mean from three independent measurements.
Assay condition: 25 mM TRIS, pH 8.0, 100 mM NaCl, 1.5% MeCN, RT.

**Supplementary Table 6 | Ca RMSD of designed KABLE1, Chai1 prediction of 6NBT-bound structures, and each monomer of its 2 crystal structures.** (PDB: 9N0I,9N0J )

| | 9N0I_A | 9N0I_B | 9N0I_C | 9N0I_D | 9N0I_E | 9N0I_F | 9N0J_A | KABLE1_Chai1 | KABLE12cryst_Chai1 | KABLE1_Design |
|---|---|---|---|---|---|---|---|---|---|---|
| 9N0I_A | | 0.5 | 0.4 | 0.6 | 0.5 | 0.6 | 0.6 | 0.7 | 0.6 | 0.8 |
| 9N0I_B | 0.5 | | 0.3 | 0.5 | 0.5 | 0.6 | 0.4 | 0.7 | 0.6 | 0.9 |
| 9N0I_C | 0.4 | 0.3 | | 0.4 | 0.5 | 0.5 | 0.4 | 0.7 | 0.5 | 0.8 |
| 9N0I_D | 0.6 | 0.5 | 0.4 | | 0.3 | 0.3 | 0.6 | 0.7 | 0.4 | 1.1 |
| 9N0I_E | 0.5 | 0.5 | 0.5 | 0.3 | | 0.4 | 0.7 | 0.7 | 0.5 | 1.0 |
| 9N0I_F | 0.6 | 0.6 | 0.5 | 0.3 | 0.4 | | 0.7 | 0.7 | 0.4 | 1.1 |
| 9N0J_A | 0.6 | 0.4 | 0.4 | 0.6 | 0.7 | 0.7 | | 0.8 | 0.7 | 0.9 |
| KABLE1_Chai1 | 0.7 | 0.7 | 0.7 | 0.7 | 0.7 | 0.7 | 0.8 | | 0.5 | 0.7 |
| KABLE12cryst_Chai1 | 0.6 | 0.6 | 0.5 | 0.4 | 0.5 | 0.4 | 0.7 | 0.5 | | 0.9 |
| KABLE1_Design | 0.8 | 0.9 | 0.8 | 1.1 | 1.0 | 1.1 | 0.9 | 0.7 | 0.9 | |

**Supplementary Table 7 | Ca RMSD of GPCRs with TM alignment (Fig.6e).**

| | 2RH1 | 6ME9 | 6HLO | 6OIK | 6VMS | 7WC4 | 8EF5 |
|---|---|---|---|---|---|---|---|
| 2RH1 | 0 | 3.03 | 2.8 | 2.65 | 2.49 | 2.79 | 3.41 |
| 6ME9 | 3.03 | 0 | 4.07 | 3.18 | 3.21 | 4.03 | 3.58 |
| 6HLO | 2.8 | 4.07 | 0 | 3.08 | 2.96 | 3.98 | 3.33 |
| 6OIK | 2.65 | 3.18 | 3.08 | 0 | 2.3 | 2.82 | 2.79 |
| 6VMS | 2.49 | 3.21 | 2.96 | 2.3 | 0 | 2.67 | 2.89 |
| 7WC4 | 2.79 | 4.03 | 3.98 | 2.82 | 2.67 | 0 | 3.33 |
| 8EF5 | 3.41 | 3.58 | 3.33 | 2.79 | 2.89 | 3.33 | 0 |

Chain A of each PDB was used to calculate the Ca RMSD of the aligned residues using TM alignment[8], except for 6VMS, whose chain E was used.

**Supplementary Table 8 | Ca RMSD of four-helical bundles with TM alignment (Fig. 6d).**

| | 6W70 | 9DWC | 9N0J | 9DM3[a] | 8TN6 | 7JRQ | 5TGY | 7JH6 | 7AH0 | 8D9P |
|---|---|---|---|---|---|---|---|---|---|---|
| 6W70 | 0 | 0.84 | 1.13 | 1.42 | 1.28 | 1.43 | 1.75 | 1.65 | 1.71 | 1.77 |
| 9DWC | 0.84 | 0 | 1.18 | 1.55 | 1.39 | 1.63 | 1.79 | 1.77 | 1.72 | 1.73 |
| 9N0J | 1.13 | 1.18 | 0 | 1.58 | 1.14 | 1.6 | 1.91 | 1.75 | 1.81 | 1.85 |
| 9DM3[a] | 1.42 | 1.55 | 1.58 | 0 | 1.62 | 2.23 | 2.08 | 2.32 | 2.49 | 3.04 |
| 8TN6 | 1.28 | 1.39 | 1.14 | 1.62 | 0 | 1.82 | 1.28 | 2.18 | 1.74 | 1.83 |
| 7JRQ | 1.43 | 1.63 | 1.6 | 2.23 | 1.82 | 0 | 1.69 | 2.27 | 1.83 | 2.22 |
| 5TGY | 1.75 | 1.79 | 1.91 | 2.08 | 1.28 | 1.69 | 0 | 1.87 | 2.11 | 2.36 |
| 7JH6 | 1.65 | 1.77 | 1.75 | 2.32 | 2.18 | 2.27 | 1.87 | 0 | 2 | 2.46 |
| 7AH0 | 1.71 | 1.72 | 1.81 | 2.49 | 1.74 | 1.83 | 2.11 | 2 | 0 | 1.85 |
| 8D9P | 1.77 | 1.73 | 1.85 | 3.04 | 1.83 | 2.22 | 2.36 | 2.46 | 1.85 | 0 |

Chain A of each PDB was used to calculate the Ca RMSD of the aligned residues using TM alignment[8].

[a]The first state of NMR structure was used.

**Supplementary Table 9 | Primer sequences used in this study.**

| Primer | Sequence |
|---|---|
| T7 | TAATACGACTCACTATAGGG |
| T7_term | GCTAGTTATTGCTCAGCGG |
| **Active Site Mutagenesis** | |
| Y9NNK | AAGTTTGAGGAG**NNK**GAGAAAATCGGG |
| I12NNK | GAGTATGAGAAA**NNK**GGGAAGCGTATT |
| K14NNK | GAGAAAATCGGG**NNK**CGTATTCTCGAA |
| I16NNK | ATCGGGAAGCGT**NNK**CTTGAACTGTTA |
| L17NNK | GGGAAGCGTATT**NNK**GAACTGTTACAG |
| L20NNK | ATTCTCGAACTG**NNK**CAGGAAGCCCGT |
| L42NNK | TTACTTCGGGAA**NNK**AAGGAGCTTTTT |
| F46NNK | TTAAAGGAGCTT**NNK**AAGAAGGATCTC |
| L50NNK | TTTAAGAAGGAT**NNK**AAGTTAGCTGAA |
| L52NNK | TTTAAGAAGGATCTCAAG**NNK**GCTGAA |
| Q75NNK | CTGCTGGAAGAA**NNK**CTCGAACGTCTC |
| L79NNK | CAGCTCGAACGT**NNK**AAGAAAATCCAG |
| I82NNK | CGTCTCAAGAAA**NNK**CAGGCGATGTTC |
| G112NNK | ATCATCGAGGAG**NNK**AAAGCACTTTTA |
| K113NNK | ATCGAGGAGGGG**NNK**GCACTTTTACCT |
| L115NNK | GAGGGGAAAGCA**NNK**TTACCTCTTGTG |
| L116NNK | GGGAAAGCACTT**NNK**CCTCTTGTGGAG |
| P117NNK | GGGAAAGCACTTTTA**NNK**CTTGTGGAG |
| L118NNK | GCACTTTTACCT**NNK**GTGGAGAAGATC |
| V119NNK | CTTTTACCTCTT**NNK**GAGAAGATCAAA |
| **NMR-Guided Directed Evolution** | |
| R15NNK | GAGAAATTTGGGCCG**NNK**ATTCTTGAA |
| I16NNK | AAATTTGGGCCGCGT**NNK**CTTGAACTG |
| L17NNK | TTTGGGCCGCGTATT**NNK**GAACTGTGG |
| E18NNK | GGGCCGCGTATTCTT**NNK**CTGTGGCAG |
| L45NNK | CGGGAATTAAAGGAG**NNK**TTTAAGAAG |
| F46NNK | GAATTAAAGGAGCTT**NNK**AAGAAGGAT |
| K47NNK | AAGGAGCTTTTT**NNK**AAGGATCTCAAG |
| K48NNK | AAGGAGCTTTTTAAG**NNK**GATCTCAAG |
| L52NNK | AAGGATCTCAAG**NNK**GCTGAAGAAATG |
| A53NNK | AAGGATCTCAAGTTA**NNK**GAAGAAATG |
| E54NNK | AAGGATCTCAAGTTAGCT**NNK**GAAATG |
| Q75NNK | GAGCTGCTGGAAGAA**NNK**CTCGAACGT |
| L76NNK | CTGCTGGAAGAACAG**NNK**GAACGTCTC |
| E77NNK | CTGGAAGAACAGCTC**NNK**CGTCTCAAG |
| R78NNK | GAAGAACAGCTCGAA**NNK**CTCAAGAAA |
| L79NNK | GAACAGCTCGAACGT**NNK**AAGAAAATC |
| K80NNK | CAGCTCGAACGTCTC**NNK**AAAATCCAG |
| I82NNK | GAACGTCTCAAGAAA**NNK**CAGGCGATG |
| Q83NNK | CGTCTCAAGAAAATC**NNK**GCGATGTTC |

| | |
|---|---|
| A84NNK | CTCAAGAAAATCCAG**NNK**ATGTTCGAG |
| G112NNK | AAGATCATCGAGGAG**NNK**AAAGCACTT |
| K113NNK | ATCATCGAGGAGGGG**NNK**GCACTTTTA |
| A114NNK | ATCGAGGAGGGGAAA**NNK**CTTTTACCT |
| L115NNK | GAGGAGGGGAAAGCA**NNK**TTACCTAAC |

**Supplementary Table 10 | Amino acid sequences of designed proteins and their mutants.**

| Protein | Amino acid sequence | DNA Sequence |
|---|---|---|
| FABLE | MHHHHHHENLYFQSIKSEFAEGAAVFVEG<br>VAVFLTMMAAFQNGDKEAVAQYLARGASL<br>YTRHEELLNRLLQKLRREGNKEAVTLMNE<br>FTATFQTGKSLANALIAAFKNGDDDSFES<br>YLQAIWKVIAKMATILDQIAKAI | ATGCACCACCACCACCACCACGA<br>AAACCTGTACTTCCAGTCAATCA<br>AGAGCGAGTTCGCCGAGGGGGCG<br>GCTGTATTTGTAGAAGGCGTCGC<br>CGTGTTTCTGACCATGATGGCCG<br>CATTTCAGAACGGCGATAAGGAG<br>GCTGTCGCACAGTACCTCGCGCG<br>GGGGGCTTCCCTGTACACCCGTC<br>ACGAGGAGTTGCTGAACCGGCTT<br>TTACAAAAGTTGCGGCGCGAAGG<br>GAACAAAGAAGCAGTGACTCTTA<br>TGAACGAATTTACCGCCACTTTT<br>CAAACTGGTAAGTCCTTAGCTAA<br>TGCTCTGATCGCGGCATTCAAAA<br>ATGGTGACGATGACTCCTTTGAA<br>AGTTACCTTCAAGCAATCTGGAA<br>GGTCATCGCCAAGATGGCAACTA<br>TTTTGGATCAAATTGCAAAAGCA<br>ATT |
| KABLE0 | MHHHHHHENLYFQSLKELFEEWLEVRKKID<br>EVFDAMKAAFEAGDMKTVGEYIKELKELYK<br>EYLAMLEELLRRARALGLEEAVALLEEYKG<br>LVEEASKAFEAAVKAFEAGDWAAFKKYLEK<br>EEEIRKKMEPLVKALEEAL | ATGCACCACCACCACCACCACGA<br>AAACCTGTACTTCCAGAGCTTAA<br>AAGAGTTATTTGAAGAGTGGTTG<br>GAAGTTCGTAAGAAGATTGATGA<br>AGTATTCGATGCAATGAAGGCCG<br>CTTTCGAGGCCGGGGATATGAAA<br>ACAGTGGGGGAGTATATCAAAGA<br>ATTGAAGGAACTTTATAAAGAAT<br>ACTTAGCTATGCTTGAGGAGTTG<br>CTCCGGCGTGCTCGTGCCTTAGG<br>TCTTGAAGAGGCAGTCGCATTAC<br>TGGAAGAGTACAAAGGGTTAGTC<br>GAGGAGGCATCGAAGGCTTTTGA<br>AGCTGCCGTAAAAGCTTTCGAGG<br>CTGGGGACTGGGCCGCCTTCAAA<br>AAATACCTTGAGAAGGAAGAGGA<br>GATCCGCAAAAAGATGGAACCGC<br>TCGTCAAAGCCCTGGAAGAGGCA<br>CTC |
| KABLE1 | MHHHHHHENLYFQSSLKEKFEEYEKIGKRI<br>LELLQEARDAFEAGDLARVDELLRELKELF<br>KKDLKLAEEMKKEAEEAGNKEAVELLEEQL<br>ERLKKIQAMFEEAVEAFRAGDRERFGELLE<br>KIIEEGKALLPLVEKIKEAI | ATGCACCACCACCACCACCACGA<br>AAACCTGTACTTCCAGAGCAGCC<br>TCAAAGAAAAGTTTGAGGAGTAT<br>GAGAAAATCGGGAAGCGTATTCT<br>CGAACTGTTACAGGAAGCCCGTG<br>ACGCTTTCGAGGCTGGTGATCTC<br>GCGCGGGTTGATGAGTTACTTCG<br>GGAATTAAAGGAGCTTTTTAAGA |

| | | |
|---|---|---|
| | | AGGATCTCAAGTTAGCTGAAGAA<br>ATGAAAAAGGAAGCGGAAGAAGC<br>TGGGAATAAAGAGGCGGTGGAGC<br>TGCTGGAAGAACAGCTCGAACGT<br>CTCAAGAAAATCCAGGCGATGTT<br>CGAGGAGGCAGTTGAGGCGTTTC<br>GGGCCGGGGACCGCGAGCGTTTT<br>GGTGAGCTCCTCGAAAAGATCAT<br>CGAGGAGGGGAAAGCACTTTTAC<br>CTCTTGTGGAGAAGATCAAAGAG<br>GCTATC |
| KABLE1.2 | MHHHHHHENLYFQSSLKEKFEEYEKIGPRI<br>LELWQEARDAFEAGDLARVDELLRELKELF<br>KKDLKLAEEMKKEAEEAGNKEAVELLEEQL<br>ERLKKIQAMFEEAVEAFRAGDRERFGELLE<br>KIIEEGKALLPLVEKIKEAI | ATGCACCACCACCACCACCACGA<br>AAACCTGTACTTCCAGAGCAGCC<br>TCAAAGAAAAGTTTGAGGAGTAT<br>GAGAAAATCGGGCCGCGTATTCT<br>TGAACTGTGGCAGGAAGCCCGTG<br>ACGCTTTCGAGGCTGGTGATCTC<br>GCGCGGGTTGATGAGTTACTTCG<br>GGAATTAAAGGAGCTTTTTAAGA<br>AGGATCTCAAGTTAGCTGAAGAA<br>ATGAAAAAGGAAGCGGAAGAAGC<br>TGGGAATAAAGAGGCGGTGGAGC<br>TGCTGGAAGAACAGCTCGAACGT<br>CTCAAGAAAATCCAGGCGATGTT<br>CGAGGAGGCAGTTGAGGCGTTTC<br>GGGCCGGGGACCGCGAGCGTTTT<br>GGTGAGCTCCTCGAAAAGATCAT<br>CGAGGAGGGGAAAGCACTTTTAC<br>CTCTTGTGGAGAAGATCAAAGAG<br>GCTATC |
| KABLE1.2$_{cryst}$ | MHHHHHHENLYFQSSLKEKFAEYEAIGPRI<br>LELWQAARNAFEAGDLARVANLLAELKELF<br>KKDLNLANAMAAEAAEAGNKEAVALLAEQL<br>ERLKKIQAMFAAAVNAFRAGDREAFGALLE<br>AIINEGKALLPLVEAIKEAI | ATGCACCACCACCACCACCACGA<br>AAACCTGTACTTCCAGAGCTCAC<br>TGAAGGAGAAATTCGCCGAATAC<br>GAGGCGATTGGGCCGCGGATTCT<br>CGAATTGTGGCAGGCAGCGCGTA<br>ATGCTTTCGAAGCTGGGGATCTC<br>GCTCGGGTTGCCAATTTGCTTGC<br>GGAATTGAAGGAACTGTTCAAGA<br>AAGACCTGAATCTTGCGAACGCA<br>ATGGCGGCCGAGGCGGCGGAAGC<br>CGGGAATAAGGAAGCTGTAGCCC<br>TTTTGGCGGAACAGCTTGAGCGG<br>TTAAAGAAAATCCAGGCAATGTT<br>CGCCGCGGCGGTGAATGCTTTCC<br>GGGCGGGGGACCGCGAGGCATTT<br>GGTGCATTGTTGGAGGCAATCAT<br>CAACGAAGGCAAGGCTCTCTTAC<br>CGCTCGTGGAGGCGATTAAGGAG<br>GCCATC |

| | | |
|---|---|---|
| KABLE1.3 | MHHHHHHENLYFQSSLKEKFEEYEKFGPRILELWQEARDAFEAGDLARVDELLRELKELFKKDLKLAEEMKKEAEEAGNKEAVELLEEQLERLKKIQAMFEEAVEAFRAGDRERFGELLEKIIEEGKALLPLVEKIKEAI | ATGCACCACCACCACCACCACGAAAACCTGTACTTCCAGAGCAGCCTCAAAGAAAAGTTTGAGGAGTATGAGAAATTTGGGCCGCGTATTCTTGAACTGTGGCAGGAAGCCCGTGACGCTTTCGAGGCTGGTGATCTCGCGCGGGTTGATGAGTTACTTCGGGAATTAAAGGAGCTTTTTAAGAAGGATCTCAAGTTAGCTGAAGAAATGAAAAAGGAAGCGGAAGAAGCTGGGAATAAAGAGGCGGTGGAGCTGCTGGAAGAACAGCTCGAACGTCTCAAGAAAATCCAGGCGATGTTCGAGGAGGCAGTTGAGGCGTTTCGGGCCGGGGACCGCGAGCGTTTTGGTGAGCTCCTCGAAAAGATCATCGAGGAGGGGAAAGCACTTTTACCTCTTGTGGAGAAGATCAAAGAGGCTATC |
| KABLE1.4 | MHHHHHHENLYFQSSLKEKFEEYEKFGPRILELWQEARDAFEAGDLARVDELLRELKELFKKDLKLAEEMKKEAEEAGNKEAVELLEEQLERLKKIQAMFEEAVEAFRAGDRERFGELLEKIIEEGKALLPNVEKIKEAI | ATGCACCACCACCACCACCACGAAAACCTGTACTTCCAGAGCAGCCTCAAAGAAAAGTTTGAGGAGTATGAGAAATTTGGGCCGCGTATTCTTGAACTGTGGCAGGAAGCCCGTGACGCTTTCGAGGCTGGTGATCTCGCGCGGGTTGATGAGTTACTTCGGGAATTAAAGGAGCTTTTTAAGAAGGATCTCAAGTTAGCTGAAGAAATGAAAAAGGAAGCGGAAGAAGCTGGGAATAAAGAGGCGGTGGAGCTGCTGGAAGAACAGCTCGAACGTCTCAAGAAAATCCAGGCGATGTTCGAGGAGGCAGTTGAGGCGTTTCGGGCCGGGGACCGCGAGCGTTTTGGTGAGCTCCTCGAAAAGATCATCGAGGAGGGGAAAGCACTTTTACCTAACGTGGAGAAGATCAAAGAGGCTATC |
| KABLE2.5 | MHHHHHHENLYFQSSLKEKFEEYEKFGPRILGLWQEARDAFEAGDLARVDELLRELKEILRKDLKLAEEMKKEAEEAGNKEAVELLEEQLEGLKKIQAMFEEAVEAFRAGDRERFGELLEKIIEEGKALLPNVEKIKEAI | ATGCACCACCACCACCACCACGAAAACCTGTACTTCCAGAGCAGCCTCAAAGAAAAGTTTGAGGAGTATGAGAAATTTGGGCCGCGTATTCTTGGTCTGTGGCAGGAAGCCCGTGACGCTTTCGAGGCTGGTGATCTCGCGCGGGTTGATGAGTTACTTCGGGAATTAAAGGAGATTTTGCGTAAGGATCTCAAGTTAGCTGAAGAAATGAAAAAGGAAGCGGAAGAAGCTGGGAATAAAGAGGCGGTGGAGCTGCTGGAAGAACAGCTCGAAGGTCTCAAGAAAATCCAGGCGATGTTCGAGGAGGCAGTTGAGGCGTTTCGGGCCGGGGACCGCGAGCGTTTTGGTGAGCTCCTCGAAAAGATCAT |

| | | |
|---|---|---|
| | | CGAGGAGGGGAAAGCACTTTTAC CTAACGTGGAGAAGATCAAAGAG GCTATCTAA |

### Supplementary Note

#### Computational design of FABLE_RosettaScript(.xml)

```
<ROSETTASCRIPTS>
  <SCOREFXNS>
    <ScoreFunction name="ligand_soft_rep" weights="ligand_soft_rep" />
    <ScoreFunction name="hard_rep" weights="ligandprime" />
    <ScoreFunction name="ref15" weights="ref2015">
      <Reweight scoretype="atom_pair_constraint" weight="5"/>
    </ScoreFunction>
    <ScoreFunction name="ref15_1" weights="ref2015">
      <Reweight scoretype="aa_composition" weight="1" />
      <Reweight scoretype="netcharge" weight="1.0" />
      <Reweight scoretype="atom_pair_constraint" weight="5"/>
      <Set aa_composition_setup_file="helix_composition.comp" />
      <Set netcharge_setup_file="helix_netcharge.charge" />
    </ScoreFunction>
  </SCOREFXNS>
  <RESIDUE_SELECTORS>
  </RESIDUE_SELECTORS>
  <TASKOPERATIONS>
    <InitializeFromCommandline name="ifcl"/>
    <ReadResfile name="resfile" filename="resfile_topology.txt"/>
    <ExtraRotamersGeneric name="extrachi" ex1="1" ex2="1" ex1_sample_level="1" ex2_sample_level="1"
extrachi_cutoff="14"/>
    <DetectProteinLigandInterface name="design_interface" cut1="6.0" cut2="8.0" cut3="10.0" cut4="12.0"
design="1" resfile="resfile_topology.txt" />
    IncludeCurrent name="include_curr" />
  </TASKOPERATIONS>
  <FILTERS>
    <PackStat name="pstat" confidence="0" threshold="0" repeats="10"/>
    <PackStat name="pstat_mc" threshold="0" repeats="10"/>
    <NetCharge name="net_charge" confidence="0"/>
    <LigInterfaceEnergy name="ligen"  scorefxn="ref15" include_cstE="0" energy_cutoff="0" confidence="0"/>
    <ScoreType name="total_score_1" scorefxn="ref15_1" score_type="total_score" threshold="0"/>
    <ScoreType name="total_score" scorefxn="ref15" score_type="total_score" threshold="0"/>
    <BuriedUnsatHbonds name="bu" report_sc_heavy_atom_unsats="true" scorefxn="ref15" cutoff="0"
residue_surface_cutoff="20.0" ignore_surface_res="true" print_out_info_to_pdb="true" confidence="0" />
  </FILTERS>
  <LIGAND_AREAS>
    <LigandArea name="docking_sidechain" chain="X" cutoff="6.0" add_nbr_radius="true"
all_atom_mode="true" minimize_ligand="10" />
    <LigandArea name="final_sidechain" chain="X" cutoff="6.0" add_nbr_radius="true" all_atom_mode="true"
/>
    <LigandArea name="final_backbone" chain="X" cutoff="7.0" add_nbr_radius="false" all_atom_mode="true"
Calpha_restraints="0.3" />
  </LIGAND_AREAS>
  <INTERFACE_BUILDERS>
    <InterfaceBuilder name="side_chain_for_docking" ligand_areas="docking_sidechain" />
    <InterfaceBuilder name="side_chain_for_final" ligand_areas="final_sidechain" />
    <InterfaceBuilder name="backbone" ligand_areas="final_backbone" extension_window="3" />
  </INTERFACE_BUILDERS>
  <MOVEMAP_BUILDERS>
```

```
        <MoveMapBuilder name="docking" sc_interface="side_chain_for_docking" minimize_water="true" />
        <MoveMapBuilder name="final" sc_interface="side_chain_for_final" bb_interface="backbone"
minimize_water="true" />
    </MOVEMAP_BUILDERS>
    <SCORINGGRIDS ligand_chain="X" width="46">
        <ClassicGrid grid_name="vdw" weight="1.0" />
    </SCORINGGRIDS>
    <MOVERS>
        <ConstraintSetMover name="atomic" cst_file="constraints_helix.cst" />
        FavorNativeResidue name="favor_native" bonus="0.00" />
        <Transform name="transform" chain="X" box_size="10.0" move_distance="0.1" angle="5" cycles="500"
repeats="1" temperature="5" rmsd="4.0" use_constraints="true" cst_fa_file="constraints_helix.cst"
cst_fa_weight="5" />
        HighResDocker name="high_res_docker" cycles="6" repack_every_Nth="3" scorefxn="ligand_soft_rep"
movemap_builder="docking" />
        <PackRotamersMover name="design_interface" scorefxn="hard_rep" task_operations="design_interface" />
        FinalMinimizer name="final" scorefxn="hard_rep" movemap_builder="final" />
        <InterfaceScoreCalculator name="add_scores" chains="X" scorefxn="hard_rep" />
        <ParsedProtocol name="low_res_dock">
            <Add mover_name="transform" />
        </ParsedProtocol>
        ParsedProtocol name="high_res_dock">
            Add mover_name="high_res_docker" />
            Add mover_name="final" />
        /ParsedProtocol>
        <PackRotamersMover name="fixbb_aro" scorefxn="ref15_1" task_operations="resfile,extrachi,ifcl"/>
        <PackRotamersMover name="pack" scorefxn="ref15_1" task_operations="ifcl,resfile,extrachi"/>
        <PackRotamersMover name="pack_fast" scorefxn="ref15_1" task_operations="ifcl,resfile"/>
        <MinMover name="min_bb" scorefxn="ref15" tolerance="0.0000001" max_iter="1000" chi="false"
bb="true">
            <MoveMap name="map_bb">
                <Span begin="1" end="200" bb="true" chi="false" />
                <Span begin="201" end="999" bb="false" chi="false"/>
            </MoveMap>
        </MinMover>
        <Idealize name="idealize"/>
        <MinMover name="min_sc" scorefxn="ref15" tolerance="0.0000001" max_iter="1000" chi="true"
bb="false">
            <MoveMap name="map_sc">
                <Span begin="1" end="200" bb="false" chi="true" />
                <Span begin="201" end="999" bb="false" chi="false"/>
            </MoveMap>
        </MinMover>
        <MinMover name="min_sc_bb" scorefxn="ref15" tolerance="0.0000001" max_iter="1000" chi="true"
bb="true">
            <MoveMap name="map_sc_bb">
                <Span begin="1" end="200" bb="true" chi="true" />
                <Span begin="201" end="999" bb="false" chi="false"/>
            </MoveMap>
        </MinMover>
        <ParsedProtocol name="parsed_pack_fast" >
            <Add mover_name="pack_fast"/>
            <Add mover_name="min_bb"/>
        </ParsedProtocol>
        <ParsedProtocol name="parsed_pack" >
            <Add mover_name="pack"/>
```

```
            <Add mover_name="min_bb"/>
            <Add mover_name="min_sc"/>
        </ParsedProtocol>
        <GenericMonteCarlo name="pack_mc" preapply="0" trials="3" temperature="0.03" filter_name="pstat_mc"
sample_type="high" mover_name="parsed_pack">
                <Filters>
                    <AND filter_name="total_score_1" temperature="15" sample_type="low"/>
                </Filters>
        </GenericMonteCarlo>
        <GenericMonteCarlo name="pack_fast_mc" preapply="0" trials="2" temperature="0.03"
filter_name="pstat_mc" sample_type="high" mover_name="parsed_pack_fast">
                <Filters>
                    <AND filter_name="total_score_1" temperature="15" sample_type="low"/>
                </Filters>
        </GenericMonteCarlo>
    </MOVERS>
    <PROTOCOLS>
        <Add mover="atomic"/>
        Add mover_name="favor_native" />
        <Add mover_name="fixbb_aro"/>
        <Add mover_name="low_res_dock" />
        <Add mover_name="design_interface" />
        Add mover_name="high_res_dock" />
        <Add mover_name="parsed_pack_fast"/>
        <Add mover_name="pack_fast_mc"/>
        <Add mover_name="pack_mc"/>
        <Add mover_name="min_sc_bb"/>
        Add mover_name="low_res_dock" />
        Add mover_name="design_interface" />
        Add mover_name="high_res_dock" />
        <Add mover_name="add_scores" />
        <Add filter_name="pstat"/>
        <Add filter_name="net_charge"/>
        <Add filter_name="ligen"/>
        <Add filter_name="total_score"/>
        <Add filter_name="bu"/>
    </PROTOCOLS>
    <OUTPUT scorefxn="ref15_1"/>
</ROSETTASCRIPTS>
```

#### Computational design of FABLE_Flag(.txt)

```
-out:path:pdb output
-out:path:score output
-extra_res_fa cou485.params
-packing:multi_cool_annealer 10
-packing:linmem_ig 10
-ignore_zero_occupancy false
-s able128_cou485.pdb
```

#### Computational design of FABLE_Command line inputs

```
~/rosetta_bin_linux_2020.08.61146_bundle/main/source/bin/rosetta_scripts.static.linuxgccrelease -database
/wynton/home/degradolab/lonelu/rosetta_bin_linux_2020.08.61146_bundle/main/database/ -parser:protocol
rscript_dock_fixbb_design_round1.xml @options_helix_1.txt -run:constant_seed -run:jran $SGE_TASK_ID -
out:suffix _$SGE_TASK_ID -out:no_nstruct_label
```

#### Computational design of FABLE_constraints (.cst)

```
AtomPair O2 1X ND1 49A HARMONIC 3.0 0.2
AtomPair C8 1X ND1 49A HARMONIC 3.7 0.2
Angle ND1 49A O2 1X C10 1X HARMONIC 1.91 0.12
Dihedral ND1 49A O2 1X C10 1X O1 1X CIRCULARHARMONIC 3.14 0.17
```

#### Computational design of FABLE_net charge (.charge)

```
DESIRED_CHARGE -5 #Desired net charge is zero.
PENALTIES_CHARGE_RANGE -10 -1 #Penalties are listed in the observed net charge range of -2 to +4.
PENALTIES 10 0 0 0 0 0 0 0 0 10 #The penalties are 10 for an observed charge of -2, 0 for an observed charge of -1
to +3, and 10 for an observed charge of +4.
BEFORE_FUNCTION QUADRATIC #Ramp quadratically for observed net charges of -3 or less.
AFTER_FUNCTION QUADRATIC #Ramp quadratically for observed net charges of +5 or greater.
```

**Computational design of FABLE_resfile (.txt)**

```
NATRO
ALLAAxc
NOTAA MTC
USE_INPUT_SC
start

1 A PIKAA S
2 A PIKAA TGIFYALSWVHM
3 A PIKAA K
4 A PIKAA S
5 A PIKAA E
6 A PIKAA TGIFYALSWVHM
7 A PIKAA A
8 A PIKAA E
9 A PIKAA TGIFYALSWVMH
10 A PIKAA KVSNLFRTDYAQWEIGHM
11 A PIKAA A
12 A PIKAA V
13 A PIKAA TGIFYALSWVH
14 A PIKAA KVSNLFRTDYAQWEIGHM
15 A PIKAA E
16 A PIKAA TGIFYALSWVNQH
17 A PIKAA KVSNLFRTDYAQWEIGHM
18 A PIKAA A
19 A PIKAA V
20 A PIKAA TGIFYALSWVHM
21 A PIKAA KVSNLFRTDYAQWEIGHM
22 A PIKAA T
23 A PIKAA TGIFYALSWVMH
24 A PIKAA KVSNLFRTDYAQWEIGHM
25 A PIKAA A
26 A PIKAA A
27 A PIKAA TGIFYALSWVHM
28 A PIKAA Q
29 A PIKAA N
30 A PIKAA G
31 A PIKAA D
32 A PIKAA K
33 A PIKAA E
34 A PIKAA A
35 A PIKAA TGIFYALSWVHM
36 A PIKAA A
37 A PIKAA Q
38 A PIKAA Y
39 A PIKAA TGIFYALSWVMH
40 A PIKAA A
41 A PIKAA R
42 A PIKAA TGIFYALSWVMNQH
43 A PIKAA A
44 A PIKAA S
45 A PIKAA L
46 A NATRO
47 A PIKAA T
48 A PIKAA R
49 A NATRO
```

50 A PIKAA E
51 A PIKAA E
52 A PIKAA L
53 A PIKAA TGIFYALSWVHM
54 A PIKAA N
55 A PIKAA R
56 A PIKAA TGIFYALSWVHM
57 A PIKAA L
58 A PIKAA Q
59 A PIKAA K
60 A PIKAA TGIFYALSWVHM
61 A PIKAA R
62 A PIKAA R
63 A PIKAA E
64 A PIKAA G
65 A PIKAA N
66 A PIKAA K
67 A PIKAA E
68 A PIKAA TGIFYALSWVHM
69 A PIKAA V
70 A PIKAA T
71 A PIKAA L
72 A PIKAA TGIFYALSWVHM
73 A PIKAA N
74 A PIKAA E
75 A PIKAA TGIFYALSWVHM
76 A PIKAA T
77 A PIKAA A
78 A PIKAA KVSNLFRTDYAQWEIGHM
79 A PIKAA TGIFYALSWVHM
80 A PIKAA Q
81 A PIKAA T
82 A PIKAA G
83 A PIKAA K
84 A PIKAA S
85 A PIKAA KVSNLFRTDYAQWEIGHM
86 A PIKAA TGIFYALSWVNQHM
87 A PIKAA N
88 A PIKAA A
89 A PIKAA TGIFYALSWVMH
90 A PIKAA KVSNLFRTDYAQWEIGHM
91 A PIKAA A
92 A PIKAA A
93 A PIKAA TGIFYALSWVMH
94 A PIKAA K
95 A PIKAA N
96 A PIKAA G
97 A PIKAA D
98 A PIKAA D
99 A PIKAA D
100 A PIKAA S
101 A PIKAA TGIFYALSWVMH
102 A PIKAA E
103 A PIKAA S
104 A PIKAA KVSNLFRTDYAQWEIGHM
105 A PIKAA VSNLFTYAQWIGHM

106 A PIKAA Q
107 A PIKAA A
108 A PIKAA TGIFYALSWVNQHM
109 A PIKAA KVSNLFRTDYAQWEIGHM
110 A PIKAA K
111 A PIKAA KVSNLFRTDYAQWEIGHM
112 A PIKAA VSNLFTYAQWIGHM
113 A PIKAA A
114 A PIKAA K
115 A PIKAA KVSNLFRTDYAQWEIGHM
116 A PIKAA KVSNLFRTDYAQWEIGHM
117 A PIKAA T
118 A PIKAA KVSNLFRTDYAQWEIGHM
119 A PIKAA TGIFYALSWVHM
120 A PIKAA KVSNLFRTDYAQWEIGHM
121 A PIKAA Q
122 A PIKAA TGIFYALSWVNQHM
123 A PIKAA A
124 A PIKAA K
125 A PIKAA A
126 A PIKAA VSNLFTYAQWIGHM

**Computational design of FABLE_cou485 parameters (.params)**

```
NAME cou
IO_STRING cou Z
TYPE LIGAND
AA UNK
ATOM  C2  aroC  X   -0.02
ATOM  O1  Oaro  X   -0.57
ATOM  C10 COO   X   0.71
ATOM  O2  OOC   X   -0.67
ATOM  C8  aroC  X   -0.02
ATOM  C4  aroC  X   -0.02
ATOM  C1  aroC  X   -0.02
ATOM  C6  aroC  X   -0.02
ATOM  C7  aroC  X   -0.02
ATOM  C3  aroC  X   -0.02
ATOM  N1  Nhis  X   -0.44
ATOM  C11 CH3   X   -0.18
ATOM  H5  Hapo  X   0.19
ATOM  H6  Hapo  X   0.19
ATOM  H7  Hapo  X   0.19
ATOM  C12 CH3   X   -0.18
ATOM  H8  Hapo  X   0.19
ATOM  H9  Hapo  X   0.19
ATOM  H10 Hapo  X   0.19
ATOM  C5  aroC  X   -0.02
ATOM  H1  Haro  X   0.21
ATOM  H3  Haro  X   0.21
ATOM  H2  Haro  X   0.21
ATOM  C9  CH1   X   0.00
ATOM  F1  F     X   -0.16
ATOM  F2  F     X   -0.16
ATOM  F3  F     X   -0.16
ATOM  H4  Haro  X   0.21
BOND_TYPE  F1   C9  1
BOND_TYPE  F2   C9  1
BOND_TYPE  F3   C9  1
BOND_TYPE  O1   C2  1
BOND_TYPE  O1   C10 1
BOND_TYPE  O2   C10 2
BOND_TYPE  N1   C3  1
BOND_TYPE  N1   C11 1
BOND_TYPE  N1   C12 1
BOND_TYPE  C1   C2  2
BOND_TYPE  C1   C4  1
BOND_TYPE  C1   C6  1
BOND_TYPE  C2   C5  1
BOND_TYPE  C3   C5  2
BOND_TYPE  C3   C7  1
BOND_TYPE  C4   C8  2
BOND_TYPE  C4   C9  1
BOND_TYPE  C5   H1  1
BOND_TYPE  C6   C7  2
BOND_TYPE  C6   H2  1
BOND_TYPE  C7   H3  1
BOND_TYPE  C8   C10 1
BOND_TYPE  C8   H4  1
```

```
BOND_TYPE  C11  H5  1
BOND_TYPE  C11  H6  1
BOND_TYPE  C11  H7  1
BOND_TYPE  C12  H8  1
BOND_TYPE  C12  H9  1
BOND_TYPE  C12  H10 1
CHI 1  C7   C3   N1   C11
CHI 2  C8   C4   C9   F1
NBR_ATOM  C2
NBR_RADIUS 5.923196
ICOOR_INTERNAL    C2      0.000000    0.000000    0.000000   C2    O1    C10
ICOOR_INTERNAL    O1      0.000000  180.000000    1.409420   C2    O1    C10
ICOOR_INTERNAL    C10     0.000000   58.463932    1.386066   O1    C2    C10
ICOOR_INTERNAL    O2    179.925876   57.937170    1.226078   C10   O1    C2
ICOOR_INTERNAL    C8   -179.925892   63.301065    1.472680   C10   O1    O2
ICOOR_INTERNAL    C4     -0.015728   56.938781    1.340468   C8    C10   O1
ICOOR_INTERNAL    C1      0.014038   61.035345    1.469219   C4    C8    C10
ICOOR_INTERNAL    C6   -179.980373   56.250303    1.401339   C1    C4    C8
ICOOR_INTERNAL    C7    179.987003   59.272543    1.399558   C6    C1    C4
ICOOR_INTERNAL    C3     -0.002786   59.626591    1.393812   C7    C6    C1
ICOOR_INTERNAL    N1   -179.998592   59.914843    1.409634   C3    C7    C6
ICOOR_INTERNAL    C11   -29.995192   59.285655    1.444938   N1    C3    C7
ICOOR_INTERNAL    H5   -160.400020   68.664840    1.095868   C11   N1    C3
ICOOR_INTERNAL    H6    118.861740   68.474544    1.096109   C11   N1    H5
ICOOR_INTERNAL    H7    121.608625   69.215932    1.095223   C11   N1    H6
ICOOR_INTERNAL    C12   179.691253   59.261398    1.444710   N1    C3    C11
ICOOR_INTERNAL    H8     79.879643   69.218588    1.095174   C12   N1    C3
ICOOR_INTERNAL    H9    119.524176   68.641682    1.095983   C12   N1    H8
ICOOR_INTERNAL    H10   118.889069   68.492185    1.095932   C12   N1    H9
ICOOR_INTERNAL    C5    179.996131   60.228042    1.390873   C3    C7    N1
ICOOR_INTERNAL    H1    179.316588   58.204851    1.086138   C5    C3    C7
ICOOR_INTERNAL    H3   -179.255672   61.873585    1.087037   C7    C6    C3
ICOOR_INTERNAL    H2   -179.746324   51.829077    1.075641   C6    C1    C7
ICOOR_INTERNAL    C9    179.986364   58.867821    1.500828   C4    C8    C1
ICOOR_INTERNAL    F1     59.996833   68.979155    1.347636   C9    C4    C8
ICOOR_INTERNAL    F2    120.009390   69.167204    1.347464   C9    C4    F1
ICOOR_INTERNAL    F3    119.943271   69.078969    1.347408   C9    C4    F2
ICOOR_INTERNAL    H4    179.999796   63.686307    1.085843   C8    C10   C4
```

**Computational design of FABLE_composition (.comp)**

```
TYPE THR
DELTA_START 0
DELTA_END 1
PENALTIES 0 100
ABSOLUTE 5
BEFORE_FUNCTION CONSTANT
AFTER_FUNCTION QUADRATIC
END_PENALTY_DEFINITION

PENALTY_DEFINITION
TYPE GLY
DELTA_START 0
DELTA_END 1
PENALTIES 0 100
ABSOLUTE 9
BEFORE_FUNCTION CONSTANT
AFTER_FUNCTION QUADRATIC
END_PENALTY_DEFINITION

PENALTY_DEFINITION
TYPE SER
DELTA_START 0
DELTA_END 1
PENALTIES 0 100
ABSOLUTE 5
BEFORE_FUNCTION CONSTANT
AFTER_FUNCTION QUADRATIC
END_PENALTY_DEFINITION

PENALTY_DEFINITION
TYPE ASN
DELTA_START 0
DELTA_END 1
PENALTIES 0 100
ABSOLUTE 8
BEFORE_FUNCTION CONSTANT
AFTER_FUNCTION QUADRATIC
END_PENALTY_DEFINITION

PENALTY_DEFINITION
TYPE ARG
DELTA_START 0
DELTA_END 1
PENALTIES 0 100
ABSOLUTE 8
BEFORE_FUNCTION CONSTANT
AFTER_FUNCTION QUADRATIC
END_PENALTY_DEFINITION

PENALTY_DEFINITION
TYPE ALA
DELTA_START 0
DELTA_END 1
PENALTIES 0 100
ABSOLUTE 18
```

BEFORE_FUNCTION CONSTANT
AFTER_FUNCTION QUADRATIC
END_PENALTY_DEFINITION

PENALTY_DEFINITION
TYPE TRP
DELTA_START 0
DELTA_END 1
PENALTIES 0 100
ABSOLUTE 3
BEFORE_FUNCTION CONSTANT
AFTER_FUNCTION QUADRATIC
END_PENALTY_DEFINITION

PENALTY_DEFINITION
TYPE LYS
DELTA_START 0
DELTA_END 1
PENALTIES 0 100
ABSOLUTE 8
BEFORE_FUNCTION CONSTANT
AFTER_FUNCTION QUADRATIC
END_PENALTY_DEFINITION

PENALTY_DEFINITION
TYPE GLN
DELTA_START 0
DELTA_END 1
PENALTIES 0 100
ABSOLUTE 8
BEFORE_FUNCTION CONSTANT
AFTER_FUNCTION QUADRATIC
END_PENALTY_DEFINITION

PENALTY_DEFINITION
TYPE TYR
DELTA_START 0
DELTA_END 1
PENALTIES 0 100
ABSOLUTE 8
BEFORE_FUNCTION CONSTANT
AFTER_FUNCTION QUADRATIC
END_PENALTY_DEFINITION

PENALTY_DEFINITION
TYPE GLU
DELTA_START 0
DELTA_END 1
PENALTIES 0 100
ABSOLUTE 8
BEFORE_FUNCTION CONSTANT
AFTER_FUNCTION QUADRATIC
END_PENALTY_DEFINITION

PENALTY_DEFINITION
TYPE ASP

```
DELTA_START 0
DELTA_END 1
PENALTIES 0 100
ABSOLUTE 9
BEFORE_FUNCTION CONSTANT
AFTER_FUNCTION QUADRATIC
END_PENALTY_DEFINITION

PENALTY_DEFINITION
TYPE VAL
DELTA_START 0
DELTA_END 1
PENALTIES 0 100
ABSOLUTE 12
BEFORE_FUNCTION CONSTANT
AFTER_FUNCTION QUADRATIC
END_PENALTY_DEFINITION

PENALTY_DEFINITION
TYPE MET
DELTA_START 0
DELTA_END 1
PENALTIES 0 100
ABSOLUTE 8
BEFORE_FUNCTION CONSTANT
AFTER_FUNCTION QUADRATIC
END_PENALTY_DEFINITION
```

#### KABLE_Design_Strategy_1_ Constraints

AtomPair N3 1X OE2 108A HARMONIC 2.5 0.1
Dihedral N1 1X N2 1X N3 1X OE2 108A CIRCULARHARMONIC 3.14 0.17
Angle OE2 108A N3 1X N2 1X HARMONIC 1.92 0.17
Angle CD 108A OE2 108A N3 1X HARMONIC 1.92 0.17

#### KABLE_Design_Strategy_2_Constraints

AtomPair N3 1X OD2 49A HARMONIC 2.5 0.1
Dihedral N1 1X N2 1X N3 1X OD2 49A CIRCULARHARMONIC 3.14 0.17
Dihedral CB 49A CG 49A OD2 49A N3 1X CIRCULARHARMONIC 3.14 0.17
Angle OD2 49A N3 1X N2 1X HARMONIC 1.92 0.17
Angle CG 49A OD2 49A N3 1X HARMONIC 1.92 0.17

#### Example of KABLE_sequence_design_LigandMPNN

```
/home/degradolab/yuda/LigandMPNN/run.py \
        --model_type "ligand_mpnn" \
        --seed 111 \
        --pdb_path "… " \
        --out_folder "…" \
        --fixed_residues "A49" \
        --omit_AA "C" \
        --batch_size 2 \
        --number_of_batches 5 \
```

#### Example of KABLE_sequence_design_Rosetta FastRelax

```
~/rosetta_bin_linux_2020.08.61146_bundle/main/source/bin/relax.static.linuxgccrelease \
    -database ~ /rosetta_bin_linux_2020.08.61146_bundle/main/database/ \
    -in:file:s $pdb_filename \
    -extra_res_fa ~/…/ 6nt.params \
    -constraints:cst_fa_file ~/…/constraints.cst \
        -constraints:cst_fa_weight "5" \
    -relax:quick \
```

**Extract_Chi1_and_Chi2.py**

```
import os
import csv
from Bio.PDB import PDBParser
from Bio.PDB.vectors import calc_dihedral
import math

# Define a mapping of residue types to the atoms involved in chi1 and chi2 calculations
residue_atoms = {
    'default': [('N', 'CA', 'CB', 'CG'), ('CA', 'CB', 'CG', 'CD')],  # Default for most amino acids
    'TYR': [('N', 'CA', 'CB', 'CG'), ('CA', 'CB', 'CG', 'CD1')],  # Tyrosine
    'HIS': [('N', 'CA', 'CB', 'CG'), ('CA', 'CB', 'CG', 'ND1')],  # Histidine
    'ASP': [('N', 'CA', 'CB', 'CG'), ('CA', 'CB', 'CG', 'OD1')],  # Aspartic acid
    'GLN': [('N', 'CA', 'CB', 'CG'), ('CA', 'CB', 'CG', 'CD')],   # Glutamine
    'LEU': [('N', 'CA', 'CB', 'CG'), ('CA', 'CB', 'CG', 'CD1')],  # Leucine
    'ILE': [('N', 'CA', 'CB', 'CG1'), ('CA', 'CB', 'CG1', 'CD1')],  # Isoleucine
    'PHE': [('N', 'CA', 'CB', 'CG'), ('CA', 'CB', 'CG', 'CD1')],  # Phenylalanine
    'THR': [('N', 'CA', 'CB', 'OG1')],  # Threonine (chi1 only)
    # Add more residue types as needed
}

def calculate_dihedrals(pdb_file, residue_number, residue_type='default'):
    parser = PDBParser()
    structure = parser.get_structure('X', pdb_file)
    atom_pairs = residue_atoms.get(residue_type, residue_atoms['default'])
    chi1_atoms = atom_pairs[0]  # Chi1 is always the first set of atoms
    chi2_atoms = atom_pairs[1] if len(atom_pairs) > 1 else None  # Chi2 is optional
    for model in structure:
        for chain in model:
            for residue in chain:
                if residue.id[1] == residue_number:
                    try:
                        # Extract vectors for chi1 calculation
                        vectors_chi1 = [residue[x].get_vector() for x in chi1_atoms]
                        chi1 = calc_dihedral(*vectors_chi1)
                        chi1_deg = math.degrees(chi1)

                        # Extract vectors for chi2 calculation if available
                        chi2_deg = None
                        if chi2_atoms:
                            vectors_chi2 = [residue[x].get_vector() for x in chi2_atoms]
                            chi2 = calc_dihedral(*vectors_chi2)
                            chi2_deg = math.degrees(chi2)
                        return chi1_deg, chi2_deg
                    except KeyError:
                        # Atoms required not found
                        return None, None
    return None, None  # Residue not found

def process_directory(directory_path, residue_number, residue_type, output_csv):
    with open(output_csv, 'w', newline='') as csvfile:
        fieldnames = ['File Name', 'Chi1 (degrees)', 'Chi2 (degrees)']
        writer = csv.DictWriter(csvfile, fieldnames=fieldnames)
        writer.writeheader()

        for file in os.listdir(directory_path):
```

```
            if file.endswith(".pdb"):
                pdb_file_path = os.path.join(directory_path, file)
                chi1, chi2 = calculate_dihedrals(pdb_file_path, residue_number, residue_type)
                writer.writerow({'File Name': file, 'Chi1 (degrees)': chi1, 'Chi2 (degrees)': chi2 if chi2 is not None else
"N/A"})

# Example usage
directory_path =
'/Users/chenyuda/Documents/Code/ABLE_Fragment/fragment_paper/md_278k_hie_202404/chi1chi2_for_H49/all_
pdb_for_H49/'
residue_number = 49  # Example residue number
residue_type = 'HIS'  # Specify residue type here
output_csv = 'Chi1Chi2_md_Thr112_6w70_ABLE_6w6x_6w70_and_pdb.csv'
process_directory(directory_path, residue_number, residue_type, output_csv)
```

**Generate_contour_from_tyr_chi1and2.py**

```
import numpy as np
import pandas as pd
import matplotlib.pyplot as plt
from scipy.stats import gaussian_kde
from matplotlib.colors import LinearSegmentedColormap

def plot_density_contour(file_path, labels, x_range, y_range, contour_color, scatter_colors, contour_line_color,
bg_color='white', contour_alpha=0.6, contour_linewidth=3, symbol_sizes=None, output_file='density_contour.svg'):
    fig, ax = plt.subplots(figsize=(10, 8), facecolor=bg_color)
    ax.set_facecolor(bg_color)

    # Read the CSV file
    data_frame = pd.read_csv(file_path, header=None)

    # Extract group names
    group_names = data_frame.iloc[0, ::2].values

    # Process contour group (first group)
    chi1_contour = data_frame.iloc[1:, 0].dropna().values.astype(float)
    chi2_contour = data_frame.iloc[1:, 1].dropna().values.astype(float)
    data_contour = np.vstack([chi1_contour, chi2_contour])
    kde_contour = gaussian_kde(data_contour)
    x_min, x_max = x_range
    y_min, y_max = y_range
    x_grid, y_grid = np.mgrid[x_min:x_max:100j, y_min:y_max:100j]
    grid_coords = np.vstack([x_grid.ravel(), y_grid.ravel()])
    density_contour = kde_contour(grid_coords).reshape(x_grid.shape)
    cmap = LinearSegmentedColormap.from_list(f'custom_cmap_{labels[0]}', ['white', contour_color])
    contour_filled = ax.contourf(x_grid, y_grid, density_contour, levels=4, cmap=cmap, alpha=contour_alpha,
extend='both')
    contour_lines = ax.contour(x_grid, y_grid, density_contour, levels=4, colors=contour_line_color,
linewidths=contour_linewidth)
    ax.scatter(chi1_contour, chi2_contour, c=contour_color, s=symbol_sizes[0], label=labels[0])
    print(f"{labels[0]}: {len(chi1_contour)} points")

    # Process scatter groups (all other groups)
    for i in range(1, len(labels)):
        chi1_scatter = data_frame.iloc[1:, 2*i].dropna().values.astype(float)
        chi2_scatter = data_frame.iloc[1:, 2*i+1].dropna().values.astype(float)

        if len(chi1_scatter) == 0 or len(chi2_scatter) == 0:
            print(f"{labels[i]}: No data found")
            continue

        ax.scatter(chi1_scatter, chi2_scatter, c=scatter_colors[i-1], s=symbol_sizes[i], label=labels[i])
        print(f"{labels[i]}: {len(chi1_scatter)} points")

    # Add color bar for the filled contour
    plt.colorbar(contour_filled, ax=ax, label='Density')

    # Customizing the range for the x and y axes
    ax.set_xlim([x_min, x_max])
    ax.set_ylim([y_min, y_max])

    ax.set_xlabel('chi1')
```

```
    ax.set_ylabel('chi2')
    ax.set_title('Density Contour Plot of chi1 and chi2')
    ax.legend()

    # Save the plot as an SVG file
    plt.savefig(output_file, format='svg')
    print(f"Plot saved as {output_file}")

    plt.show()

# File path to the CSV file
file_path = '…' # Replace with your actual file path

# Labels for the datasets
labels = ['complexed ABLE MD Simulation', 'Uncomplexed ABLE', 'Uncomplexed ABLE H49A', 'ABLE-apixaban complex']

# Colors for the contour plot
contour_color = '#F39191'

# Colors for the scatter plots
scatter_colors = ['#1873E5', '#59B358', '#5E4B56']

# Color for the contour lines
contour_line_color = '#ffc8c8'

# Custom ranges for x and y axes
x_range = [225, 340]
y_range = [-125, 50]

# Symbol sizes for the datasets (contour and five scatter plots)
symbol_sizes = [20, 80, 80, 80]

# Call the function with desired customizations and specify output file
output_file = '…'
plot_density_contour(file_path, labels, x_range, y_range, contour_color, scatter_colors, contour_line_color,
bg_color='white', contour_alpha=0.8, contour_linewidth=0.1, symbol_sizes=symbol_sizes, output_file=output_file)
```

#### Count_water.py.

```
import prody as pr
import matplotlib.pyplot as plt
import csv
#import warnings
#warnings.filterwarnings('ignore')

####### Script settings ######
discard_frames = […]

lig_dist = 3.5

traj_pdb_path = '…'

lig_chain = '…'

lig_resnum = 127

output_csv_file = '…'

######## Functions #######

def main():
    # Load MD trajectory
    traj_pdb = pr.parsePDB(traj_pdb_path)
    print('-'*20, 'Total number of frames in traj: ', len(traj_pdb.getCoordsets()), '-'*20)

    # Perform analysis
    num_waters_per_frame = [] # Each element in this list represents a single frame.
    num_frames_processed = 0
    # Iterate over each frame (model), which is a Coordset in prody
    for frame_ix, coordset in enumerate(traj_pdb.getCoordsets()):
        if frame_ix in discard_frames:
            continue
        num_frames_processed += 1
        # Make a standalone prody obj for each frame
        par = pr.atomic.atomgroup.AtomGroup()
        par.setCoords(coordset)
        par.setNames(traj_pdb.getNames())
        par.setResnames(traj_pdb.getResnames())
        par.setChids(traj_pdb.getChids())
        par.setResnums(traj_pdb.getResnums())
        par.setElements(traj_pdb.getElements())

        # Define ligand.
        if lig_chain:
            pr_lig = par.select(f'chain {lig_chain} and resnum {lig_resnum}')
        else:
            pr_lig = par.select(f'resnum {lig_resnum}')

        lig_resname = set(pr_lig.getResnames())
        lig_heavyatoms = pr_lig.select('not element H')
        if len(lig_resname) > 1:
            print(f'Note: more than one ligand selected. Residue names {lig_resname}.')
```

```
        if len(num_waters_per_frame) == 0:
            print(f'Ligand {lig_resname} selected with {len(lig_heavyatoms)} heavy atoms.')

        # Count waters in the active site, defined by a distance cutoff from the ligand.
        wat_neighbs = par.select(
                f'(water and name O) within {lig_dist} of Lig', Lig=pr_lig)
        if wat_neighbs is None:
            num_waters_in_frame = 0
        else:
            num_waters_in_frame = len(wat_neighbs)

        # Record the frame number (1-indexed) and the number of waters in that frame.
        num_waters_per_frame.append(tuple([frame_ix+1, num_waters_in_frame]))

    framenumbers = [i[0] for i in num_waters_per_frame]
    numberwaters = [i[1] for i in num_waters_per_frame]

    # Write out the results to a csv file.
    with open(output_csv_file, 'w', newline='') as csvfile:
        writer = csv.writer(csvfile)
        writer.writerow(['Frame #', '# waters'])
        for frame, waters in zip(framenumbers, numberwaters):
            writer.writerow([frame, waters])

    # Plot
    plt.plot(framenumbers, numberwaters)
    plt.xlabel('Frame #')
    plt.ylabel('# waters in active site')
    plt.title(traj_pdb_path)
    plt.grid(True)
    plt.show()

    print(f'Number of frames processed: {num_frames_processed}')
    print(f'Number of frames discarded: {len(discard_frames)}')



######## Main #######
if __name__ == "__main__":
    main()
```

**Generate_landscape.py.**

```
import numpy as np
import plotly.graph_objects as go
import plotly.io as pio

# Force the renderer to "browser" so that the full mode bar is visible.
pio.renderers.default = "browser"

# ----------------------------
# 1. Parameters for Each Peak (Anisotropic)
# ----------------------------
# For each peak, parameters are now: (amplitude, sigma_x, sigma_y)

# ABLE: (0.25, 2.74, 2.74)
ABLE_shift_x = -10     # Because "position = (10, 50)" => shift of -10
ABLE_shift_y = -50     # position = 50 => shift of -50
ABLE_params  = (0.25, 2.74, 2.74)

# KABLE: (0.25, 2.236, 2.236)
KABLE_shift_x = -75
KABLE_shift_y = -80
KABLE_params  = (0.25, 2.236, 2.236)

# ABLE "promiscuous" peak: (0.04, 8.66, 8.66)
PROM_shift_x = -35
PROM_shift_y = -50
PROM_params  = (0.04, 8.66, 8.66)

# FABLE: (0.25, 2.74, 2.74)
FABLE_shift_x = -90
FABLE_shift_y = -20
FABLE_params  = (0.25, 2.74, 2.74)

# Transition (fifth peak): (0.0175, 1, 1)
TRANS_shift_x = -20
TRANS_shift_y = -50
TRANS_params  = (0.0175, 1, 1)

# ----------------------------
# 2. Gaussian & Surface Setup
# ----------------------------
noise_level = 0.000

def generate_grid(x_start, x_end, y_start, y_end, num_points):
    """Generate a meshgrid of x and y."""
    x = np.linspace(x_start, x_end, num_points)
    y = np.linspace(y_start, y_end, num_points)
    X, Y = np.meshgrid(x, y)
    return X, Y

def calculate_gaussian_z(x, y, shift_x, shift_y, params):
    """
    Calculate an anisotropic Gaussian.

    params = (a, sigma_x, sigma_y)
      a: amplitude
```

```
    sigma_x: width in x-direction
    sigma_y: width in y-direction
    """
    a, sigma_x, sigma_y = params
    z = a * np.exp(-((x + shift_x)**2 / (2 * sigma_x**2) + (y + shift_y)**2 / (2 * sigma_y**2)))
    return z

def add_noise(z, noise_level):
    """Add random noise to z values (if desired)."""
    np.random.seed(0)
    noise = noise_level * np.random.normal(size=z.shape)
    return z + noise

def calculate_combined_z(x, y, noise_level):
    """
    Sum five anisotropic Gaussians: ABLE, KABLE, promiscuous, FABLE, and transition.
    """
    z_able = calculate_gaussian_z(x, y, ABLE_shift_x, ABLE_shift_y, ABLE_params)
    z_kabl = calculate_gaussian_z(x, y, KABLE_shift_x, KABLE_shift_y, KABLE_params)
    z_prom = calculate_gaussian_z(x, y, PROM_shift_x, PROM_shift_y, PROM_params)
    z_fabl = calculate_gaussian_z(x, y, FABLE_shift_x, FABLE_shift_y, FABLE_params)
    z_trans = calculate_gaussian_z(x, y, TRANS_shift_x, TRANS_shift_y, TRANS_params)

    z_sum = z_able + z_kabl + z_prom + z_fabl + z_trans
    return add_noise(z_sum, noise_level)

# ----------------------------
# 3. Plotting Function with Thick Grid Overlay and Modified Aspect Ratio
# ----------------------------
def plot_landscape_interactive(x, y, z):
    """
    Plots the 3D surface using a uniform grey color with an overlaid grid
    constructed from Scatter3d lines to simulate thicker grid lines.
    The axes and labels are removed, and the scene's aspect ratio is set so that
    the height (z-axis) is adjusted relative to x and y.
    """
    # Define a uniform grey colorscale for the surface.
    custom_color_scale = [
        [0, 'rgb(210,227,226)'],
        [1, 'rgb(210,227,226)']
    ]

    # Use lighting settings that minimize shading effects.
    lighting_settings = dict(
        ambient=1,
        diffuse=0,   # Turn off diffuse shading
        specular=0,  # Turn off specular highlights
        roughness=0,
        fresnel=0
    )

    # Create the surface plot without any unsupported properties.
    surface = go.Surface(
        x=x,
        y=y,
        z=z,
```

```
    colorscale=custom_color_scale,
    showscale=False,
    lighting=lighting_settings,
    contours={
        "x": {"show": False},
        "y": {"show": False},
        "z": {"show": False}
    }
)

# Define the highlight sets and colors.
highlight_x_values = {10, 75, 30,35, 40, 90}
highlight_y_values = {50, 80, 20}
highlight_color = 'rgb(82,128,136)'
default_color = 'rgb(151,179,183)'

# Build grid overlay as Scatter3d lines.
grid_lines = []
thick_line_width = 10  # Consistent line thickness

# Set grid lines with spacing of 5 units
x_values = np.arange(x.min(), x.max() + 1, 5)
y_values = np.arange(y.min(), y.max() + 1, 5)

# Add a small offset to grid lines to prevent Z-fighting.
z_offset = 0.001

# Overlay vertical grid lines (constant x, varying y)
for x_val in x_values:
    grid_color = highlight_color if x_val in highlight_x_values else default_color
    y_line = np.linspace(y.min(), y.max(), 100)
    z_line = calculate_combined_z(np.full_like(y_line, x_val), y_line, noise_level)
    z_line = 0.25 * z_line + z_offset  # Scale and add offset
    grid_lines.append(go.Scatter3d(
        x=[x_val] * len(y_line),
        y=y_line,
        z=z_line,
        mode='lines',
        line=dict(color=grid_color, width=thick_line_width),
        showlegend=False
    ))

# Overlay horizontal grid lines (constant y, varying x)
for y_val in y_values:
    grid_color = highlight_color if y_val in highlight_y_values else default_color
    x_line = np.linspace(x.min(), x.max(), 100)
    z_line = calculate_combined_z(x_line, np.full_like(x_line, y_val), noise_level)
    z_line = 0.25 * z_line + z_offset  # Use same scaling and offset
    grid_lines.append(go.Scatter3d(
        x=x_line,
        y=[y_val] * len(x_line),
        z=z_line,
        mode='lines',
        line=dict(color=grid_color, width=thick_line_width),
        showlegend=False
    ))
```

```
    # Create the figure with the surface and grid lines.
    fig = go.Figure(data=[surface] + grid_lines)

    # Set the scene to use orthographic projection and a square container for equal x and y dimensions.
    fig.update_layout(
        title="",
        autosize=False,
        width=1200,
        height=1200,
        scene=dict(
            xaxis=dict(visible=False),
            yaxis=dict(visible=False),
            zaxis=dict(visible=False),
            aspectmode='manual',
            aspectratio=dict(x=1, y=1, z=0.75),
            camera=dict(
                projection=dict(type='orthographic')
            )
        ),
        margin=dict(l=0, r=0, b=0, t=0)
    )

    # Enable the download button (toImage) in the mode bar with high-res image options.
    fig.show(config={
        'displayModeBar': True,
        'modeBarButtonsToAdd': ['toImage'],
        'toImageButtonOptions': {
            'format': 'png',
            'filename': 'landscape_high_res',
            'scale': 10
        }
    })

    # Optionally, you can automatically export an image:
    # fig.write_image("landscape_high_res.png", scale=10)

# ----------------------------
# 4. Generate Grid and Plot
# ----------------------------
if __name__ == "__main__":
    # Define grid boundaries and resolution.
    x_start, x_end, y_start, y_end = 0, 100, 0, 100
    num_points = 1000  # Increased resolution by 10× (from 100 to 1000)

    # Generate the grid.
    x, y = generate_grid(x_start, x_end, y_start, y_end, num_points)
    # Calculate the combined Gaussian surface.
    z = calculate_combined_z(x, y, noise_level)

    # Decrease the overall height of the landscape by scaling the z-values.
    z = 0.25 * z

    # Plot the interactive landscape with the thick grid overlay and modified aspect ratio.
     plot_landscape_interactive(x, y, z)
```
